# Sequential breakdown of the complex *Cf-9* leaf mould resistance locus in tomato by *Fulvia fulva*

**DOI:** 10.1101/2023.08.27.554972

**Authors:** Silvia de la Rosa, Christiaan R. Schol, Ángeles Ramos Peregrina, David J. Winter, Anne M. Hilgers, Kazuya Maeda, Yuichiro Iida, Mariana Tarallo, Ruifang Jia, Henriek G. Beenen, Mercedes Rocafort, Pierre J.G.M. de Wit, Joanna K. Bowen, Rosie E. Bradshaw, Matthieu H.A.J. Joosten, Yuling Bai, Carl H. Mesarich

**Affiliations:** Laboratory of Molecular Plant Pathology, School of Agriculture and Environment, Massey University, Palmerston North, New Zealand; Plant Breeding, Wageningen University & Research, Wageningen, The Netherlands; Laboratory of Phytopathology, Wageningen University & Research, Wageningen, The Netherlands; Bioinformatics Group, School of Natural Sciences, Massey University, Palmerston North, New Zealand; Laboratory of Plant Pathology, Faculty of Agriculture, Setsunan University, Hirakata, Osaka, Japan; Laboratory of Molecular Plant Pathology, School of Natural Sciences, Massey University, Palmerston North, New Zealand; The New Zealand Institute for Plant and Food Research Limited, Mount Albert Research Centre, Auckland, New Zealand; Bioprotection Aotearoa, Massey University, Palmerston North, New Zealand

**Keywords:** *Avr9* and *Avr9B* avirulence effector genes, *Cf-9C* and *Cf-9B* resistance genes, *Cf-9* locus, sequential resistance breakdown, tomato leaf mould disease, fungus, *Fulvia fulva* (*Cladosporium fulvum*), *Solanum lycopersicum*

## Abstract

- Leaf mould, caused by *Fulvia fulva*, is a devastating disease of tomato plants. In many commercial tomato cultivars, resistance to this disease is governed by the *Cf-9* locus, which comprises five paralogous genes (*Cf-9A–9E*) that encode receptor-like proteins. Two of these proteins contribute to resistance: Cf-9C recognizes the previously identified *F. fulva* effector Avr9 and provides resistance during all plant growth stages, while Cf-9B recognises the yet-unidentified *F. fulva* effector Avr9B and provides mature plant resistance only. In recent years, *F. fulva* strains have emerged that have overcome the *Cf-9* locus, with *Cf-9C* circumvented through *Avr9* deletion. To understand how *Cf-9B* is circumvented, we set out to identify *Avr9B*.
- Comparative genomics, *in planta* transient expression assays and gene complementation experiments were used to identify *Avr9B*, while gene sequencing was used to assess *Avr9B* allelic variation across a worldwide strain collection.
- A strict correlation between *Avr9* deletion and resistance-breaking mutations in *Avr9B* was observed in strains recently collected from *Cf-9* cultivars, whereas *Avr9* deletion but no mutations in *Avr9B* were observed in older strains.
- This research showcases how *F. fulva* has evolved to sequentially break down the two functional resistance genes of the complex *Cf-9* locus and highlights that this locus now has limited value for controlling leaf mould disease in worldwide commercial tomato production.

## Introduction

Leaf mould, caused by the hemibiotrophic fungus *Fulvia fulva* (formerly known as *Cladosporium fulvum* and *Passalora fulva*) (Videira *et al*., 2017), is a devastating disease of tomato plants grown in humid settings, such as in greenhouse and high tunnel environments, where it is responsible for severe defoliation and yield losses (Thomma *et al*., 2005). During infection, *F. fulva* resides in the apoplastic environment located between the mesophyll cells of leaves, where it secretes an arsenal of effector proteins to promote host colonization and lifecycle completion (de Wit, 2016; Mesarich *et al*., 2023; Rocafort *et al*., 2020). This arsenal includes at least 75 small, secreted proteins (SSPs) of <300 amino acid residues in length, most of which are stabilized with disulphide bonds that prevent their degradation by apoplastic tomato proteases (Joosten *et al*., 1997; Luderer *et al*., 2002a; Mesarich *et al*., 2018).

Resistance to *F. fulva* in tomato is governed by *Cf* (resistance to *Cladosporium fulvum*) immune receptor genes which, based on those cloned to date, encode cell surface-localized receptor-like proteins (RLPs) (Kang & Yeom 2018; Thomas *et al*., 1998). These RLPs possess an extracellular leucine-rich repeat (LRR) domain that is responsible for the direct or indirect recognition of specific *F. fulva* effectors (termed avirulence (Avr) effectors), as well as a transmembrane domain and a short cytoplasmic tail (Ngou *et al*., 2023; Snoeck *et al*., 2023). As RLPs do not have signalling capacity themselves, they typically need to interact with receptor-like kinases (RLKs), such as SOBIR1 and BAK1, which are involved in transducing defence response signals following apoplastic effector recognition (Gust & Felix, 2014; Liebrand *et al*., 2013; Liebrand *et al*., 2014; Postma *et al*., 2016; van der Burgh *et al*., 2019). Following recognition and signalling, immune responses such as a swift burst of reactive oxygen species (ROS) (Huang *et al*., 2020), callose deposition and the hypersensitive response (HR), which is a localized form of programmed cell death (de Wit *et al*., 2009), are initiated that halt *F. fulva* growth.

To date, five tomato *Cf* gene/*F. fulva Avr* effector gene pairs have been cloned. These are *Cf-2*/*Avr2* (Dixon *et al*., 1996; Luderer *et al*., 2002b), *Cf-4*/*Avr4* (Thomas *et al*., 1997; Joosten *et al*., 1994), *Cf-4E*/*Avr4E* (Takken *et al*., 1999; Thomas *et al*., 1997; Westerink *et al*., 2004), *Cf-5*/*Avr5* (Dixon *et al*., 1998; Mesarich *et al*., 2014), and *Cf-9C*/*Avr9* (Jones *et al*., 1994; van den Ackerveken *et al*., 1992; van Kan *et al*., 1991). In each of these cases, strains (races) of *F. fulva* have emerged that can overcome resistance mediated by the matching *Cf* gene. With regards to the corresponding *Avr* effector genes, circumvention has been achieved through gene deletion, indels (resulting in frame-shift mutations and truncations), transposable element insertions, or point mutations (leading to amino acid substitutions) (Joosten *et al*., 1994; Joosten *et al*., 1997; Luderer *et al*., 2002b; Mesarich *et al*., 2014; Westerink *et al*., 2004; van Kan *et al*., 1991).

Most commercial *F. fulva*-resistant tomato cultivars deployed worldwide are designated Ff:A-E, which indicates that they carry the *Cf-9* resistance locus from the wild tomato species *Solanum pimpinellifolium* (Jones *et al*., 1993; van der Beek *et al*., 1992). This locus is made up of five paralogous genes, namely *Cf-9A*, *Cf-9B*, *Cf-9C* (commonly referred to as *Cf-9* in the literature), *Cf-9D* and *Cf-9E* (Jones *et al*., 1994; Parniske *et al*., 1997). Of the RLPs that are encoded by these five genes, two provide resistance against *F. fulva*. More specifically, Cf-9C, as mentioned above, recognizes the Avr9 effector protein of *F. fulva* (van den Ackerveken *et al*., 1992; van Kan *et al*., 1991) and provides resistance during all stages of plant growth, while Cf-9B recognises the putative Avr9B effector and provides resistance to mature (flowering and fruiting) plants only (Jones *et al*., 1994; Laugé *et al*., 1998; Panter *et al*., 2002). The molecular mechanism governing this mature plant resistance is currently unknown, however developmental control of *Cf-9B* promoter activity does not appear to be responsible (Panter *et al*., 2002).

In addition to the timing of mounting actual resistance, differences also exist in the strength of the resistance response initiated by Cf-9C and Cf-9B. Indeed, Cf-9C is associated with an HR that generally restricts the hyphal growth of *F. fulva* to within one or two epidermal cell lengths of the penetration site (Hammond-Kosack and Jones, 1994; Laugé *et al*., 1998; Parniske *et al*., 1997), while Cf-9B is associated with leaf chlorosis and a strong accumulation of pathogenesis-related (PR) proteins that ultimately halt fungal growth after some hyphal extension between the mesophyll cells (Laugé *et al*., 1998; Panter *et al*., 2002; Parniske *et al*., 1997).

Perhaps unsurprisingly, given its extensive use in commercial tomato production worldwide, resistance provided by the *Cf-9* locus has already been partially overcome by many *F. fulva* strains. The first evidence of this circumvention was provided in the Netherlands, Poland, and France, where following the original introgression event of the *Cf-9* locus into commercial tomato cultivars during the 1970s (Schouten *et al*., 2019), race 9 (*Avr9*^−^) strains of *F. fulva* started to emerge that could overcome resistance mediated by the *Cf-9C* gene (Laterrot, 1986; Lindhout *et al*., 1989). In another example from Japan, race 9 strains were identified in 2009, just three years after commercial tomato cultivars carrying the *Cf-9* locus were introduced (Enya *et al*., 2009; Iida *et al*., 2010; Iida *et al*., 2015; Yoshida *et al*., 2021). More recently, a report stated that all 36 strains of *F. fulva* sampled in Cuba could overcome *Cf-9C*- mediated resistance (Bernal-Cabrera *et al*., 2021). In the cases where these studies have been supported or followed up with molecular diagnostics, circumvention of *Cf-9C*-mediated resistance has been shown to be exclusively the result of *Avr9* gene deletion (Bernal-Cabrera *et al*., 2021; Iida *et al*., 2015; Stergiopoulos *et al*., 2007a; Yoshida *et al*., 2021; van Kan *et al*., 1991).

Interestingly, even though *Cf-9C*-mediated resistance has been overcome by many strains of *F. fulva* globally, *Cf-9B*-mediated resistance has, until relatively recently, proven to be quite durable. However, over the last decade or so, growers from around the world have reported an increase in the incidence of *F. fulva* strains on mature *Cf-9* tomato plants. This is in addition to a study involving a New Zealand *F. fulva* strain, referred to as IPO 2679, collected in the 1980s, which was shown to have overcome both *Cf-9C-* and *Cf-9B*-mediated resistance (Laugé *et al*., 1998). With these points in mind, we set out to (i) identify the *Avr9B* gene, (ii) determine whether the *Avr9B* gene has been deleted or mutated across a worldwide collection of race 9 *F. fulva* strains isolated from mature *Cf-9* tomato plants, and (iii) ascertain whether *Cf-9C*- and *Cf-9B*-mediated resistance have been sequentially overcome by *F. fulva*.

## Results

### Genome sequencing of *F. fulva* strain IPO 2679 reveals two candidate *Avr9B* genes

As a starting point for the identification of *Avr9B*, the genome of New Zealand *F. fulva* strain IPO 2679, which has overcome *Cf-9B*-mediated resistance (Laugé *et al*., 1998), was sequenced (**Table S1**). A total of 119 previously identified (candidate) effector gene sequences from the *F. fulva* reference strain, 0WU (de Wit *et al*., 2012; Mesarich *et al*., 2014; Mesarich *et al*., 2018), which has not overcome *Cf-9B*-mediated resistance, were then compared with the corresponding gene sequences of strain IPO 2679 to identify those genes that have been mutated or deleted (**Table S2**). Sequence alignments revealed that 21 of the 119 genes have non-synonymous substitutions in IPO 2679, when compared to strain 0WU, while *Avr2* has a 140-base pair (bp) deletion, and a further two genes, including *Avr9*, have been deleted (**Table S2**).

Of the 24 genes that have been deleted or mutated in IPO 2679, two were of particular interest based on resistance-breaking mutations observed in other *Avr* effector genes of *F. fulva*. The first is *Ecp5* (GenBank ID: EF104527.1), a previously identified candidate *Avr* effector gene corresponding to the *Cf-Ecp5* resistance gene of tomato (Haanstra *et al*., 2000; Iakovidis *et al*., 2020; Laugé *et al*., 2000) that, like *Avr4* in many *Cf-4* resistance-breaking strains of *F. fulva* (Joosten *et al*., 1997), encodes a protein with a cysteine-to-tyrosine substitution (amino acid position 30 in Ecp5; **Table S2**). This non-synonymous substitution was subsequently confirmed by PCR amplicon sequencing. The second gene of interest is *CfCE54* (GenBank ID: KX943086.1) which, like *Avr9* in *Cf-9C*-resistance-breaking strains of *F. fulva* (van Kan *et al*., 1991), is deleted. A sequence alignment comparison to the recently published chromosome-level assembly of *F. fulva* strain Race 5 (Zaccaron *et al*., 2022), which is expected to carry a functional copy of *Avr9B*, estimated that the deleted region encompassing *CfCE54* in IPO 2679 is 5,429 bp in length (**Fig. S1a**). Deletion of this region was subsequently confirmed by PCR (**Fig. S1b**).

*CfCE54* is 566 nucleotides (nt) in length and comprises two introns and three exons (**Supplementary Information 1**). The gene is predicted to encode a protein of 152 amino acids with an N-terminal signal peptide of 21 amino acids for extracellular targeting, followed by a repeat-rich region made up of four direct imperfect 11-amino acid repeats and a cysteine-rich region with eight cysteine residues (**Fig. S2**). The repeat-rich region largely overlaps with a predicted intrinsically disordered region (IDR) (**Supplementary Information 1**). Like Ecp5, CfCE54 is not predicted to possess a transmembrane domain or a glycosylphosphatidylinositol (GPI) anchor modification site for attachment to the plasma membrane of fungal cells. Unlike Ecp5, CfCE54 was not identified by proteomic analysis in apoplastic washing fluid of *F. fulva*- infected tomato plants (Mesarich *et al*., 2018).

### *CfCE54* restores avirulence of *F. fulva* strain IPO 2679 on mature *Cf-9* tomato plants

To determine whether *Ecp5* or *CfCE54* is in fact *Avr9B*, complementation assays were carried out by introducing a single functional copy of the *Ecp5* or *CfCE54* gene from strain 0WU into strain IPO 2679 and then testing the ability of these complemented strains to cause disease on mature ‘Moneymaker’ (MM)-Cf-9 tomato plants carrying the *Cf-9* locus (**Fig. 1 and Fig. S3**). As expected, inoculation of mature MM-Cf-0 plants with the wild-type (WT) strains IPO 2679 (*Avr9*^−^/*Avr9B*^−^) and ICMP 7320 (*Avr9*^−^/*Avr9B*^+^), an IPO 2679 strain carrying the pFBTS1 empty vector (EV), or IPO 2679 strains complemented for *Ecp5* or *CfCE54*, resulted in disease (**Fig. 1 and Fig. S3**). Likewise, as anticipated, inoculation of mature MM-Cf-9 tomato plants with WT IPO 2679 or the IPO 2679 strain carrying the EV, but not WT ICMP 7320, also resulted in disease (**Fig. 1 and Fig. S3**). Notably, however, IPO 2679 strains complemented with *CfCE54*, but not with *Ecp5*, were unable to infect mature MM-Cf-9 plants (**Fig. 1 and Fig. S3**). As such, only *CfCE54* restores avirulence of strain IPO 2679 on *Cf-9* plants, indicating that *CfCE54*, but not *Ecp5*, is likely to be *Avr9B*.

**Figure 1.**
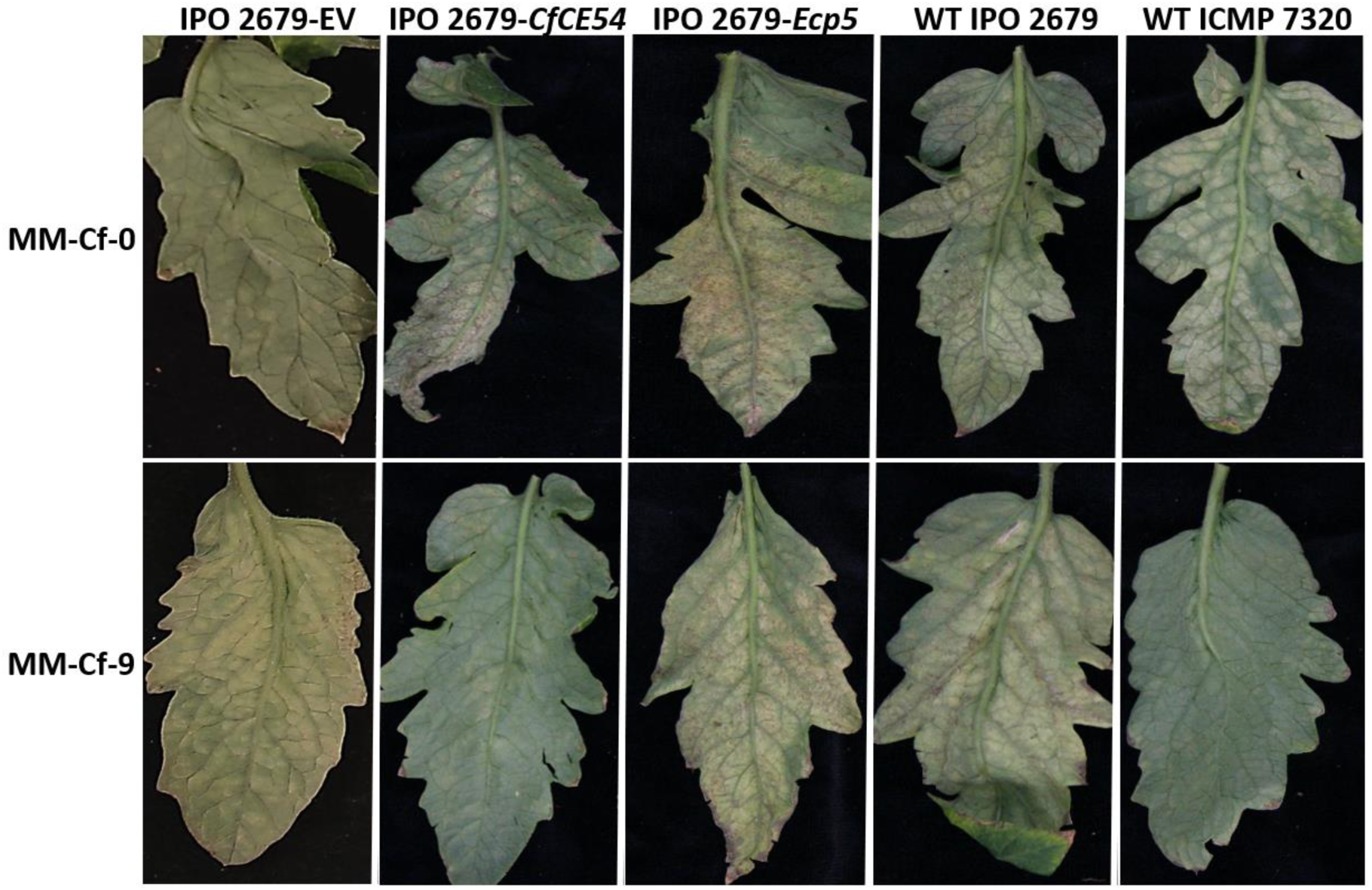
*CfCE54*, but not *Ecp5*, restores avirulence of *Fulvia fulva* strain IPO 2679 on mature *Solanum lycopersicum* plants carrying the *Cf-9* resistance locus. *F. fulva* strains were inoculated onto mature (13-week-old) ‘Moneymaker’ (MM)-Cf-0 (no *Cf* genes) and MM-Cf- 9 (carrying the *Cf-9* resistance locus) tomato plants, with photographs taken at 23 days post-inoculation. IPO 2679 EV, strain IPO 2679 (*Avr9*^−^/*Avr9B*^−^) complemented with the pFBTS1 empty vector (no insert); IPO 2679-*CfCE54*, strain IPO 2679 complemented with the *CfCE54* gene from wild-type (WT) strain 0WU; IPO 2679-*Ecp5*, strain IPO 2679 complemented with the *Ecp5* gene from WT strain 0WU; WT ICMP 7320, WT ICMP 7320 (*Avr9*^−^/*Avr9B^+^*) strain; WT IPO 2679, WT IPO 2679 strain. In total, two independent IPO 2679 EV, five independent IPO 2679-*CfCE54*, and two independent IPO 2679-*Ecp5* complementation transformants were tested. Complementation transformants #2 of IPO 2679 EV, #2 of IPO 2679-*CfCE54*, and *#*2 of IPO 2679-*Ecp5* are shown. Results are representative of all complementation strains tested.

### CfCE54 triggers a Cf-9B-dependent cell death response in *Nicotiana tabacum*

The MM-Cf-9 tomato plants used above in the complementation assays carry all five genes of the *Cf-9* resistance locus. Thus, it remains possible that an RLP encoded by this locus other than Cf-9B is responsible for recognizing CfCE54, leading to the observed disease resistance in mature plants. Notably, Cf-9B alone often triggers a cell death response in *Nicotiana benthamiana* but not in *Nicotiana tabacum* (Chakrabarti *et al*., 2009). Given the auto-activity of Cf-9B in *N. benthamiana*, we set out to confirm that CfCE54 from strain 0WU specifically triggers a Cf-9B-dependent cell death response upon co-expression with Cf-9B in *N. tabacum* using an *Agrobacterium tumefaciens*-mediated transient transformation assay (ATTA). Here, the CfCE54 (or Ecp5) protein, without its endogenous predicted signal peptide, was fused at its N-terminus to the PR1a signal peptide for extracellular targeting in *N. tabacum* to the apoplastic environment, followed by a 3xFLAG tag for detection by Western blotting. As expected, co-expression of Cf-9C with Avr9 (positive control; Hammond-Kosack *et al*., 1998) triggered a strong cell death response, while Avr9, Cf-9C or Cf-9B alone (negative controls) did not (**Fig. 2a**). Notably, CfCE54, but not Ecp5 (whether originating from strain 0WU or IPO 2679), triggered a strong cell death response upon co-expression with Cf-9B (**Fig. 2a,b**). This response was specific to Cf-9B, as CfCE54 did not trigger this strong response upon co-expression with Cf-9C (**Fig. 2a**). Taken together, these results confirm that CfCE54 is indeed Avr9B.

**Figure 2.**
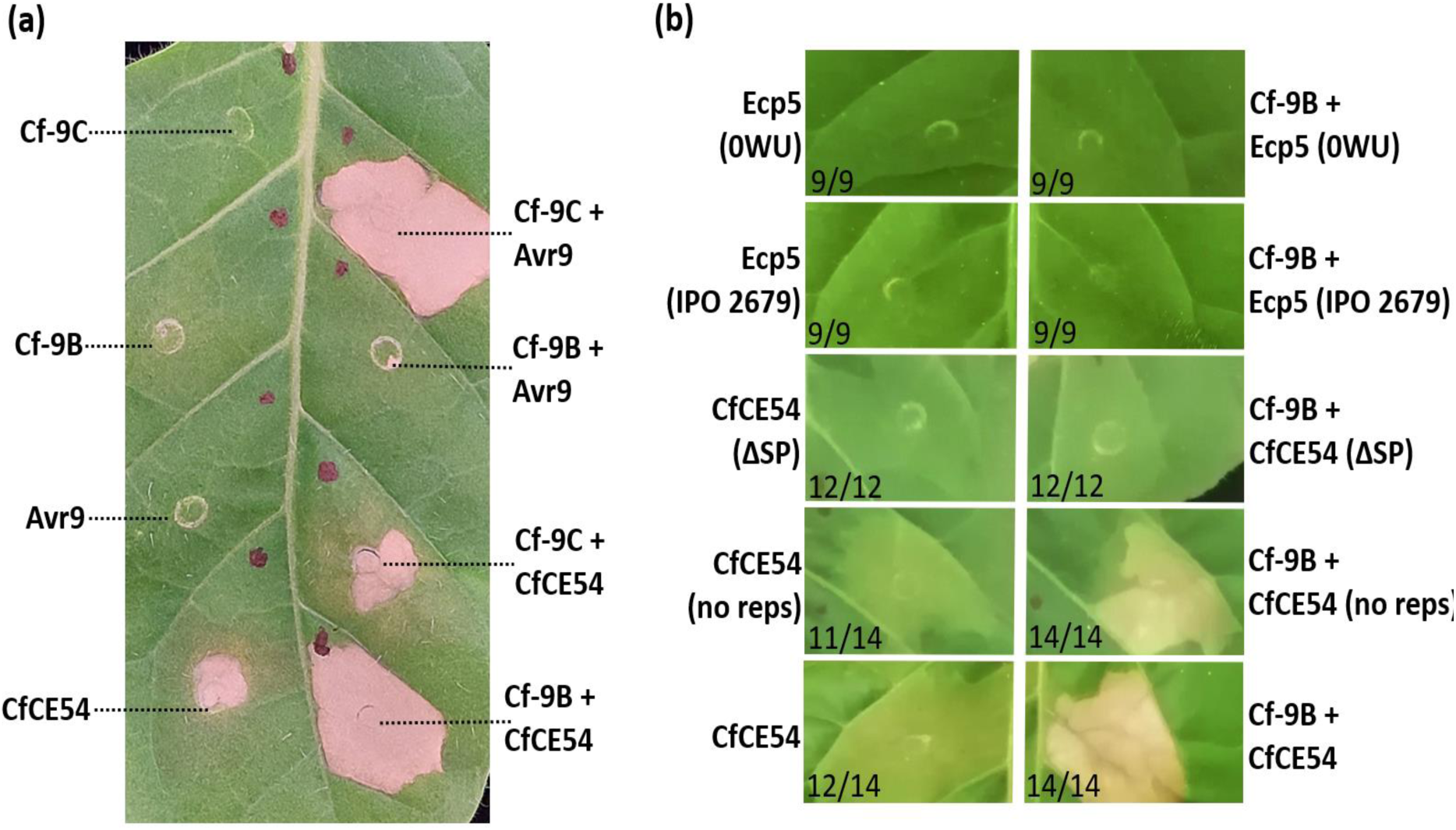
CfCE54, but not Ecp5, of *Fulvia fulva* triggers a Cf-9B-dependent cell death response in *Nicotiana tabacum*. Wild-type CfCE54 from strain 0WU **(a)**, or a variant of this protein with no signal peptide (ΔSP) or no repeat region (no reps), as well as wild-type Ecp5 from strain 0WU or a natural Cys30Tyr variant of this protein from strain IPO 2679 **(b)**, were co-expressed with Cf-9B or Cf-9C from tomato in *N. tabacum* using an *Agrobacterium tumefaciens*- mediated transient transformation assay (ATTA). As a positive control for cell death in (a), Cf- 9C was co-expressed with Avr9, whereas as negative controls for cell death in (a), Cf-9, Avr9 and Cf-9B were expressed alone. Please note that the ability of CfCE54 to trigger either chlorosis or a small patch of cell death when expressed alone varied from experiment to experiment (compare (a) and (b)). Leaves were photographed at 5 days post-infiltration and the results shown are representative of at least three independent ATTAs. In (b), numbers on the bottom left represent the number of times the response was observed (left) out of the number of times the infiltration was performed (right).

Interestingly, when Avr9B was expressed in *N. tabacum* either alone or in combination with any other of the proteins tested above, chlorosis and/or a small patch of cell death, was observed (**Fig. 2a,b**). Both the Cf-9B-dependent cell death response and the Cf-9B- independent chlorosis/cell death could be eliminated upon expression of an Avr9B variant lacking a signal peptide for secretion to the apoplastic environment (**Fig. 2b**), suggesting that extracellular targeting, or post-translational modifications associated with the endoplasmic reticulum–Golgi secretory pathway (e.g. disulphide bond formation and/or glycosylation), are required for these responses. Interestingly, a secreted version of Avr9B with the repeat region (part of the predicted IDR) deleted, maintained the ability to trigger Cf-9B-independent chlorosis/cell death and a Cf-9B-dependent cell death response (**Fig. 2b**), indicating that the four tandem repeats are not required for these responses in *N. tabacum*. Unfortunately, despite several attempts, at no stage following an ATTA could Avr9B or the version of this protein without a signal peptide or the repeat region be detected by Western blotting using antibodies against the 3xFLAG tag (**Fig. S4**). Ecp5 could, however, be detected by Western blotting (**Fig. S4**).

To determine whether the chlorotic/weak cell death response triggered by Avr9B, independent of Cf-9B, is dependent on the co-receptor SOBIR1, and thus is the result of recognition by an endogenous RLP in *N. tabacum*, we transiently expressed Avr9B in WT and Δ*sobir1 N. benthamiana* plants (Huang *et al*., 2021) using ATTAs, and compared the responses. As expected, the Cf-9C/Avr9 pair (Hammond-Kosack *et al*., 1998) triggered cell death in WT (positive control) but not Δ*sobir1 N. benthamiana* plants (negative control) (**Fig. S5**), consistent with the previous finding that cell death triggered by the Cf-9C/Avr9 pair is SOBIR1-dependent (Huang *et al*., 2021). In contrast to the chlorotic/weak cell death response observed in *N. tabacum*, Avr9B triggered a strong cell death response when expressed alone in WT *N. benthamiana* (**Fig. S5**). This response was also observed in Δ*sobir1* plants (**Fig. S5**), indicating that SOBIR1 is not required for the Cf-9B-independent cell death response triggered by Avr9B in *N. benthamiana*. Given this result, we anticipate that SOBIR1 is likely also not required for the Cf-9B-independent chlorosis/weak cell death response triggered by Avr9B in *N. tabacum*. Notably, as Avr9B alone triggered cell death in both WT and Δ*sobir1 N. benthamiana* plants, it was not possible to conclude whether cell death triggered by the Cf-9B/Avr9B pair is SOBIR1-dependent (**Fig. S5**).

### Avr9B triggers a Cf-9B-dependent cell death response in young *Cf-9* tomato plants, as well as chlorosis, weak necrosis and leaf curling in a wild tomato accession

To gain insights into whether the Cf-9B resistance pathway can be activated in young tomato plants, we systemically expressed Avr9B in young MM-Cf-9 tomato plants, as well as in young transgenic MM-Cf-0 lines expressing both *Cf-9A* and *Cf-9B* (*Cf-9A* + *Cf-9B*), using the Potato virus X (PVX)-based (pSfinx) expression system (Hammond-Kosack *et al*., 1995; Takken *et al*., 2000), and compared any responses to those observed in young MM-Cf-0 plants or a young transgenic MM-Cf-0 line expressing *Cf-9C*. For negative and positive controls, we also tested PVX alone (pSfinx empty vector, EV) and Avr9, respectively. As expected, PVX alone did not trigger an HR in any tomato line tested (**Fig. 3**), whereas Avr9 alone triggered a systemic HR in MM-Cf-9 and transgenic *Cf-9C* plants, as evidenced by light necrosis on the leaves and cotyledon drop (**Fig. 3**). Likewise, as anticipated, Avr9B failed to trigger an HR in MM-Cf-0 or transgenic *Cf-9C* plants (**Fig. 3**). Interestingly, Avr9B triggered a systemic HR in both MM-Cf-9 and transgenic *Cf-9A* + *Cf-9B* plants, which was even stronger than the response triggered by Avr9 in MM-Cf-9 and transgenic *Cf-9C* plants (**Fig. 3**). This indicates that the Cf-9B resistance pathway can be activated by Avr9B, in the absence of *F. fulva*, in young tomato plants. Curiously, systemic expression of Avr9B in MM-Cf-0 plants resulted in exaggerated PVX symptoms, when compared to EV-expressing control plants (**Fig. S6a**). These exaggerated PVX symptoms were not observed when several other characterized and candidate Avr effectors of *F. fulva* (Avr2, Avr4, Avr4E, Avr5, Avr9, Ecp5 and Ecp11-1) were systemically expressed in MM-Cf-0 tomato plants using the PVX-based expression system (**Fig. S6b**).

**Figure 3.**
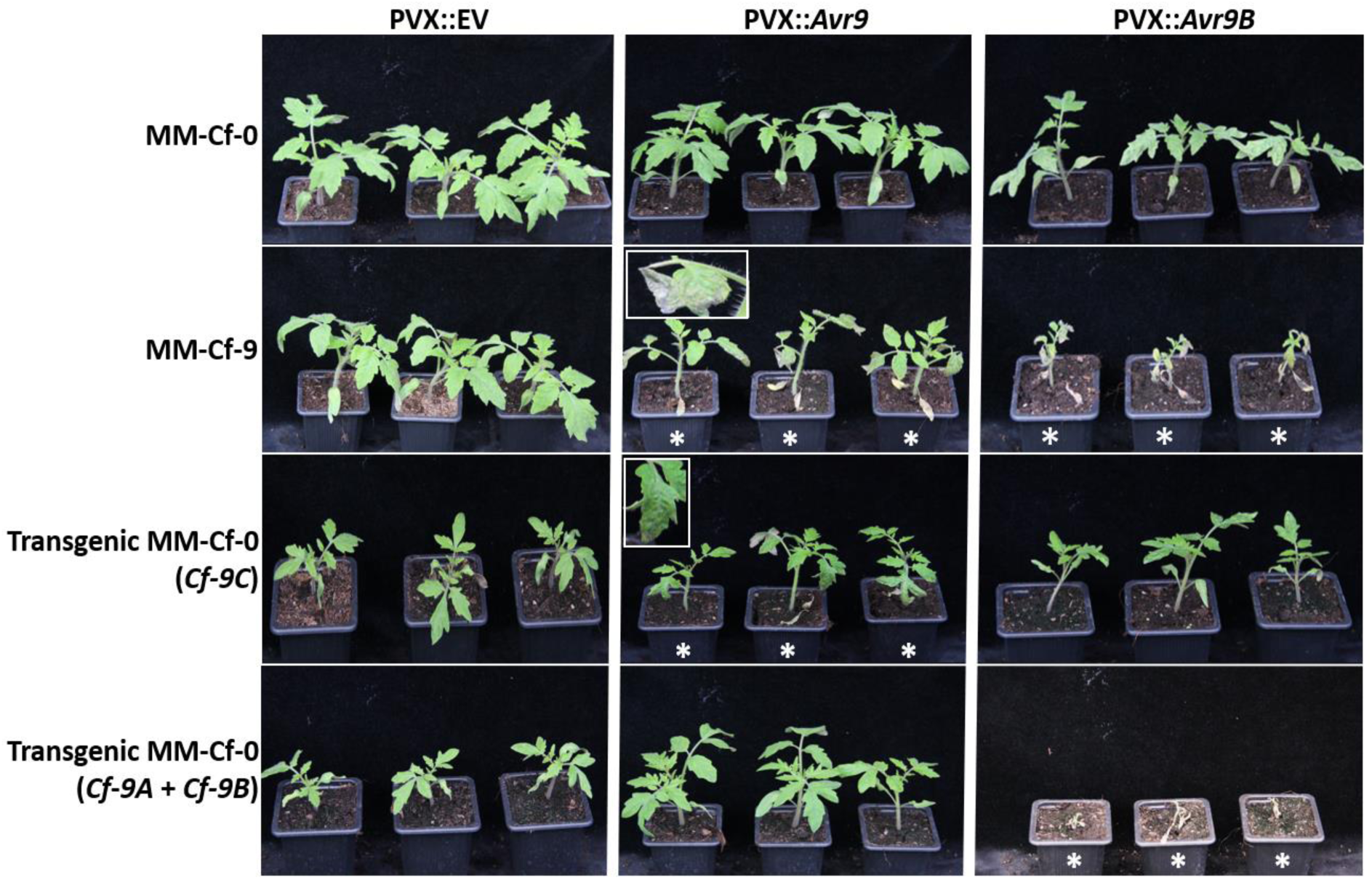
Avr9B triggers a hypersensitive response (HR) in young *Solanum lycopersicum* plants carrying the *Cf-9B* gene. Avr9 and Avr9B were systemically expressed in ‘Moneymaker’ (MM)-Cf-9 (carrying the *Cf-9* resistance locus), MM-Cf-0 (no *Cf* resistance genes) and transgenic MM-Cf-0 plants carrying either the *Cf-9C* gene or both the *Cf-9A* and *Cf-9B* genes (*Cf-9A* + *Cf-9B*), using the Potato virus X (PVX)-based expression system. Recombinant viruses PVX::*Avr9* and PVX::*Avr9B*, as well as PVX::EV (pSfinx empty vector), were delivered through cotyledon infiltration of 10-day-old tomato seedlings using *Agrobacterium tumefaciens*-mediated transient transformation. Three representatives of each tomato accession were included in the experiment. White asterisks indicate plants undergoing a systemic HR, as evidenced by necrosis (insets) and cotyledon drop. Photographs were taken at 10 days post- infiltration

In conjunction with the PVX-based expression system, we next sought to determine whether Avr9B triggers a systemic HR in wild tomato species, using the same collection of accessions tested in Mesarich *et al*. (2018). Based on this experiment, only one wild tomato accession (CGN14353; *S. pimpinellifolium*) responded to Avr9B, with leaves showing chlorosis, weak necrosis and inward curling relative to the EV-expressing control plants (**Fig. S7a,b**). Unlike that observed in MM-Cf-0 plants, there was no evidence that the PVX symptoms were exaggerated, however this may be influenced by the apparent defence responses triggered by Avr9B in this wild species.

### *Avr9B* expression is transcriptionally upregulated during infection of tomato by *F. fulva*

To determine whether *Avr9B* is transcriptionally upregulated during infection of tomato, relative to growth of the fungus in culture, a real-time quantitative polymerase chain reaction (RT-qPCR) experiment was performed using *F. fulva* strain Race 5. Based on this analysis, *Avr9B* expression was found to be strongly induced during infection, peaking at 4 and 8 days post-inoculation (dpi), with negligible expression at 12 and 16 dpi, or in culture during growth in potato dextrose broth (PDB) (**Fig. 4a**).

**Figure 4.**
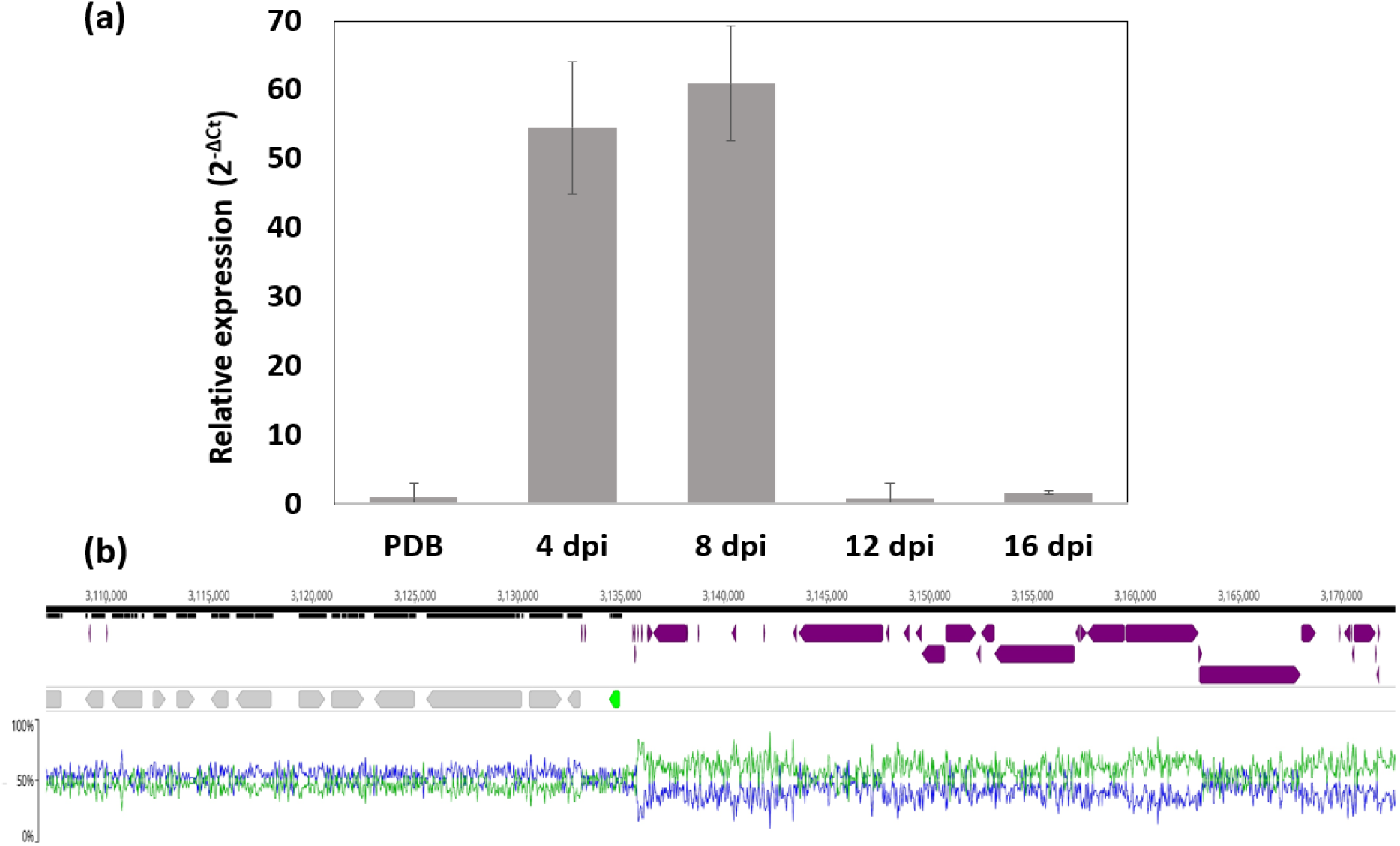
The *Avr9B* gene is transcriptionally upregulated during infection of tomato, relative to growth in culture, and is flanked by repetitive elements in the genome of *Fulvia fulva* strain Race 5. (a) Expression profile of *Avr9B*. Expression was analysed using a real-time quantitative polymerase chain reaction (RT-qPCR) experiment, from a compatible *F. fulva* Race 5– *Solanum lycopersicum* ‘Moneymaker’ (MM)-Cf-0) interaction at 4, 8, 12 and 16 days post-inoculation (dpi), as well as in culture in potato dextrose broth (PDB) at 4 dpi. Expression was normalized to the *F. fulva actin* gene according to the 2^-ΔCt^ method. Error bars represent the standard deviation across three biological replicates. **(b)** Location of *Avr9B* in the genome of *F. fulva* strain Race 5. A zoomed-in region of Chromosome 10 is shown. The *Avr9B* gene (encoded by CLAFUR5_14678; National Centre for Biotechnology Information accession: XP_047767027.1) is shown in green, while the other genes present in the region are shown in grey. Repetitive elements are shown in purple. The G+C content of the genomic region is shown by a blue line, while the A+T content is shown by a green line.

### *Avr9B* is adjacent to repetitive elements in the genome of *F. fulva* strain Race 5

To determine whether the *Avr9B* gene is associated with repeats in the *F. fulva* genome, its position relative to previously annotated repetitive elements, identified as part of the strain Race 5 chromosome-level genome assembly (Zaccaron *et al*., 2022), was investigated. This revealed that *Avr9B* resides on chromosome 10 of the strain Race 5 genome sequence (location 3134996–3134431) and is adjacent to a region rich in repeats at its 5′ end (**Fig. 4b**).

### Avr9B-like proteins are restricted to members of the Mycosphaerellaceae and Pleosporaceae

To determine whether Avr9B-like proteins are present in other fungi, a BLAST (basic local alignment search tool) analysis was performed against fungal genomes and proteins present in the National Centre for Biotechnology Information (NCBI) and the Joint Genome Institute (JGI) Mycocosm databases using Avr9B (as well as any identified Avr9B-like protein) as a query. This analysis revealed that genes encoding Avr9B-like proteins are exclusively present in plant-pathogenic species from the Mycosphaerellaceae and Pleosporaceae families of the Dothideomycetes class of fungi, with several found in tomato pathogens (**Fig. 5a**). An alignment of all predicted Avr9B-like proteins revealed three main groups. Group 1 proteins, of which one is Avr9B itself, are characterised by eight conserved cysteine residues, while Group 2 proteins are characterized by ten conserved cysteine residues. For the latter, the first eight cysteine residues are shared with Group 1 proteins, while the last two cysteine residues form part of a C-terminal extension. Finally, Group 3 proteins are characterized by eight conserved cysteine residues, with the first six cysteine residues shared with Group 1 and 2 proteins, and the last two cysteine residues shared with the C-terminal extension of Group 2 proteins. Some members of Group 3 also possess an additional cysteine residue (**Fig. 5a and Supplementary Information 1**). All Avr9B-like proteins are predicted to possess an N-terminal signal peptide for extracellular targeting, while only the Avr9B-like proteins from Group 3 are predicted to possess a transmembrane domain (**Supplementary Information 1**). Like Avr9B, many Avr9B-like proteins are also predicted to possess an N-terminal IDR (**Supplementary Information 1**).

**Figure 5.**
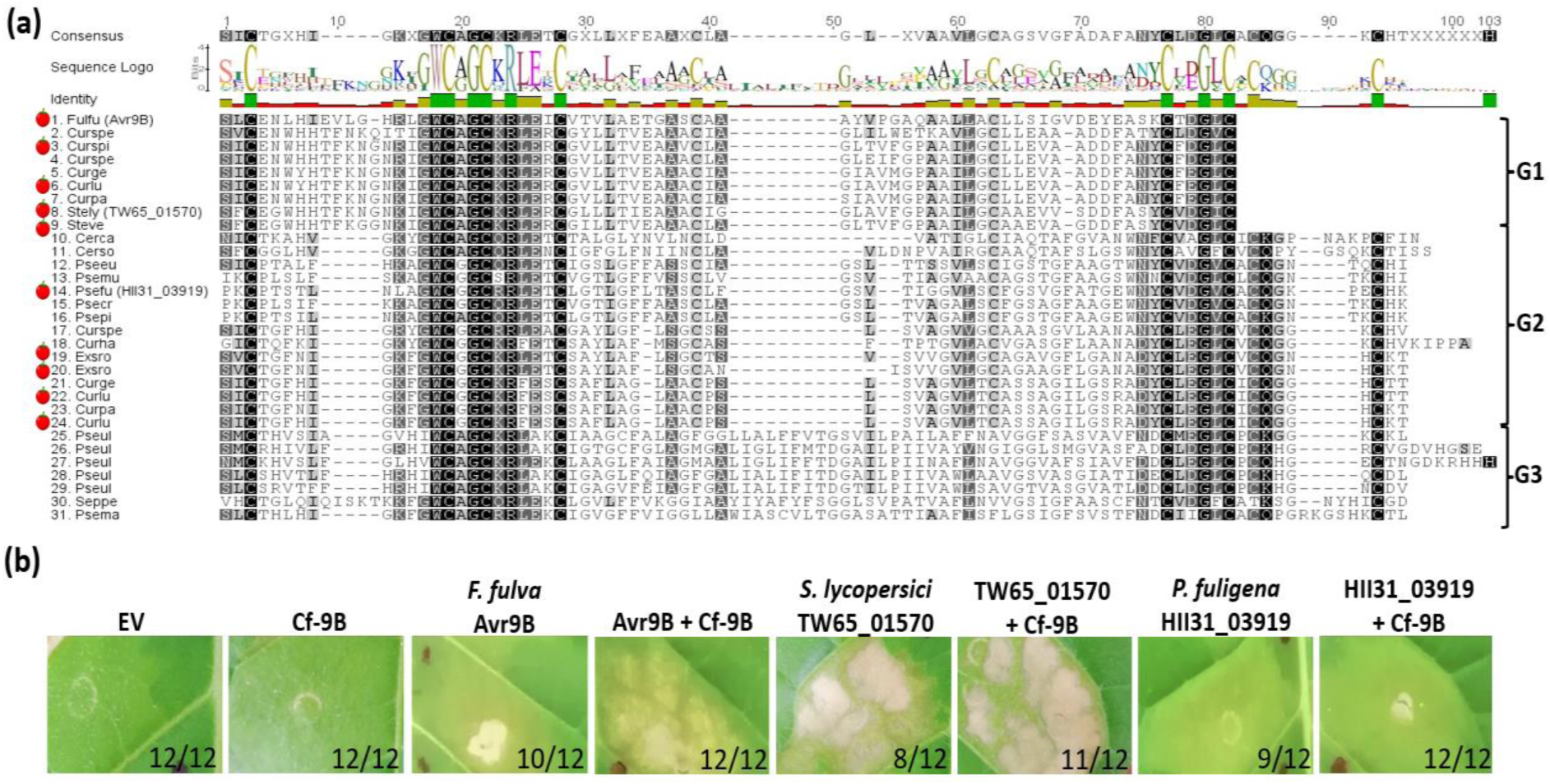
Avr9B-like proteins are restricted to plant-pathogenic Dothideomycete fungi and trigger chlorosis or cell death in *Nicotiana tabacum*. **(a)** Alignment of Avr9B-like proteins from Dothideomycete fungi. Only the cysteine-rich region from each protein is aligned. Group 1 (G1) proteins, sequences 1–9; Group 2 (G2) proteins, sequences 10–24; Group 3 (G3) proteins, sequences 25–31. Cerca, *Cercospora canescens*; Cerso, *Ce*. *sojina*; Curge, *Curvularia geniculata*; Curha, *Cu*. *hawaiiensis*; Curlu, *Cu*. *lunata*; Curpa, *Cu*. *papendorfii*; Curspe, *Cu*. sp. ZM96; Curspi, *Cu*. *spicifera*; Exsro, *Exserohilum rostratum*; Fulfu, *Fulvia fulva*; Psecr, *Pseudocercospora cruenta*; Pseeu, *P*. *eumusae*; Psefu, *P*. *fuligena*; Psema, *P*. *macadamiae*; Psemu, *P*. *musae*; Psepi, *P*. *pini-densiflorae*; Pseul, *P*. *ulei*; Seppe, *Septoria petroselini*; Stely, *Stemphylium lycopersici*; Steve, *St*. *vesicarium*. Tomato pathogens are indicated by a tomato figure on the left. **(b)** The Avr9B-like proteins from *S. lycopersici* and *P. fuligena* trigger strong cell death and chlorosis in *N. tabacum* plants, respectively. Avr9B-like proteins TW65_01570 (Franco *et al*., 2015) and HII31_03919 (Zaccaron *et al*., 2020) from *S. lycopersici* and *P. fuligena*, respectively, were either expressed alone or co-expressed with Cf-9B in *N. tabacum* using an *Agrobacterium tumefaciens*-mediated transient transformation assay (ATTA). As positive controls for cell death, Avr9B was either expressed alone (chlorosis/weak cell death) or together with Cf-9B (strong cell death). As negative controls, empty vector (EV; pICH86988) and Cf-9B were expressed alone. Photos were taken at 5 days post-infiltration. Fractions on the bottom right represent the number of times the response was observed (left) out of the number of times the infiltration was performed (right) across at least three biological replicates.

### Avr9B-like proteins trigger responses in *Nicotiana benthamiana, Nicotiana tabacum* and *Solanum lycopersicum* in the absence of Cf-9B

To determine whether Avr9B-like proteins from other fungi also trigger a Cf-9B-dependent cell death response, we expressed the proteins identified from the tomato pathogens *Stemphylium lycopersici* (TW65_01570, NCBI accession: KNG51100.1; Franco *et al*., 2015) and *Pseudocercospora fuligena* (HII31_03919, NCBI accession: KAF7194657.1; Zaccaron *et al*., 2020) in *N. tabacum* using an ATTA (**Fig. 5b**). TW65_01570 triggered a strong cell death response by itself, as well as when co-expressed with Cf-9B or Cf-9C (**Fig. 5b**). Hence, it was not possible to determine whether TW65_01570 is recognized by Cf-9B in *N. tabacum*. Similar to Avr9B, HII31_03919 triggered a chlorotic response alone in *N. tabacum* (**Fig. 5b**). However, unlike Avr9B, a cell death response was not observed when HII31_03919 was co-expressed with Cf-9B (**Fig. 5b**). As recognition could have theoretically resulted in chlorosis, and because chlorosis is triggered by HII31_03919 independently of Cf-9B, it was again not possible to determine whether HII31_03919 is recognized by Cf-9B in *N. tabacum*. Unlike Avr9B, both Avr9B-like proteins could be detected by Western blotting (**Fig. S4**). Similar to Avr9B, both Avr9B-like proteins triggered a strong cell death response in the absence of Cf-9B in *N. benthamiana*, with these responses not dependent on SOBIR1 (**Fig. S5**).

To determine whether the Avr9B-like proteins trigger a Cf-9B-dependent HR in tomato, we systemically expressed TW65_01570 in MM-Cf-9 plants using the PVX-based expression system and compared the responses to those observed in MM-Cf-0 (**Fig. S8**). In contrast to expression of Avr9B, no Cf-9B-dependent HR was observed in MM-Cf-9 plants, suggesting that TW65_01570 is not recognized by Cf-9B (**Fig. S8**). Surprisingly, however, systemic expression of TW65_01570 consistently resulted in stunted growth regardless of the plant genotype (**Fig. S8**). This stunting was most obvious when TW65_01570 was systemically expressed in the wild *S. pimpinellifolium* accession CGN14353 (**Fig. S7a**).

### The *Cf-9* resistance locus of tomato has been sequentially broken down by *F. fulva*

To determine whether resistance provided by the *Cf-9* locus in tomato has been sequentially broken down by *F. fulva*, with *Cf-9C* overcome prior to *Cf-9B*, the *Avr9* and *Avr9B* genes were screened for deletion or mutation across a collection of 190 geographically diverse *F. fulva* strains, using PCR followed by PCR amplicon sequencing (**Table S3**). Of these strains, 149 were collected relatively recently (during or after 2005), and originated from Japan (99), France (27), China (six), Germany (three), New Zealand (three), the Netherlands (two) and Tanzania (one). A further 33 strains were collected prior to 1990, and originated from the Netherlands (17), France (nine), Poland (three), New Zealand (three) and Belgium (one). Of the strains screened, eight were of unknown origin and six did not have a collection date. In addition to strain 0WU from the Netherlands, which was collected in 1997, one strain from Japan was also collected in 1998. In total, 52 of all strains screened are known to have been isolated from *Cf-9*/Ff:A-E tomato plants (**Table S3**).

The PCR analysis determined (or confirmed from previous studies) that 93 of the *F. fulva* strains that were screened, including all strains isolated from *Cf-9*/Ff:A-E tomato plants, were race 9, as the *Avr9* gene was absent (**Table S3**). Notably, in nine out of 11 race 9 strains collected prior to 1990 that lacked the *Avr9* gene (i.e. except strains IPO 2679 from New Zealand and Race 1 from the Netherlands), no mutations were observed in the coding sequence of *Avr9B* (**Table S3**). Strikingly, however, mutations in *Avr9B* were observed in all but one of 76 race 9 strains collected during or after 2005, with the only exception being strain 2.4.9 from Japan, which was collected in 2018 (**Table S3**). These mutations included premature stops associated with codons 2 (p.Arg2*), 55 (p.Ser55*), 83 (p.Cys83*), and 107 (p.Cys107*), as well as a frame-shift mutation in codon 64, leading to an alteration in protein sequence and a premature stop after position 69 (p.Gly64Valfs*70) (**Fig. 6a and Table S3**). Additional mutations identified were an amino acid substitution associated with codon 97 (p.Trp97Cys), as well as an insertion of a 289-bp miniature inverted-repeat transposable element (MITE) located in codon 124 (**Fig. 6a and Table S3**), leading to an alteration in protein sequence and a stop after position 185 (p.Val123_Pro124ins*63), and an 18-bp deletion starting in codon 128, resulting in the replacement of seven amino acids (including Cys 133) with a single Val residue (p.Ala128_Leu134delinsVal) (**Fig. 6a and Table S3**). In the case of the MITE, a single identical copy was found approximately 1.2 Mb away from *Avr9B* at location 1915484– 1915772 on chromosome 10 in the strain Race 5 genome, suggesting it may originate from this location (**Fig. S9**). Remarkably, no synonymous mutations were observed in the coding sequence of the *Avr9B* gene between strains (**Table S3**). Interestingly, only one non-race 9 strain (Kaminokawa, isolated in Japan in 2016) was predicted to possess a WT *Avr9* gene but a mutant *Avr9B* gene (**Table S3**).

**Figure 6.**
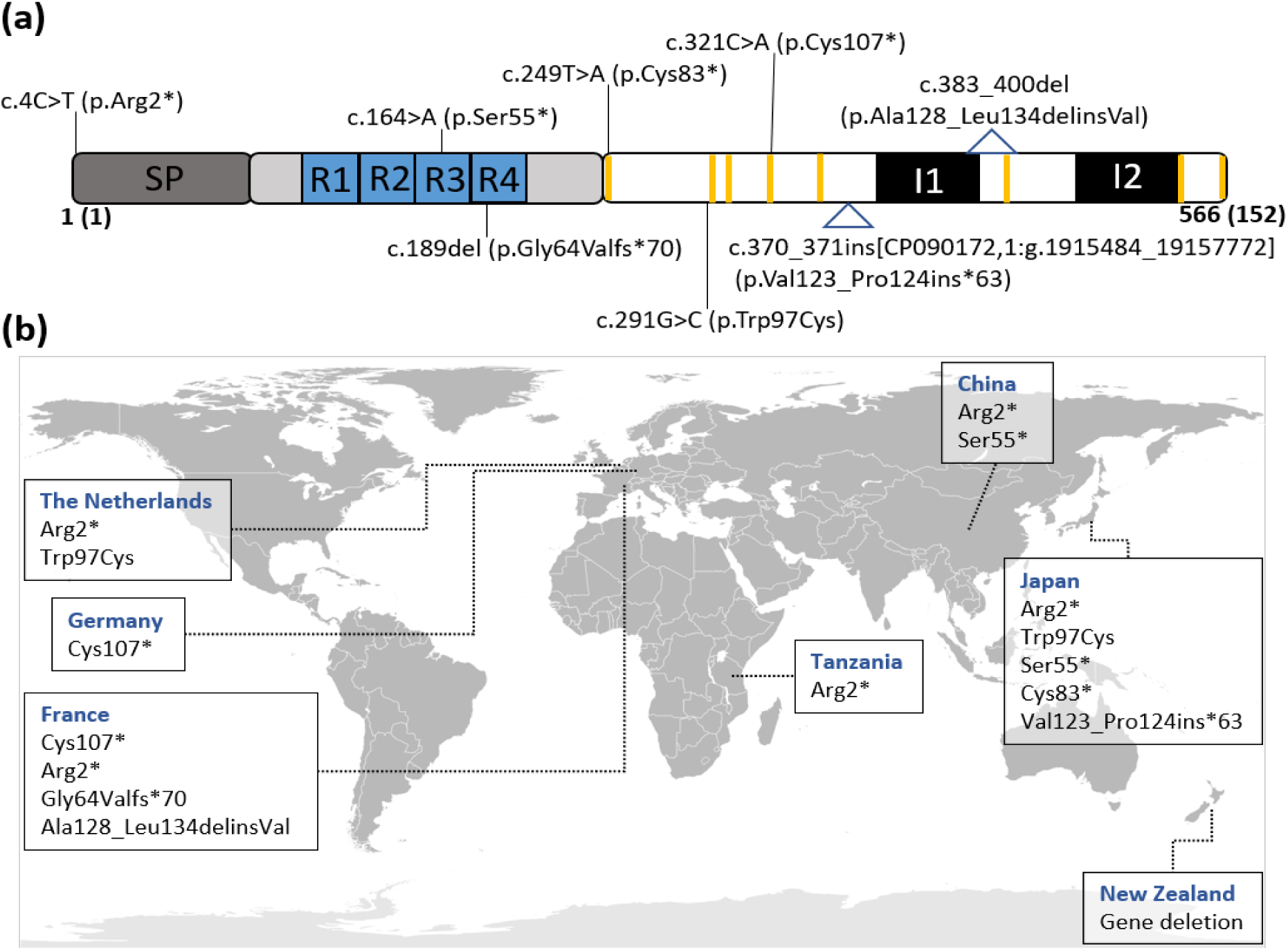
Mutations identified in Avr9B and distribution of these mutations across *Fulvia fulva* strains collected from around the world. **(a)** Allelic variation in Avr9B of *Cf-9B*-resistance- breaking strains. The predicted signal peptide (SP) for extracellular targeting is highlighted in dark grey. The four imperfect 11-amino acid tandem repeats (R1–R4) are shown as blue squares. The cysteine residues are shown in vertical yellow lines. The predicted intrinsically disordered region (IDR) is highlighted in light grey. The introns (I1 and I2) are shown as black boxes. c., coding sequence numbering; p., protein sequence numbering. **(b)** Distribution of Avr9B mutations across strains collected from around the world. The description of each mutation is based on the nomenclature set forth by the human genome variation society (HGVS) (http://varnomen.hgvs.org/). CP090172.1:g.1915484_1915772 represents the likely original location of the miniature inverted-repeat transposable element (MITE) on chromosome 10 of the *F. fulva* strain Race 5 genome (Zaccaron *et al*., 2022). del, deletion; delins, deletion- insertion; fs, frameshift; ins, insertion. Global base map generated by F. Bennet in the public domain and accessed from: https://commons.wikimedia.org/wiki/File:BlankMap-FlatWorld6.svg.

Of the mutations identified, all mutations except p.Arg2*, which was identified in five countries, were restricted to one or two countries only (**Fig. 6b**). Interestingly, the *Avr9B* gene deletion, as seen in IPO 2679, was restricted to New Zealand strains only, with the same region of deleted sequence shown to be absent by PCR in IPO 2679 collected in the 1980s as well as in strains NZ-C1, NZ-P1 and NZ-M6 collected in 2022 (**Fig. S1b**).

In addition to investigating *Avr9* and *Avr9B*, the mating type of each strain was determined by PCR or obtained from previous research (Stergiopoulos *et al*., 2007a; Stergiopoulos *et al*., 2007b). Based on this analysis, all mutation types were found to be restricted to strains of a particular mating type, except for p.Ser55*, which was observed in strains of both mating types that were predominantly collected from the Gifu prefecture of Japan (**Table S3**).

Taken together, the PCR and PCR amplicon sequencing results suggest that *Cf-9C*- mediated resistance was overcome first, with most strains collected prior to 1990 lacking *Avr9* but still carrying a functional copy of *Avr9B*, and *Cf-9B*-mediated resistance was overcome second, with strains collected during or after 2005 lacking *Avr9* and possessing a resistance-breaking mutation in *Avr9B*. Furthermore, the mating type data suggest that specific mutations have evolved once in *F. fulva*, with a strong correlation between a given mutation and a particular mating type.

### The W97C mutation in Avr9B restores virulence of *F. fulva* on mature MM-Cf-9 tomato plants

All mutations identified in Avr9B, except for the p.Trp97Cys amino acid substitution, hereafter referred to as W97C, could be confidently assumed to result in the circumvention of Cf-9B-mediated resistance by *F. fulva* on mature *Cf-9* tomato plants. To test whether the W97C mutation also leads to the restoration of *F. fulva* virulence on tomato harbouring the *Cf-9* resistance locus, strain P31 (*Δavr9*/*Avr9B^W97C^*) from Japan was inoculated onto mature MM-Cf-0 and MM-Cf-9 plants and its ability to cause disease was assessed (**Fig. 7a**). At the same time, strain P18 (*Δavr9*/*Avr9B^C107*^*) from Germany, which is expected to overcome both Cf- 9C- and Cf-9B-mediated resistance in mature *Cf-9* tomato plants due to deletion of *Avr9* and a p.Cys107* mutation, hereafter referred to as C107*, in Avr9B, was also tested. As expected, both the P31 and P18 strains caused disease on MM-Cf-0 plants (**Fig. 7a and Fig. S10**). Notably, both strains were also able to cause disease on MM-Cf-9 plants (**Fig. 7a and Fig. S10**), indicating that, like the C107* mutation in strain P18, the W97C mutation enables the P31 strain to evade recognition by Cf-9B and thus restores virulence of *F. fulva* on MM-Cf-9 plants. To support this observation, we tested the W97C mutant of Avr9B for its ability to trigger a Cf-9B-dependent cell death response in *N. tabacum* plants using an ATTA. In line with the gene complementation assay, the W97C mutant of Avr9B was unable to trigger a Cf- 9B-dependent cell death response in *N. tabacum* (**Fig. 7b**).

**Figure 7.**
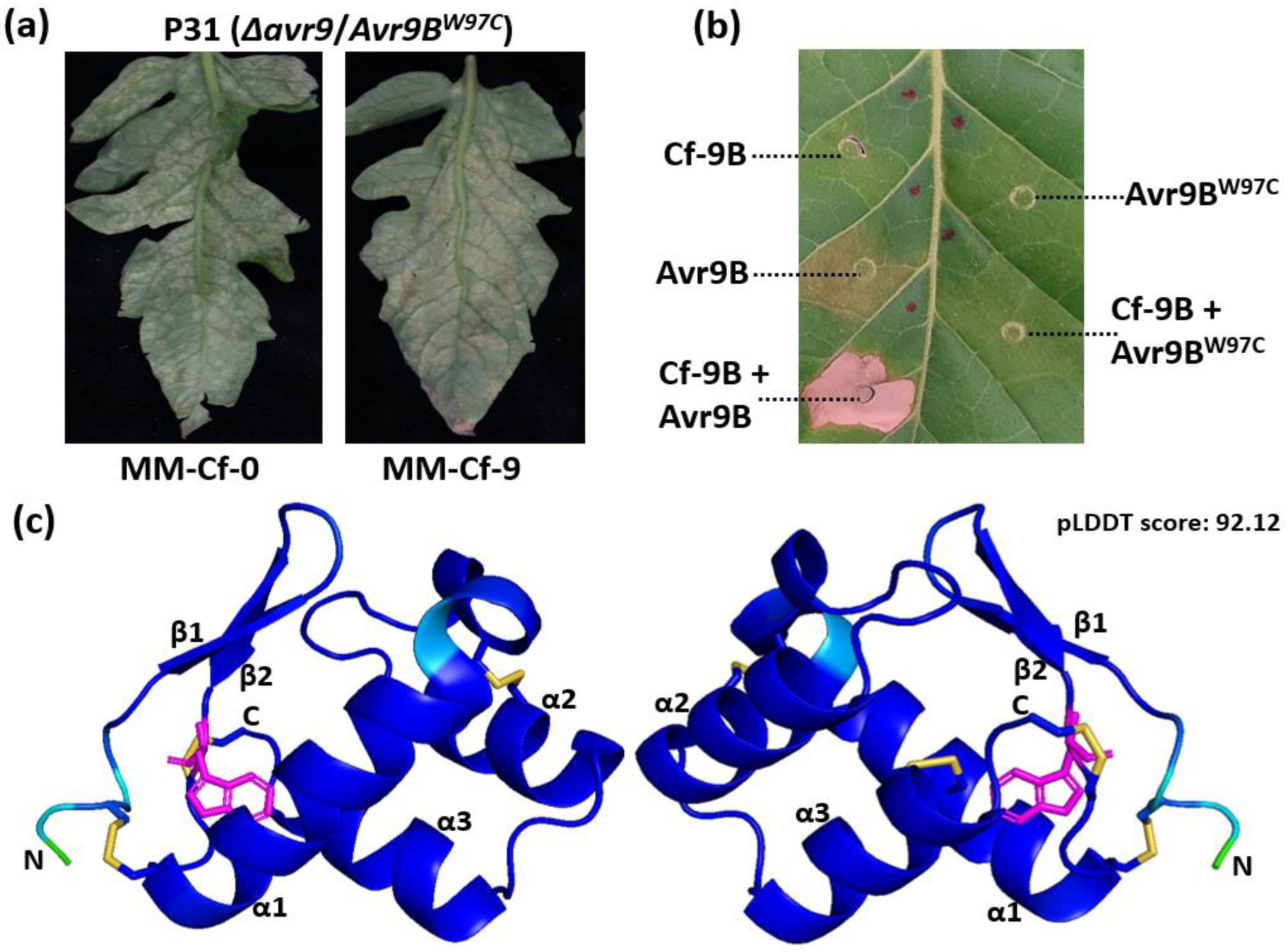
The W97C mutation in Avr9B results in circumvention of Cf-9B-mediated resistance in *Solanum lycopersicum* and the inability to trigger both Cf-9B-dependent cell death and Cf-9B-independent chlorosis/cell death in *Nicotiana tabacum*. **(a)** *F. fulva* strain P31 (*Δavr9*/*Avr9B^W97C^*) was inoculated onto mature (13-week-old) ‘Moneymaker’ (MM)-Cf-0 (no *Cf* genes) and MM-Cf-9 (carrying the *Cf-9* resistance locus) plants, with photographs taken at 23 days post-inoculation. Results are representative of three biological replicates and the inoculations were performed at the same time as the virulence assays shown in Fig. 1. **(b)** The W97C mutant of Avr9B was co-expressed with Cf-9B in *N. tabacum* using an *Agrobacterium tumefaciens*-mediated transient transformation assay (ATTA). As positive controls for cell death, Avr9B was expressed alone (chlorosis/weak cell death) or together with Cf-9B (strong cell death). As a negative control for cell death, Cf-9B was expressed alone. The leaf was photographed five days post-infiltration and is representative of three independent ATTAs. **(c)** Predicted tertiary structure of the cysteine-rich region from Avr9B, rotated 180° around its y-axis, showing that Trp 97 is unlikely to be surface-exposed. The tertiary structure was predicted in conjunction with a multiple sequence alignment of Avr9B-like proteins from Fig. 5a and is coloured according to predicted Local Distance Difference Test (pLDDT) score (confidence: red = very low, yellow = low, green = okay, cyan = high, dark blue = very high). Trp 97 is shown as sticks and is coloured magenta. β-strands and α-helices are numbered sequentially. Predicted disulphide bonds are coloured yellow. N- and C**-**termini are indicated.

To gain insights into whether Trp 97 is predicted to be surface-exposed, AlphaFold2 (Jumper *et al*., 2021), in conjunction with ColabFold (Mirdita *et al*., 2022), was used to predict the tertiary structure of the cysteine-rich region of Avr9B. The predicted structure is characterized by three α-helices and two β-strands and is stabilized by four disulphide bonds (**Fig. 7c**). Based on this tertiary structure, all eight cysteine residues are predicted to be disulphide-bonded with the connectivity pattern Cys83-Cys101, Cys98-Cys152, Cys107- Cys147, and Cys118-Cys133, based on numbering of the full-length protein sequence. Comparison of the predicted structure to other solved tertiary structures revealed no significant structural similarity to other proteins. Intriguingly, Trp 97 is not predicted to be surface- exposed but, rather, to occupy an internal position within the predicted Avr9B tertiary structure (**Fig. 7c**).

## Discussion

The *Cf-9* locus, harbouring the *Cf-9B* and *Cf-9C* resistance genes, is present worldwide in most commercially available tomato cultivars that have resistance to leaf mould disease. While the *Cf-9C* resistance gene, which provides protection during all stages of plant growth, was rapidly broken down soon after deployment of the *Cf-9* locus in the 1970s (e.g. Laterrot, 1986; Lindhout *et al*., 1989), the *Cf-9B* resistance gene, which provides protection to mature plants during flowering and fruiting, proved to be more durable. Over the last 20 years, however, the incidence of leaf mould disease on fruiting and flowering plants has increased. While the breakdown of *Cf-9C*-mediated resistance has been shown to be the result of *Avr9* gene deletion in *F. fulva* strains (Bernal-Cabrera *et al*., 2021; Iida *et al*., 2015; Stergiopoulos *et al*., 2007a; Yoshida *et al*., 2021; van Kan *et al*., 1991), the molecular mechanism underlying *Cf-9*B-mediated resistance breakdown remained unclear. To understand how *Cf-9B*-mediated resistance is circumvented, we identified the corresponding *Avr9B* gene of *F. fulva* using a comparative genomics approach based on candidate effector genes identified in previous studies (Mesarich *et al*. 2014; Mesarich *et al*., 2018).

Like other *Avr* effector genes of *F. fulva* identified to date, *Avr9B* is up-regulated during infection of tomato and encodes a small, secreted cysteine-rich protein. Despite these similarities, Avr9B is unique among *F. fulva* Avr effector proteins in that it is predicted to have a large, repeat-rich IDR at its N-terminus and, therefore, can be classified as a repeat-containing effector protein (Mesarich *et al*., 2015). These repeats, however, do not appear to be necessary for recognition by Cf-9B, because an ATTA, using a version of Avr9B without this region, did not result in compromised Cf-9B-mediated cell death in *N. tabacum*. As such, the C-terminal cysteine-rich part of the protein appears to be the relevant region for recognition by Cf-9B. In line with Avr9B being recognized by an extracellular immune receptor, such as an RLP, Cf- 9B-mediated cell death was abolished in *N. tabacum* when a version of Avr9B lacking a signal peptide for secretion to the apoplast was tested. It must be pointed out, however, that as Avr9B did not pass through the ER–Golgi secretory pathway, it would not have undergone appropriate post-translational modifications such as disulphide bond formation, which are anticipated to be required for stability, function and/or recognition.

A closer inspection of the genomic region surrounding the *Avr9B* gene in the recently assembled genome sequence of *F. fulva* strain Race 5 (Zaccaron *et al*., 2022) revealed its proximity to repetitive elements, in common with several other *Avr* effector genes of *F. fulva* identified to date, including *Avr4E*, *Avr5* and *Avr9* (Zaccaron *et al*., 2022). Interestingly, like these three *Avr* effector genes (van Kan *et al*., 1991; Westerink *et al*., 2004; Mesarich *et al*., 2014), the *Avr9B* effector gene was also found to be deleted in some strains of *F. fulva*. This is perhaps not surprising, as it has been reasoned that gene deletion events are largely influenced by the proximity of *Avr* effector genes to repetitive elements (Gout *et al*., 2007). Eight other confirmed or presumed *Cf-9B*-breaking mutations were identified in the *Avr9B* gene sequence of *F. fulva* strains collected from different geographical locations and were made up of four premature stop codons, one point mutation resulting in a non-synonymous amino acid substitution (W97C), one indel resulting in a five-amino acid deletion (including a cysteine residue predicted to be disulphide-bonded), one frame-shift mutation, and one MITE insertion. Curiously, in the case of the W97C substitution, the tryptophan residue in question was not predicted to be surface-exposed, in line with it being a large hydrophobic amino acid well-suited to the hydrophobic core of proteins. As cysteine is a much smaller hydrophobic amino acid than tryptophan, it is possible that the W97C substitution negatively affects the folding and/or stability of Avr9B, such that it is no longer functional and/or recognized by Cf-9B.

While most resistance-breaking mechanisms described above are not uncommon for *Avr* effector genes of *F. fulva*, this is the first time where circumvention of *Cf*-mediated resistance involving a MITE insertion has been reported. Another MITE insertion has, however, been previously found in an *Avr* effector gene of the stem rust fungus, *Puccinia graminis* f. sp. *tritici* (*AvrSr35*), enabling this pathogen to overcome resistance mediated by the corresponding *Sr35* resistance gene in wheat (Salcedo *et al*., 2017). MITEs are a group of non-autonomous Class II transposable elements that do not encode their own transposase but, instead, can commandeer transposases from other mobile genetic elements for their mobilization (Wicker *et al*., 2007). For *F. fulva*, inspection of the strain Race 5 genome revealed that the MITE in question likely originated from a gene-rich region of the genome that is approximately 1.2 Mb away from *Avr9B* on chromosome 10.

Several of the *Cf-9B*-breaking mutations identified in *Avr9B* were restricted to specific geographical areas, indicating independent mutation events around the world. In line with this, deletion of *Avr9B* (and more specifically, the same deletion profile) was, for example, only detected in a New Zealand strain collected in the 1980s and then again in strains collected in 2022. The finding that nine independent and mostly geographically specific *Cf-9B*-breaking mutations have so far been identified in the *Avr9B* gene is perhaps not surprising, given the extensive deployment of tomato cultivars carrying the *Cf-9* resistance locus around the world. Remarkably, the p.Ser55* mutation observed in Japan is present in strains of *F. fulva* that carry opposite mating type genes, suggesting that this mutation has independently evolved twice, or that sexual recombination has occurred. In support of the latter, the strains of opposite mating type that share the p.Ser55* mutation are from the same Japanese prefecture (Gifu). While *F. fulva* was originally thought to be an asexual fungus (Thomma *et al*., 2005), this information, together with allelic variation data collected from other *Avr* effector genes of this fungus (Stergiopoulos *et al*. 2007b), suggests that *F. fulva* does indeed undergo sexual reproduction, albeit rarely. Future experiments can now investigate this further by looking for evidence of meiotic recombination between the abovementioned *F. fulva* strains collected from this prefecture.

An interesting observation in our study was that, when Avr9B was systemically expressed in young tomato plants using the PVX-based expression system, the Cf-9B resistance pathway was found to be active. Indeed, the response triggered by Avr9B in *Cf-9* plants was stronger in our hands than the response triggered by Avr9. This is despite the fact that Cf-9B only provides resistance to *F. fulva* in mature tomato plants at flowering and fruiting but is in line with the finding that *Cf-9B* expression is not developmentally regulated (Laugé *et al*., 1998; Panter *et al*., 2002). Of note, although *Avr9B* is highly expressed during infection of tomato, the protein was not detected in apoplastic washing fluid samples collected from young *F. fulva*-infected tomato leaves (Mesarich *et al*., 2018). Likewise, the Avr9B protein could not be detected in total protein samples from *N. tabacum* ATTAs using Western blotting. Together, these findings may suggest that the stability (and thus abundance) of Avr9B is compromised in young tomato plants, rendering Cf-9B-mediated resistance ineffective. Correspondingly, in mature tomato plants, presumably due to qualitative and/or quantitative differences in the apoplastic protease profile, Avr9B is more stable (and thus more abundant), rendering Cf-9B- mediated resistance effective. With regards to the PVX experiment described above, the Cf-9B resistance pathway being activated in young tomato plants could then be explained by the largescale production and secretion of Avr9B at the site of the Cf-9B resistance protein in the plasma membrane, above and beyond what would normally be produced and secreted by *F. fulva* in the apoplast, thereby effectively bypassing the issue of stability/abundance. In accordance with this, Avr4 variants from *F. fulva* that naturally evade Cf-4-mediated resistance in tomato due to reduced stability in the leaf apoplast are still able to trigger a Cf-4-dependent HR in tomato plants carrying the *Cf-4* resistance gene using the PVX-based expression system (Joosten *et al*., 1997).

It could also be possible that *F. fulva* produces, for example, another effector that specifically suppresses Cf-9B-mediated resistance in young tomato plants. In line with this, suppression of resistance protein-mediated defence has been shown in other pathosystems. Examples include the AvrLm4-7 effector of the blackleg pathogen of *Brassica* crops, *Leptosphaeria maculans*, which suppresses Rlm3-mediated resistance triggered by the effector AvrLm3 (Plissonneau *et al*., 2016), as well as the Avr1 effector of the tomato wilt pathogen of tomato, *Fusarium oxysporum* f. sp. *lycopersici*, which suppresses I-1- and I-3-mediated resistance triggered by the effectors Avr1 and Avr3, respectively (Houterman *et al*., 2008). Yet another possibility could be that Avr9B is produced more abundantly in mature tomato plants, thereby exceeding the threshold required for Cf-9B-mediated resistance. However, the expression profile of *Avr9B* in mature tomato plants has not yet been investigated.

In any case, it may well be that, in young tomato plants, one or all of these phenomena has played a significant role in the sequential breakdown of the *Cf-9* resistance locus by *F. fulva.* This can be explained as follows: First, although *Cf-9B* and *Cf-9C* are stacked in *Cf-9* tomato cultivars, the instability/low abundance of Avr9B or the presence of a Cf-9B resistance-suppressing effector in the apoplastic leaf environment of young plants resulted in only Cf-9C being able to provide effective resistance. As such, there was significant selection pressure on *F. fulva* to overcome *Cf-9C*-mediated resistance through deletion of the corresponding *Avr9* effector gene. This gave *F. fulva* the ability to infect young *Cf-9* plants. Second, during the maturation of young, infected *Cf-9* plants, increasing selection pressure was exerted on *F. fulva* to overcome *Cf-9B*-mediated resistance through deletion or mutation of the corresponding *Avr9B* effector gene, concomitant with increased stability/higher abundance of Avr9B or the absence of a Cf-9B resistance-suppressing effector, subsequently giving *F. fulva* the ability to infect mature *Cf-9* plants.

Another factor that could be relevant to the sequential breakdown of the *Cf-9* resistance locus by *F. Fulva* is the stability of the genome regions carrying *Avr9* and *Avr9B*. For *Avr9*, the gene is known to be located in a gene-sparse, sub-telomeric, repeat-rich region of the strain Race 5 genome (Zaccaron *et al*., 2022) whereas, for *Avr9B*, the surrounding genome region is, for the most part, much more gene-rich. Reflective of a much more unstable genome environment, loss of *Avr9* through gene deletion may then occur more quickly than loss or mutation of *Avr9B*, enabling *Cf-9C*-mediated resistance to be overcome prior to *Cf-9B*- mediated resistance. In support of this, all *Cf-9C*-breaking strains of *F. fulva* identified to date lack the *Avr9* gene, with very few other mutations identified in this gene across non*-Cf-9C*- breaking strains (Bernal-Cabrera *et al*., 2021; Iida *et al*., 2015; Stergiopoulos *et al*., 2007a; Yoshida *et al*., 2021; van Kan *et al*., 1991).

Importantly, regardless of the hypotheses mentioned above, the sequential breakdown of the *Cf-9* locus is supported by our gene sequencing data, which showed that most *F. fulva* strains collected soon after the deployment of *Cf-9* cultivars lost *Avr9* but still carried a functional copy of *Avr9B*, whereas more recently collected strains not only lacked *Avr9*, but also possessed a resistance-breaking mutation in *Avr9B*. The relative ease with which both *Avr9* and *Avr9B* can be lost or mutated without an obvious impact on pathogen fitness (as shown for strain IPO 2679) suggests that neither gene plays a significant role in the virulence of *F. fulva* on tomato, or that their roles in virulence are redundant with other effector genes of this fungus. This possibility, though, would need to be examined further with *F. fulva* biomass quantification assays involving various *Avr9*/*Avr9B* deletion/complementation strains during infection of young and mature tomato plants.

Another interesting finding in our study was that Avr9B triggered chlorosis or, in some cases, weak cell death in *N. tabacum*, as well as strong necrosis in *N*. *benthamiana*, in the absence of Cf-9B. Likewise, although unable to trigger Cf-9B-dependent chlorosis or cell death, Avr9B-like proteins from the black leaf mould pathogen of tomato, *P. fuligena*, as well as the grey leaf spot pathogen of tomato, *S. lycopersici*, triggered Cf-9B-independent chlorosis or strong necrosis in *N. benthamiana* and *N. tabacum*. In *N. benthamiana*, the Cf-9B- independent responses triggered by Avr9B and the Avr9B-like proteins were found not to be dependent on the RLP co-receptor, SOBIR1, perhaps indicating that these effectors interact with an endogenous resistance protein that does not require SOBIR1 for transducing defence response signals, following apoplastic effector recognition (e.g. an RLK or a SOBIR1- independent RLP). It should be noted that Avr9B is not the first effector of *F. fulva* to trigger cell death in these non-host plants, with the candidate Avr effector Ecp2-1 also triggering cell death in several *Nicotiana* species (de Kock *et al*., 2004; Laugé *et al*., 2000). In this example, recognition is thought to be encoded by an endogenous resistance protein that is not homologous to Cf RLPs (de Kock *et al*., 2004). Another possibility could be that Avr9B and the abovementioned Avr9B-like proteins trigger Cf-9B-independent responses in different way, such as by interacting with and perturbing the plant plasma membrane, as has been shown for NLP effectors of various microbial pathogens (Pirc *et al*., 2022). Whether Avr9B and Avr9B-like proteins interact with the plasma membrane of plant cells remains to be determined, but in a possible connection to this ability, a subset of Avr9B-like proteins (Group 3 proteins) were predicted to have a transmembrane domain.

Intriguingly, expression of the Avr9B-like protein from *S. lycopersici* in tomato using the PVX-based expression system resulted in stunted growth regardless of the tomato genotype, which could indicate that it has a specific function in modulating host physiology. Alternatively, it could be that this protein is recognised by an endogenous resistance protein in tomato, but that this recognition does not result in an HR. This may be similar to a homolog of the Ecp2-1 protein from the black sigatoka pathogen, *Pseudocercospora fijiensis*, which triggers weak necrosis in tomato, independent of genotype (Stergiopoulos *et al*., 2010). In any case, the response triggered by the Avr9B-like protein from *S. lycopersici* was different to that exhibited in tomato plants expressing Avr9B, since the systemic expression of Avr9B did not result in stunting but, rather, exaggerated PVX symptoms. Such exaggerated PVX symptoms were not observed when any other Avr effector protein of *F. fulva* was systemically expressed in tomato, suggesting that this is a property unique to Avr9B. Whether these exaggerated PVX symptoms have resulted from, for example, suppression of the plant immune system by Avr9B, enabling PVX to more effectively colonise MM-Cf-0 tomato plants, remains to be determined.

While the predicted structural fold of the cysteine-rich region from Avr9B did not reveal any insights into the function of this protein, the plant phenotypes mentioned above, together with the apparent intrinsic ability of Avr9B and Avr9B-like proteins to trigger chlorosis or cell death in *Nicotiana* species, provide leads for future research into the virulence roles that these proteins might play during host colonization. Certainly, it will be interesting to determine whether Avr9B is directly or indirectly recognized by Cf-9B, since all mutations in *Avr9B* appear to abolish a functional protein. This may suggest that Avr9B is indirectly recognized by Cf-9B through a guarded host virulence target, since direct recognition is more likely to be overcome by non-synonymous mutations in residues that are located on the surface of the protein, in line with what has been shown for the direct recognition of Avr effector proteins from the flax rust pathogen, *Melampsora lini* (Dodds *et al*., 2006).

In conclusion, we have identified the *Avr9B* effector gene from *F. fulva*, corresponding to the *Cf-9B* resistance gene of tomato, and provided evidence to support the sequential breakdown of the *Cf-9* resistance locus through the evolution of *F. fulva* strains over time. Ultimately, due to the worldwide distribution of *F. fulva* strains capable of overcoming both *Cf-9B* and *Cf-9C*, the *Cf-9* locus, which is present in many commercially available tomato cultivars that are resistant to this fungus, likely now has limited value for controlling leaf mould disease. For breeders, the rate at which *F. fulva* can overcome single *Cf* resistance genes should be of concern and considered in future breeding programmes. Strategies such as the stacking of novel *Cf* resistance genes from wild tomato accessions into tomato cultivars could provide a means of durable resistance (Mesarich *et al*., 2018). However, as our study highlights, it is crucial to first understand the dynamic relationship between Avr effector proteins and their corresponding resistance proteins to ensure that stacked resistance genes are not sequentially overcome by pathogens. Finally, by identifying *Avr9B*, strains of *F. fulva* that have overcome *Cf-9B* can now be rapidly identified which, in turn, may inform tomato cultivar selection or deployment.

## Materials and Methods

### Fungal strains and plant material

A list of the *F. fulva* strains used in this study, including their location and year of isolation, where known, is shown in **Table S3**. All strains are currently stored at the Laboratory of Phytopathology, Wageningen University and Research, the Netherlands, The Johanna Westerdijk Institute, the Netherlands, or the Laboratory of Plant Pathology, Setsunan University, Japan. Tomato plants used in this study were the WT near-isogenic *S. lycopersicum* lines MM-Cf-0 (no *Cf* genes) and MM-Cf-9 (carrying the *Cf-9* resistance locus made up of *Cf-9A*, *Cf-9B*, *Cf-9C*, *Cf-9D*, and *Cf-9E*) (Tigchelaar, 1984), as well as transgenic lines of MM-Cf-0 carrying *Cf-9C* alone, or both *Cf-9A* and *Cf-9B* (2/9-75) (Parniske *et al*., 1997). Tobacco plants used in this study were WT and Δ*sobir1 N. benthamiana* (Huang *et al*., 2021), as well as WT *N. tabacum* cultivar Wisconsin 38.

### Genome sequencing and assembly

*F. fulva* strain IPO 2679 was cultured in potato dextrose broth (PDB) in the dark at 22°C with gentle orbital shaking for two weeks. High-quality genomic DNA was extracted according to the method described by Schwessinger and McDonald (2017). A TruSeq™ Nano library was prepared and sequenced on the Illumina MiSeq™ (PE150) platform by Novagene (https://www.novogene.com). Fastp v.0.20.0 (Chen *et al*., 2017) was used to remove all low-quality bases from the sequencing reads. A *de novo* genome sequence was then assembled using SPAdes v3.11.1 (Bankevich *et al*., 2012), with the final assembly generated from a set of different kmers (21,33,55,77,99,127). The final genome assembly was assessed for quality using QUAST v5.0.2 (Gurevich *et al*., 2013) and searched for potential adapter (Illumina oligonucleotide sequence) contamination at the NCBI UniVec database using BLASTn.

### Bioinformatic analyses for the identification of candidate *Avr9B* genes

Alignments of *in planta*-expressed (candidate) effector genes and their encoding protein sequences from the *F. fulva* reference strain 0WU and strain IPO 2679 were generated using Geneious v.9.1.8 software (Kearse *et al*., 2012). These (candidate) effectors were previously identified by Mesarich *et al*. (2018) or Mesarich *et al*. (2014). N-terminal signal peptide predictions were carried out with SignalP v4.1 (Nielsen, 2017), while transmembrane domain, GPI anchor and IDR predictions were performed using TMHMM v2.0 (Krogh *et al*., 2001), Big-PI (Eisenhaber *et al*., 1999), and PONDR VLXT (Romero *et al*., 2001), respectively. BLASTn, tBLASTn and BLASTp were used to identify *Avr9B*-like genes and Avr9B-like proteins in NCBI and JGI sequence databases. Preference was given to cysteine number and spacing rather than a low E-value score. Gene exon–intron boundaries were predicted using expression data available at JGI and then mapped onto genes for which expression data were not available.

### PCR screens and analysis of allelic variation

Genomic DNA was extracted from *F. fulva* strains according to the protocol of van Kan *et al*. (1991), with the presence or absence of *Avr9* and *Avr9B* across these strains subsequently investigated using a PCR experiment in conjunction with the Avr9-F/Avr9-R and Avr9B- F/Avr9B-R primer pairs, respectively (**Table S4**). For this purpose, Phusion High-Fidelity DNA Polymerase (New England Biolabs), Phire Plant Direct PCR Master Mix (Thermo Fisher Scientific) or KOD One PCR Master Mix (TOYOBO) was used, with PCRs carried out as per the supplier’s instructions. PCR amplicons were then resolved on a 1% agarose gel by electrophoresis. Finally, to assess allelic variation in *Avr9* and *Avr9B* across *F. fulva* strains, PCR amplicons were purified using an E.Z.N.A. Gel Extraction Kit (Omega Bio-Tek) or the ExoSAP-IT PCR Cleanup Reagent (Applied Biosystems) and then directly sequenced using Sanger sequencing by the Massey Genome Service, Eurofins Scientific or Macrogen Japan with the same gene-specific forward and/or reverse primer employed for PCR amplification. To confirm deletion of the *Avr9B* gene, the region spanning the deleted region in strain IPO 2679 was determined through a nucleotide sequence alignment to the genome of isolate Race 5 (Zaccaron *et al*., 2022) using Geneious v9.1.8 software (Kearse *et al*., 2012). A PCR experiment was then carried out using the Avr9B_deletion-F_1/Avr9B_deletion-R_1 and Avr9B_deletion-F_2/Avr9B_deletion-R_2 primer pairs (**Table S4**), which flank the deleted region, in conjunction with GoTaq G2 DNA Polymerase (Promega), as per the supplier’s instructions.

### RT-qPCR analysis of gene expression

*F. fulva* strain Race 5 (carrying a functional copy of *Avr9B*) was inoculated onto young (4- week-old) *S. lycopersicum* MM-Cf-0 plants and the fourth composite leaf was harvested at 2, 4, 8, 12 and 16 dpi. *F. fulva* spore preparation and plant inoculations were identical to those described by Mesarich *et al*. (2014), with subsequent growth of inoculated plants carried out in a greenhouse without climate control under natural light conditions. *F. fulva* Race 5 was also cultured in PDB, and the mycelia were harvested at four dpi. In all cases, samples were harvested in triplicate (i.e. from independent plants or cultures to give three biological replicates) and flash-frozen in liquid nitrogen. Total RNA was extracted from 100 mg of ground leaf or mycelial material using 1 ml TRIzol (Thermo Fisher Scientific) and purified with an RNeasy Mini kit (Qiagen). cDNA was then synthesized from 5 µg of total RNA, using the QuantiTect Reverse Transcription Kit (Qiagen). Genomic DNA was eliminated through use of the gDNA Wipeout Buffer contained in the same kit and confirmed through PCR using the Avr9B-F/Avr9B-R primer pair (**Table S4**). RT-qPCR experiments were performed on cDNA samples according to Mesarich *et al*. (2014) using the qCfActin-F/qCfActin-R and qAvr9B-F/qAvr9B-R primer pairs (**Table S4**), which were designed with Primer3 (Untergasser *et al*. 2012). The efficiency and specificity of these primers were determined using a dilution series of cDNA for the *Avr9B* candidate genes, and genomic DNA for the *F. fulva actin* gene before use. The *actin* gene was used as a reference for normalization of gene expression as per Mesarich *et al*. (2014), and results were analysed according to the 2^-ΔCt^ method described by Livak and Schmittgen (2001). The results are the average of three biological replicates.

### *A. tumefaciens*-mediated transient transformation assays (ATTAs)

Candidate *Avr9B* genes *Ecp5* and *CfCE54*, a variant of *CfCE54* without its repeat region, as well as *Avr9B-*like genes from *S. lycopersici* and *P. fuligena*, each behind the nucleotide sequence encoding the PR1a signal peptide for secretion to the leaf apoplast and a 3xFLAG tag for detection by Western blotting, were synthesized in the ATTA expression vector, pICH86988 (Weber *et al*., 2011), by Twist Bioscience. To generate a variant of *CfCE54* without the nucleotide sequence encoding the PR1a signal peptide, the corresponding coding sequence was amplified from the synthesized version of *CfCE54* using the NSPCfCE54_BsaI-F/CfCE54_BsaI-R primer pair (**Table S4**). Likewise, to create a variant of *CfCE54* with a codon conferring the W97C substitution, the corresponding codon sequence was amplified from the synthesized version of *CfCE54*, using the CfCE54_BsaI-F/W97C-R and W97C- F/CfCE54_BsaI-R primer pairs (**Table S4**). In each case, the primers introduced *Bsa*I restriction sites, which enabled their subsequent assembly into pICH86988, using the Golden Gate cloning system (Engler *et al*., 2008). Sequence-verified ATTA expression vectors were transformed into *A. tumefaciens* GV3101 (Holsters *et al*., 1980), while *A. tumefaciens* strains carrying pCBJ10-based binary ATTA expression vectors with *Cf-9C* or *Cf-9B* (Chakrabarti *et al*., 2009) were kindly provided by David Jones (The Australian National University). The *Avr9* ATTA expression construct was generated by Honée *et al*. (1998). ATTAs were performed in leaves of *N. benthamiana* or *N. tabacum* according to Guo *et al*. (2020), with a final *A. tumefaciens* OD_600_ of 0.5 used. Symptoms were assessed at 7 days post-infiltration.

### Protein immunoblotting

*N. tabacum* leaves used for ATTAs were flash-frozen in liquid nitrogen at 2 days post-agroinfiltration. Samples were ground to a fine powder in liquid nitrogen, resuspended in GTEN buffer (10% v/v glycerol, 100 mM Tris pH 7.5, 1 mM EDTA, 150 mM NaCl, 10 mM DTT, 0.2% IGEPAL CA360, 10 µl/ml protease inhibitor cocktail (Sigma-Aldrich), 1% w/v PVP), incubated at 4°C with gentle orbital shaking for 30 min, and then centrifuged at 4°C and 5,000 *g* for 20 min. The supernatant was collected and transferred to a 1.5 mL microtube through Miracloth (Millipore). Protein samples were loaded onto a 12% Tris-glycine gel, transferred to a PVDF membrane (Bio-Rad) and probed with mouse anti-FLAG (1:5,000) primary antibody (Sigma-Aldrich) and chicken anti-mouse IgG-HRP (1:20,000) secondary antibody (Santa Cruz Biotechnology). Blotted membranes were incubated with SuperSignal West Pico chemiluminescent substrate (Thermo Fisher Scientific) and visualized with a C600 Gel Imaging System (Azure Biosystems).

### Gene complementation and virulence assays

Genomic DNA from *F. fulva* strain 0WU was extracted as per van Kan *et al*. (1991). The open reading frames of *Ecp5* and *CfCE54*, including approximately 1 kb from each 5′ (promoter) and 3′ (terminator) untranslated region, were amplified by PCR using the primer pairs GibEcp5-F/GibEcp5-R and GibCfCE54-F/GibCfCE54-R, which contained an additional 18–25 bp at the 5′ end homologous to the pFBTS1 backbone. Amplicons were extracted and ligated into pFBTS1 (a modified version of pFBT004) (Bolton *et al*., 2008) using a Gibson Assembly approach (Gibson *et al*., 2009). Amplification of the pFBTS1 vector backbone by PCR was performed with the primer pairs pFBTS1-F/pFBTS1BB-R and pFBTS1BB-F/pFBTS1-R. Sequence-verified complementation constructs, as well as the empty vector pFBTS1, were transformed into *A. tumefaciens* AGL1 and then introduced into WT strain IPO 2679 (*Avr9*^−^/*Avr9B*^−^) as per Ökmen *et al*. (2013). Genomic DNA was extracted from transformants and complementation confirmed with the Comp-F/CompEcp5-R and Comp-F/CompCfCE54-R primer pairs for Ecp5 and CfCE54, respectively (**Table S4**). In addition, complementation with a single-copy gene of either the *Ecp5* or *CfCE54* was confirmed by quantitative PCR using the qAvr9B-F/qAvr9B-R and qECP5-F/qECP5-R primer pairs (**Table S4**) in conjunction with the formula: Ratio = [(E_target_)^ΔCt target (control-sample)^]/[(E_reference_)^ΔCt reference (control-sample)^]. Here, the F. *fulva actin* gene was used as a reference single-copy-gene in conjunction with the qCfActin-F/qCfActin-R primer pair (**Table S4**). For virulence assays, conidia from nine independent transformants, representing pFBTS1 EV (two), pFBTS1::*Ecp5* (two), and pFBTS1::C*fCE54* (five), as well as the WT strains IPO 2679 (*Avr9*^−^/*Avr9B*^−^; control), ICMP 7320 (*Avr9*^−^/*Avr9B*^+^; control), P13 (*Avr9*^+^/*Avr9B*^+^; control), P18 (*Δavr9*/*Avr9B^W97C^*; natural variant) and P31 (*Δavr9*/*Avr9B^C107*^*; natural variant), were inoculated onto the four compound leaves immediately below the first flowering truss of mature (13-week-old) MM-Cf-0 and MM-Cf-9 tomato plants. For this purpose, conidia preparation, inoculation, and growth conditions were as described by Mesarich *et al*. (2014), with two exceptions. Firstly, prior to inoculation, tomato plants were maintained in high humidity conditions (approx. 100% RH), with the main shoot above the first flowering truss, including the apical meristem, removed after appearance of the first inflorescence/flowering truss. Secondly, the lateral shoots of tomato plants were instead periodically removed throughout the experiment. The level of disease severity was assessed at 23 dpi.

### PVX-mediated systemic transient expression assays

PVX-mediated systemic transient expression of Avr2, Avr4, Avr4E, Avr5, Avr9, Avr9B, Ecp5 and Ecp11-1, as well as TW65_01570 and pSfinx alone (EV) in MM-Cf-0 and MM-Cf-9 plants, transgenic MM-Cf-0 lines expressing *Cf-9C* or both *Cf-9A* and *Cf-9B* (*Cf-9A* + *Cf-9B*), and *S. pimpinellifolium* accession CGN14353, was performed as per Mesarich *et al*. (2018). In short, both cotyledons of 10-day-old tomato seedlings were infiltrated on the abaxial side with the appropriate *A. tumefaciens* culture, with symptoms assessed 10–18 dpi. The pSfinx::EV, pSfinx::*Avr2*, pSfinx::*Avr4*, pSfinx::*Avr4E*, pSfinx::*Avr5*, pSfinx::*Avr9*, pSfinx::*Ecp5*, and pSfinx::*Ecp11-1* expression constructs were generated in previous studies (Hammond-Kosack *et al*., 1995; Laugé *et al*., 2000; Luderer *et al*., 2002b; Mesarich *et al*., 2014; Mesarich *et al*., 2018; Joosten *et al*., 1997; Westerink *et al*., 2004).

### Prediction of Avr9B tertiary structure

The tertiary structure of the cysteine-rich region from Avr9B was predicted using AlphaFold2 (Jumper *et al*., 2021), in conjunction with ColabFold (Mirdita *et al*., 2021), and was visualized using PyMOL (DeLano 2002). The tertiary structure prediction was assisted through use of a custom multiple sequence alignment, which was generated using Geneious v9.1.8 (Kearse *et al*., 2012), and was based on the cysteine-rich region of Avr9B and all Avr9B-like proteins identified from other fungi (**Table S3**). The Foldseek server (van Kempen *et al*., 2023) was used to identify possible structural homologs of the cysteine-rich region from Avr9B in the RCSB Protein Data Bank.

## Acknowledgements

We thank Matt Templeton (The New Zealand Institute for Plant and Food Research) for critically reviewing the manuscript and David Jones (The Australian National University) for providing the *A. tumefaciens* strains carrying the *Cf-9B* and *Cf-9C* expression constructs. We acknowledge Marie Turner and Claudie Monot of the VEGENOV institute (https://www.vegenov.com/), Phillip Nicot of the National Research Institute for Agriculture, Food and the Environment in Montfavet, as well as Yubo Liu and Roelf Schreuder from Gourmet Mokai Ltd for providing *F. fulva* strains, and both the HARNESSTOM project (HARNESSTOM Grant Agreement N°: 101000716) and JSPS KAKENHI (20H02993) (YI) for partially funding the research. SdlR was supported by a Massey University Doctoral Scholarship.

## Author Contributions

CHM, SdlR, CRS, MHAJJ, YI, JKB, REB, PJGMdW and YB designed the research; SdlR, CRS, ÁRP, AMH, KM, YI, MT, RJ, HGB and MR performed the research; SdlR, CRS, DJW, and ÁRP carried out the data analysis, collection, or interpretation; SdlR, CHM and CRS wrote the manuscript. All authors reviewed the manuscript and approved it for publication.

## Data Availability

The assembled genome sequence of *F. fulva* strain IPO 2679 has been deposited under NCBI BioProject ID PRJNA994185, BioSample accession SAMN36418479, and genome accession JAUKVN000000000.

## References

Bankevich A, Nurk S, Antipov D, Gurevich AA, Dvorkin M, Kulikov AS, Lesin VM, Nikolenko SI, Pham S, Prjibelski AD et al. 2012. SPAdes: a new genome assembly algorithm and its applications to single-cell sequencing. Journal of Computational Biology 19: 455–477.

Bernal-Cabrera A, Martínez-Coca B, Herrera-Isla L, Ynfante-Martínez D, Peteira-Delgado B, Portal O, Leiva-Mora M, de Wit PJGM. 2021. The first report on the occurrence race 9 of the tomato leaf mold pathogen *Cladosporium fulvum* (syn. Passalora fulva) in Cuba. European Journal of Plant Pathology 160: 731–736.

Bolton MD, van Esse HP, Vossen JH, de Jonge R, Stergiopoulos I, Stulemeijer IJE, van den Burg GCM, Borrás-Hidalgo O, Dekker HL, de Koster CG, et al. 2008. The novel *Cladosporium fulvum* lysin motif effector Ecp6 is a virulence factor with orthologues in other fungal species. Molecular Microbiology 69: 119–136.

Chakrabarti A, Panter SN, Harrison K, Jones JDG, Jones DA. 2009. Regions of the Cf-9B disease resistance protein able to cause spontaneous necrosis in *Nicotiana benthamiana* lie within the region controlling pathogen recognition in tomato. Molecular Plant–Microbe Interactions 22: 1214–1226.

Chen S, Huang T, Zhou Y, Han Y, Xu M, Gu J. 2017. AfterQC: automatic filtering, trimming, error removing and quality control for fastq data. BMC Bioinformatics 18: 91–100.

de Kock MJD, Iskandar HM, Brandwagt BF, Laugé R, de Wit PJGM, Lindhout P. 2004. Recognition of *Cladosporium fulvum* Ecp2 elicitor by non-host *Nicotiana* spp. is mediated by a single dominant gene that is not homologous to known *Cf*-genes. Molecular Plant Pathology 5: 397–408.

DeLano WL. 2002. Pymol: An open-source molecular graphics tool. CCP4 Newsletter on Protein Crystallography 40: 82–92.

de Wit PJGM. 2016. *Cladosporium fulvum* effectors: weapons in the arms race with tomato. Annual Review of Phytopathology 54: 1–23.

de Wit PJGM, van der Burgt A, Ökmen B, Stergiopoulos I, Abd-Elsalam KA, Aerts AL, Bahkali AH, Beenen HG, Chettri P, Cox MP et al. 2012. The genomes of the fungal plant pathogens *Cladosporium fulvum* and *Dothistroma septosporum* reveal adaptation to different hosts and lifestyles but also signatures of common ancestry. PLoS Genetics 8: e1003088.

de Wit PJGM, Joosten MHAJ, Thomma BHPJ, Stergiopoulos I. 2009. Gene for gene models and beyond: the *Cladosporium fulvum*-tomato pathosystem. In: Deising, H.B. (eds) Plant Relationships. The Mycota, vol 5. Springer, Berlin, Heidelberg.

Dixon MS, Hatzixanthis K, Jones DA, Harrison K, Jones JDG. 1998. The tomato *Cf-5* disease resistance gene and six homologs show pronounced allelic variation in leucine-rich repeat copy number. The Plant Cell 10: 1915–1925.

Dixon MS, Jones DA, Keddie JS, Thomas CM, Harrison K, Jones JDG. 1996. The tomato *Cf-2* disease resistance locus comprises two functional genes encoding leucine-rich repeat proteins. Cell 84: 451–459.

Dodds PN, Lawrence GJ, Catanzariti AM, Teh T, Wang CI, Ayliffe MA, Kobe B, Ellis JG. 2006. Direct protein interaction underlies gene-for-gene specificity and coevolution of the flax resistance genes and flax rust avirulence genes. Proceedings of the National Academy of Sciences 103: 8888–8893.

Eisenhaber B, Bork P, Eisenhaber F. 1999. Prediction of potential GPI-modification sites in proprotein sequences. Journal of Molecular Biology 292: 741–758.

Engler C, Kandzia R, Marillonnet S. 2008. A one pot, one step, precision cloning method with high throughput capability. PloS One 3: e3647.

Enya J, Ikeda K, Takeuchi T, Horikoshi N, Higashi T, Sakai T, Iida Y, Nishi K, Kubota M. 2009. The first occurrence of leaf mold of tomato caused by races 4.9 and 4.9.11 of *Passalora fulva* (syn. *Fulvia fulva*) in Japan. Journal of General Plant Pathology 75: 76–79.

Franco MEE, López S, Medina R, Saparrat MCN, Balatti P. 2015. Draft genome sequence and gene annotation of *Stemphylium lycopersici* strain CIDEFI-216. Genome Announcements 3: e01069–15.

Gibson DG, Young L, Chuang R-Y, Venter JC, Hutchison III CA, Smith HO. 2009. Enzymatic assembly of DNA molecules up to several hundred kilobases. Nature Methods 6: 343–345.

Gout L, Kuhn ML, Vincenot L, Bernard-Samain S, Cattolico L, Barbetti M, Moreno-Rico O, Balesdent M-H, Rouxel T. 2007. Genome structure impacts molecular evolution at the *AvrLm1* avirulence locus of the plant pathogen *Leptosphaeria maculans*. Environmental Microbiology 9: 2978–2992.

Guo Y, Hunziker L, Mesarich CH, Chettri P, Dupont P-Y, Ganley RJ, McDougal RL, Barnes I, Bradshaw RE. 2020. DsEcp2-1 is a polymorphic effector that restricts growth of *Dothistroma septosporum* in pine. Fungal Genetics and Biology 135: 103300.

Gurevich A, Saveliev V, Vyahhi N, Tesler G. 2013. QUAST: quality assessment tool for genome assemblies. Bioinformatics 29: 1072–1075.

Gust AA, Felix G. 2014. Receptor like proteins associate with SOBIR1-type of adaptors to form bimolecular receptor kinases. Current Opinion in Plant Biology 21: 104–111.

Haanstra JPW, Meijer-Dekens F, Lauge R, Seetanah DC, Joosten MHAJ, de Wit PJGM, Lindhout P. 2000. Mapping strategy for resistance genes against *Cladosporium fulvum* on the short arm of chromosome 1 of tomato: *Cf-ECP5* near the *Hcr9* Milky Way cluster. Theoretical and Applied Genetics 101: 661–668.

Hammond-Kosack KE, Jones JDG. 1994. Incomplete dominance of tomato *Cf* genes for resistance to *Cladosporium fulvum*. Molecular Plant–Microbe Interactions 7: 58–58.

Hammond-Kosack KE, Staskawicz BJ, Jones JDG, Baulcombe DC. 1995. Functional expression of a fungal avirulence gene from a modified potato virus X genome. Molecular Plant–Microbe Interactions 8: 181–185.

Hammond-Kosack KE, Tang S, Harrison K, Jones JDG. 1998. The tomato *Cf-9* disease resistance gene functions in tobacco and potato to confer responsiveness to the fungal avirulence gene product Avr9. The Plant Cell 10: 1251–1266.

Holsters M, Silva B, van Vliet F, Genetello C, de Block M, Dhaese P, Depicker A, Inzé D, Engler G, Villarroel R et al. 1980. The functional organization of the nopaline *A. tumefaciens* plasmid pTiC58. Plasmid 3: 212–230.

Honée G, Buitink J, Jabs T, de Kloe J, Sijbolts F, Apotheker M, Weide R, Sijen T, Stuiver M, de Wit PJGM. 1998. Induction of defense-related responses in Cf9 tomato cells by the AVR9 elicitor peptide of *Cladosporium fulvum* is developmentally regulated. Plant Physiology 117: 809–820.

Houterman PM, Cornelissen BJ, Rep M. 2008. Suppression of plant resistance gene-based immunity by a fungal effector. PLoS Pathogens 4: e1000061.

Huang WRH, Schol C, Villanueva SL, Heidstra R, Joosten MHAJ. 2021. Knocking out *SOBIR1* in *Nicotiana benthamiana* abolishes functionality of transgenic receptor-like protein Cf-4. Plant Physiology 185: 290–294.

Iakovidis M, Soumpourou E, Anderson E, Etherington G, Yourstone S, Thomas C. 2020. Genes encoding recognition of the *Cladosporium fulvum* effector protein Ecp5 are encoded at several loci in the tomato genome. G3: Genes, Genomes, Genetics 10: 1753–1763.

Iida Y, Iwadate Y, Kubota M, Terami F. 2010. Occurrence of a new race 2.9 of leaf mold of tomato in Japan. Journal of General Plant Pathology 76: 84–86.

Iida Y, van’t Hof P, Beenen H, Mesarich C, Kubota M, Stergiopoulos I, Mehrabi R, Notsu A, Fujiwara K, Bahkali A, et al. 2015. Novel mutations detected in avirulence genes overcoming tomato *Cf* resistance genes in isolates of a Japanese population of *Cladosporium fulvum*. PLoS One 10: e0123271.

Jones DA, Dickinson MJ, Balint-Kurti PJ, Dixon MS, Jones JDG. 1993. Two complex resistance loci revealed in tomato by classical and RFLP mapping of the *Cf-2*, *Cf-4*, *Cf-5*, and *Cf-9* genes for resistance to *Cladosporium fulvum*. Molecular Plant–Microbe Interactions 6: 348–348.

Jones DA, Thomas CM, Hammond-Kosack KE, Balint-Kurti PJ, Jones JDG. 1994. Isolation of the tomato *Cf-9* gene for resistance to *Cladosporium fulvum* by transposon tagging. Science 266: 789–793.

Joosten MHAJ, Cozijnsen TJ, de Wit PJGM. 1994. Host resistance to a fungal tomato pathogen lost by a single base-pair change in an avirulence gene. Nature 367: 384–386.

Joosten MHAJ, Vogelsang R, Cozijnsen TJ, Verberne MC, de Wit PJGM. 1997. The biotrophic fungus *Cladosporium fulvum* circumvents *Cf-4*-mediated resistance by producing unstable AVR4 elicitors. The Plant Cell 9: 367–379.

Jumper J, Evans R, Pritzel A, Green T, Figurnov M, Ronneberger O, Tunyasuvunakool K, Bates R, Žídek A, Potapenko A, et al. 2021. Highly accurate protein structure prediction with AlphaFold. Nature 596: 583–589.

Kang WH, Yeom SI. 2018. Genome-wide identification, classification, and expression analysis of the receptor-like protein family in tomato. The Plant Pathology Journal. 34, 435– 444.

Kearse M, Moir R, Wilson A, Stones-Havas S, Cheung M, Sturrock S, Buxton S, Cooper A, Markowitz S, Duran C et al. 2012. Geneious Basic: an integrated and extendable desktop software platform for the organization and analysis of sequence data. Bioinformatics 28: 1647– 1649.

Krogh A, Larsson B, von Heijne G, Sonnhammer ELL. 2001. Predicting transmembrane protein topology with a hidden Markov model: application to complete genomes. Journal of Molecular Biology 305: 567–580.

Laterrot H. 1986. Race 2.5.9, a new race of *Cladosporium fulvum* (*Fulvia fulva*) and sources of resistance in tomato. Netherlands Journal of Plant Pathology 92: 305–307.

Laugé R, Dmitriev AP, Joosten MHAJ, de Wit PJGM. 1998. Additional resistance gene(s) against *Cladosporium fulvum* present on the *Cf-9* introgression segment are associated with strong PR protein accumulation. Molecular Plant–Microbe Interactions 11: 301–308.

Laugé R, Goodwin PH, de Wit, PJGM, Joosten MHAJ. 2000. Specific HR-associated recognition of secreted proteins from *Cladosporium fulvum* occurs in both host and non-host plants. The Plant Journal 23: 735–745.

Liebrand TW, van den Berg GC, Zhang Z, Smit P, Cordewener JH, America AH, Sklenar J, Jones AM, Tameling WI, Robatzek S, et al. 2013. Receptor-like kinase SOBIR1/EVR interacts with receptor-like proteins in plant immunity against fungal infection. Proceedings of the National Academy of Sciences 110: 10010–10015.

Liebrand TWH, van den Burg HA, Joosten MHAJ. 2014. Two for all: receptor-associated kinases SOBIR1 and BAK1. Trends in Plant Science 19: 123–132.

Lindhout P, Korta W, Cislik M, Vos I, Gerlagh T. 1989. Further identification of races of *Cladosporium fulvum* (*Fulvia fulva*) on tomato originating from the Netherlands, France and Poland. Netherlands Journal of Plant Pathology 95: 143–148.

Livak KJ, Schmittgen TD. 2001. Analysis of relative gene expression data using real-time quantitative PCR and the 2^−ΔΔ*C*T^ method. Methods 25: 402–408.

Luderer R, de Kock MJD, Dees RHL, de Wit PJGM, Joosten MHAJ. 2002a. Functional analysis of cysteine residues of ECP elicitor proteins of the fungal tomato pathogen *Cladosporium fulvum*. Molecular Plant Pathology 3: 91–95.

Luderer R, Takken FLW, de Wit PJGM, Joosten MHAJ. 2002b. *Cladosporium fulvum* overcomes *Cf-2*-mediated resistance by producing truncated AVR2 elicitor proteins. Molecular Microbiology 45: 875–884.

Mesarich CH, Barnes I, Bradley EL, de la Rosa S, de Wit PJGM, Guo Y, Griffiths SA, Hamelin RC, Joosten MHAJ, Lu M et al. 2023. Beyond the genomes of *Fulvia fulva* (syn. Cladosporium fulvum) and Dothistroma septosporum: new insights into how these fungal pathogens interact with their host plants. Molecular Plant Pathology 24: 474–494.

Mesarich CH, Bowen JK, Hamiaux C, Templeton MD. 2015. Repeat-containing protein effectors of plant-associated organisms. Frontiers in Plant Science 6: 872.

Mesarich CH, Griffiths SA, van der Burgt A, Ökmen B, Beenen HG, Etalo DW, Joosten MHAJ, de Wit PJGM. 2014. Transcriptome sequencing uncovers the *Avr5* avirulence gene of the tomato leaf mold pathogen *Cladosporium fulvum*. Molecular Plant–Microbe Interactions 27: 846–857.

Mesarich CH, Ӧkmen B, Rovenich H, Griffiths SA, Wang C, Karimi Jashni M, Mihajlovski A, Collemare J, Hunziker L, Deng CH, et al. 2018. Specific hypersensitive response–associated recognition of new apoplastic effectors from *Cladosporium fulvum* in wild tomato. Molecular Plant–Microbe Interactions 31: 145–162.

Mirdita M, Schütze K, Moriwaki Y, Heo L, Ovchinnikov S, Steinegger M. 2022. ColabFold: making protein folding accessible to all. Nature Methods 19: 679–682.

Nielsen H. 2017. Predicting secretory proteins with SignalP. In: Kihara D. (eds) Protein function prediction. Methods in Molecular Biology, vol. 1611. Humana Press, New York, NY, 59–73.

Ngou BPM, Wyler M, Schmid MW, Kadota Y, Shirasu K. 2023. Evolutionary trajectory of pattern recognition receptors in plants. bioRxiv 2023.07.04.547604.

Ökmen B, Etalo DW, Joosten MHAJ, Bouwmeester HJ, de Vos RCH, Collemare J, de Wit PJGM. 2013. Detoxification of α-tomatine by *Cladosporium fulvum* is required for full virulence on tomato. New Phytologist 198: 1203–1214.

Panter SN, Hammond-Kosack KE, Harrison K, Jones JDG, Jones DA. 2002. Developmental control of promoter activity is not responsible for mature onset of *Cf-9B*- mediated resistance to leaf mold in tomato. Molecular Plant–Microbe Interactions 15: 1099– 1107.

Parniske M, Hammond-Kosack KE, Golstein C, Thomas CM, Jones DA, Harrison K, Wulf BBH, Jones JDG. 1997. Novel disease resistance specificities result from sequence exchange between tandemly repeated genes at the *Cf-4*/*9* locus of tomato. Cell 91: 821–832.

Pirc K, Albert I, Nürnberger T, Anderluh G. 2023. Disruption of plant plasma membrane by Nep1-like proteins in pathogen–plant interactions. New Phytologist 237: 746–750.

Plissonneau C, Daverdin G, Ollivier B, Blaise F, Degrave A, Fudal I, Rouxel T, Balesdent MH. 2016. A game of hide and seek between avirulence genes *AvrLm4-7* and *AvrLm3* in *Leptosphaeria maculans*. New Phytologist 209: 1613–1624.

Postma J, Liebrand TWH, Bi G, Evrard A, Bye RR, Mbengue M, Kuhn H, Joosten MHAJ, Robatzek S. 2016. Avr4 promotes Cf-4 receptor-like protein association with the BAK1/SERK3 receptor-like kinase to initiate receptor endocytosis and plant immunity. New Phytologist 210: 627–642.

Rocafort M, Fudal I, Mesarich CH. 2020. Apoplastic effector proteins of plant-associated fungi and oomycetes. Current Opinion in Plant Biology 56: 9–19.

Romero P, Obradovic Z, Li X, Garner EC, Brown CJ, Dunker AK. 2001. Sequence complexity of disordered protein. Proteins: Structure, Function, and Bioinformatics 42: 38– 48.

Salcedo A, Rutter W, Wang S, Akhunova A, Bolus S, Chao S, Anderson N, De Soto MF, Rouse M, Szabo L et al. 2017. Variation in the *AvrSr35* gene determines *Sr35* resistance against wheat stem rust race Ug99. Science. 358: 1604–1606.

Schouten HJ, Tikunov Y, Verkerke W, Finkers R, Bovy A, Bai Y, Visser RG. 2019. Breeding has increased the diversity of cultivated tomato in the Netherlands. Frontiers in Plant Science 10: 1606.

Schwessinger B, McDonald MC. 2017. High quality DNA from Fungi for long read sequencing e.g. PacBio, Nanopore MinION V.4. Protocols.io 10.17504/protocols.io.k6qczdw.

Snoeck S, Garcia AG, Steinbrenner AD. 2023. Plant receptor-like proteins (RLPs): structural features enabling versatile immune recognition. Physiological and Molecular Plant Pathology, 102004.

Stergiopoulos I, de Kock MJD, Lindhout P, de Wit PJGM. 2007a. Allelic variation in the effector genes of the tomato pathogen *Cladosporium fulvum* reveals different modes of adaptive evolution. Molecular Plant–Microbe Interactions 20: 1271–1283.

Stergiopoulos I, Groenewald M, Staats M, Lindhout P, Crous PW, de Wit PJGM. 2007b. Mating-type genes and the genetic structure of a world-wide collection of the tomato pathogen *Cladosporium fulvum*. Fungal Genetics and Biology 44: 415–429.

Stergiopoulos I, van den Burg HA, Ökmen B, Beenen HG, van Liere S, Kema GHJ, de Wit PJGM. 2010. Tomato Cf resistance proteins mediate recognition of cognate homologous effectors from fungi pathogenic on dicots and monocots. Proceedings of the National Academy of Sciences 107: 7610–7615.

Takken FL, Luderer R, Gabriëls SH, Westerink N, Lu R, de Wit PJGM, Joosten MHAJ. 2000. A functional cloning strategy, based on a binary PVX-expression vector, to isolate HR- inducing cDNAs of plant pathogens. The Plant Journal 24: 275–283.

Takken FLW, Thomas CM, Joosten MHAJ, Golstein C, Westerink N, Hille J, Nijkamp JJ, de Wit PJGM, Jones JDG. 1999. A second gene at the tomato *Cf-4* locus confers resistance to *Cladosporium fulvum* through recognition of a novel avirulence determinant. The Plant Journal 20: 279–288.

Tarallo M, McDougal RL, Chen Z, Wang Y, Bradshaw RE, Mesarich CH. 2022. Characterization of two conserved cell death elicitor families from the Dothideomycete fungal pathogens *Dothistroma septosporum* and *Fulvia fulva* (syn. Cladosporium fulvum). Frontiers in Microbiology. 13, 964851.

Thomas CM, Jones DA, Parniske M, Harrison K, Balint-Kurti PJ, Hatzixanthis K, Jones JD. 1997. Characterization of the tomato *Cf-4* gene for resistance to *Cladosporium fulvum* identifies sequences that determine recognitional specificity in Cf-4 and Cf-9. The Plant Cell 9: 2209–2224.

Thomas CM, Dixon MS, Parniske M, Golstein C, Jones JD. 1998. Genetic and molecular analysis of tomato *Cf* genes for resistance to *Cladosporium fulvum*. Philosophical Transactions of the Royal Society of London. 353: 1413–1424

Thomma BPHJ, van Esse HP, Crous PW, de Wit PJGM. 2005. *Cladosporium fulvum* (syn. Passalora fulva), a highly specialized plant pathogen as a model for functional studies on plant pathogenic Mycosphaerellaceae. Molecular Plant Pathology 6: 379–393.

Tigchelaar EC. 1984. Collections of isogenic tomato stocks. Report of the Tomato Genetics Cooperative 34: 55–57.

Untergasser A, Cutcutache I, Koressaar T, Ye J, Faircloth B, Remm M, Rozen S. 2012. Primer3-new capabilities and interfaces. Nucleic Acids Research 40: e115.

van der Beek JG, Verkerk R, Zabel P, Lindhout P. 1992. Mapping strategy for resistance genes in tomato based on RFLPs between cultivars: *Cf9* (resistance to *Cladosporium fulvum*) on chromosome 1. Theoretical and Applied Genetics 84: 106–112.

van den Ackerveken GFJM, van Kan JAL, de Wit PJGM. 1992. Molecular analysis of the avirulence gene *avr9* of the fungal tomato pathogen *Cladosporium fulvum* fully supports the gene-for-gene hypothesis. The Plant Journal 2: 359–366.

van der burgh AM, Postma J, Robatzek S, Joosten MHAJ. 2019. Kinase activity of SOBIR1 and BAK1 is required for immune signalling. Molecular Plant Pathology 20: 410– 422.

van Kan JAL, van den Ackerveken GFJM, de Wit PJGM. 1991. Cloning and characterization of cDNA of avirulence gene *avr9* of the fungal pathogen *Cladosporium fulvum*, causal agent of tomato leaf mold. Molecular Plant–Microbe Interactions 4: 52–59.

van Kempen M, Kim SS, Tumescheit C, Mirdita M, Lee J, Gilchrist CL, Söding J, Steinegger M. 2023. Fast and accurate protein structure search with Foldseek. Nature Biotechnology 10.1038/s41587-023-01773-0.

Videira SIR, Groenewald JZ, Nakashima C, Braun U, Barreto RW, de Wit PJGM, Crous PW. 2017. Mycosphaerellaceae–chaos or clarity? Studies in Mycology 87: 257–421.

Weber E, Engler C, Gruetzner R, Werner S, Marillonnet S. 2011. A modular cloning system for standardized assembly of multigene constructs. PLoS One 6: e16765.

Westerink N, Brandwagt BF, de Wit PJGM, Joosten MHAJ. 2004. *Cladosporium fulvum* circumvents the second functional resistance gene homologue at the *Cf-4* locus (*Hcr9-4E*) by secretion of a stable avr4E isoform. Molecular Microbiology 54: 533–545.

Wicker T, Sabot F, Hua-Van A, Bennetzen JL, Capy P, Chalhoub B, Flavell A, Leroy P, Morgante M, Panaud O et al. 2007. A unified classification system for eukaryotic transposable elements. Nature Reviews Genetics 8: 973–982.

Yoshida K, Asano S, Sushida H, Iida Y. 2021. Occurrence of tomato leaf mold caused by novel race 2.4.9 of *Cladosporium fulvum* in Japan. Journal of General Plant Pathology 87: 35–38.

Zaccaron AZ, Chen LH, Samaras A, Stergiopoulos I. 2022. A chromosome-scale genome assembly of the tomato pathogen *Cladosporium fulvum* reveals a compartmentalized genome architecture and the presence of a dispensable chromosome. Microbial Genomics 8: 000819.

Zaccaron AZ, Stergiopoulos I. 2020. First draft genome resource for the tomato black leaf mold pathogen *Pseudocercospora fuligena*. Molecular Plant–Microbe Interactions 33: 1441– 1445.

