## Supplementary material for "Sequential breakdown of the complex *Cf-9* leaf mould resistance locus in tomato by *Fulvia fulva*": Fig. S1 - S10

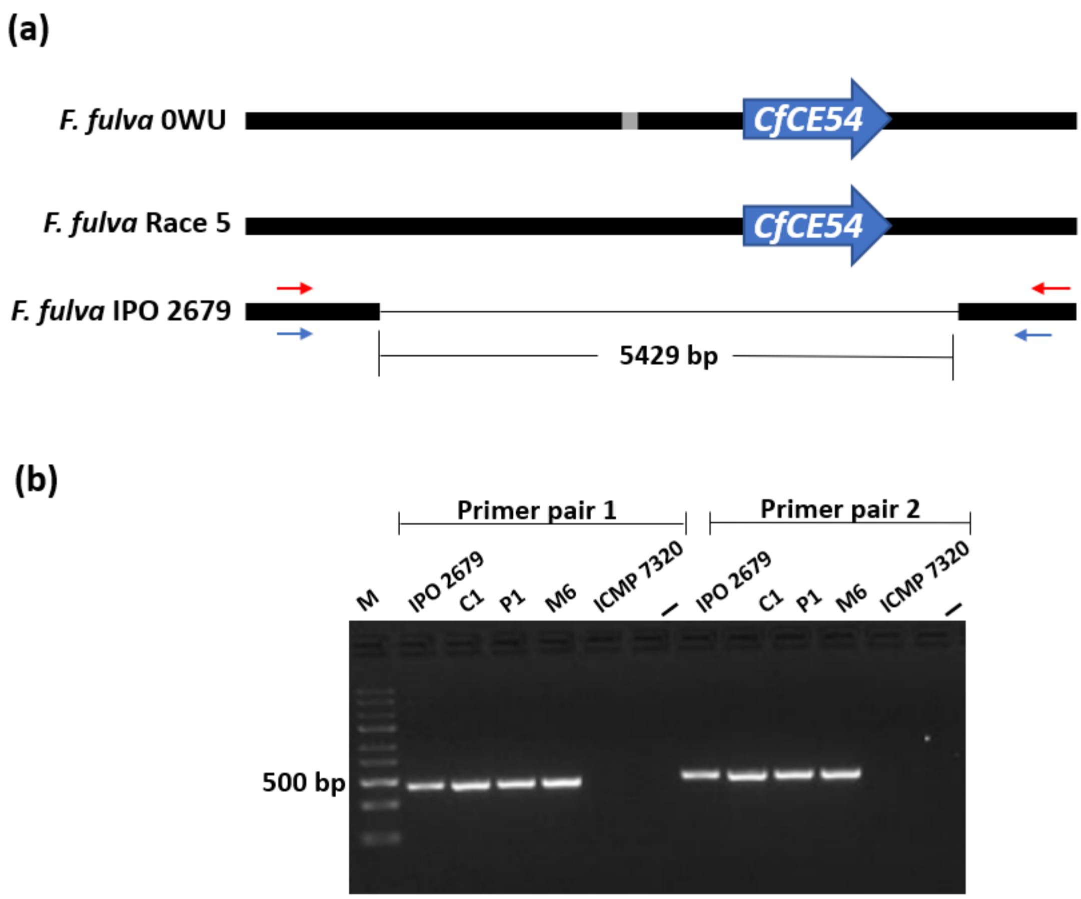


**Figure S1.** Characterization of the *CfCE54* genomic region in *Fulvia fulva* strains. **(a)** Alignment of the deleted *CfCE54* genomic region in *F. fulva* strain IPO 2679 ($Avr9B^{-}$) to the corresponding region from reference strains 0WU ($Avr9B^{+}$; de Wit *et al.*, 2012) and Race 5 ($Avr9B^{+}$; Zaccaron *et al.*, 2022). In strain 0WU, a region of Ns is shown in grey. In strain IPO 2679, the deleted region encompassing *Avr9B* is shown by a thin black line. **(b)** Confirmation of deletion for the *CfCE54* genomic region in New Zealand strains IPO 2679, C1, P1 and M8 by polymerase chain reaction (PCR) using two different primer pairs flanking the deleted region. Primer binding site locations are shown in (a), with blue representing primer pair 1 and red representing primer pair 2. Expected PCR amplicon sizes for deletion: primer pair 1 (in blue; 451 bp); primer pair 2 (in red; 486 bp). M, ladder marker; –, negative control (no DNA template). Primer locations are shown in (a). There are no PCR amplicons for strain ICMP 7320 ($Avr9B^{+}$), as the PCR extension time did not allow for amplification of a large genomic DNA region (5880 bp for primer pair 1 and 5915 bp for primer pair 2).

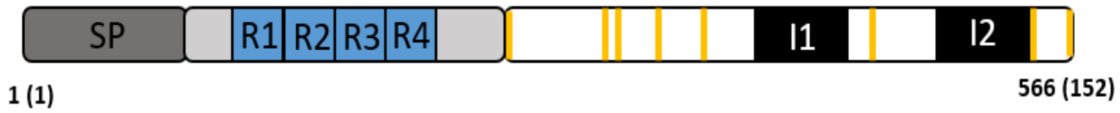


**Figure S2**. Schematic representation of the *CfCE54* gene from *Fulvia fulva.* Nucleotide sequence encoding the predicted signal peptide (SP) of CfCE54 is shown in dark grey. Nucleotide sequence encoding the predicted intrinsically disordered region of CfCE54 is shown in light grey and contains four imperfect tandem repeats (R1–R4; turquoise boxes). Nucleotide sequence encoding the cysteine-rich region of CfCE54 is shown in white and contains two introns (I1 and I2; black boxes). Nucleotide sequence encoding the eight cysteine residues of CfCE54 is shown by vertical yellow lines. The gene length (nucleotides) of *CfCE54* is shown below the schematic, and protein length (amino acids) is shown in brackets.

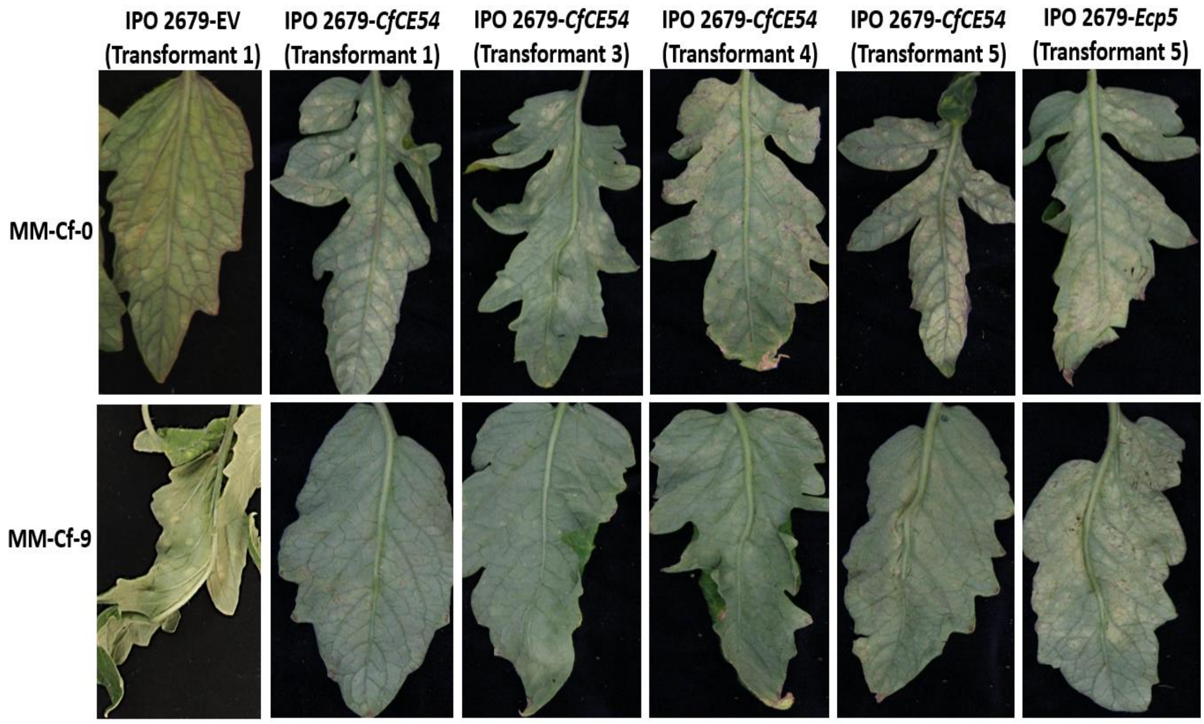


**Figure S3.** *CfCE54*, but not *Ecp5*, restores avirulence to strain IPO 2679 of *Fulvia fulva* on mature *Solanum lycopersicum* plants carrying the *Cf-9* resistance locus. *F. fulva* strains not included in Fig. 1 were inoculated onto mature (13-week-old) 'Moneymaker' (MM)-Cf-0 (no *Cf* genes) and MM-Cf-9 (carrying the *Cf-9* resistance locus) plants, with photographs taken at 23 days post-inoculation. IPO 2679 EV, strain IPO 2679 (*Avr9*⁻/*Avr9B*⁻) complemented with the pFBTS1 empty vector (no insert); IPO 2679-*CfCE54*, strain IPO 2679 complemented with the *CfCE54* gene from wild-type (WT) strain 0WU; IPO 2679-*Ecp5*, strain IPO 2679 complemented with the *Ecp5* gene from WT strain 0WU.

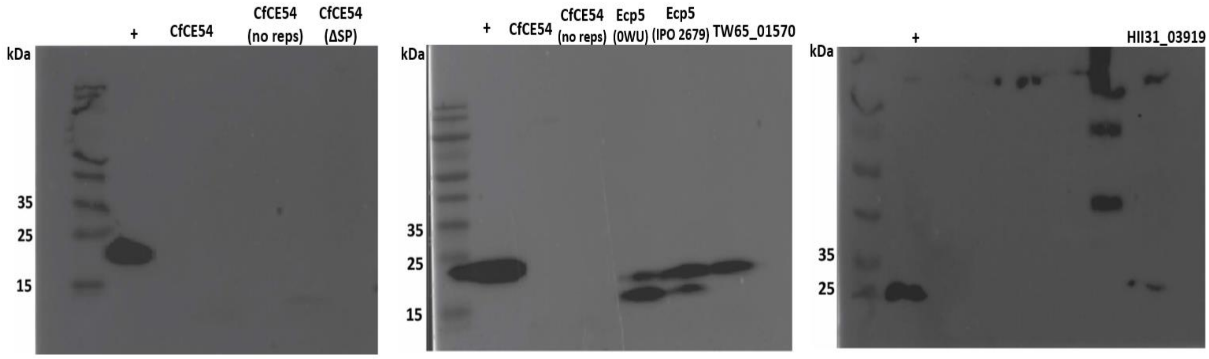


**Figure S4**. Detection of *Fulvia fulva*, *Stemphylium lycopersici* and *Pseudocercospora fuligena* secreted proteins by Western blotting following expression in leaves of *Nicotiana tabacum* using *Agrobacterium tumefaciens*-mediated transient transformation assays (ATTAs). Total protein was collected from leaf samples at 2 days post-agroinfiltration and the 3xFLAG tag was used for detection. Expected protein sizes: CfCE54 (Avr9B) from strain 0WU, 17 kDa; CfCE54 from strain 0WU with no repeat region (no reps), 11 kDa; CfCE54 from strain 0WU with no signal peptide (ΔSP), 16 kDa; Ecp5 from strain 0WU or IPO 2679, 20 kDa; TW65_01575 (Avr9B-like) from *S. lycopersici*, 20 kDa; HII31_03919 (Avr9B-like) from *P. fuligena*, 24 kDa. Positive control (+) is a *Dothistroma septosporum* candidate effector (25 kDa) from (Tarallo et al., 2022).

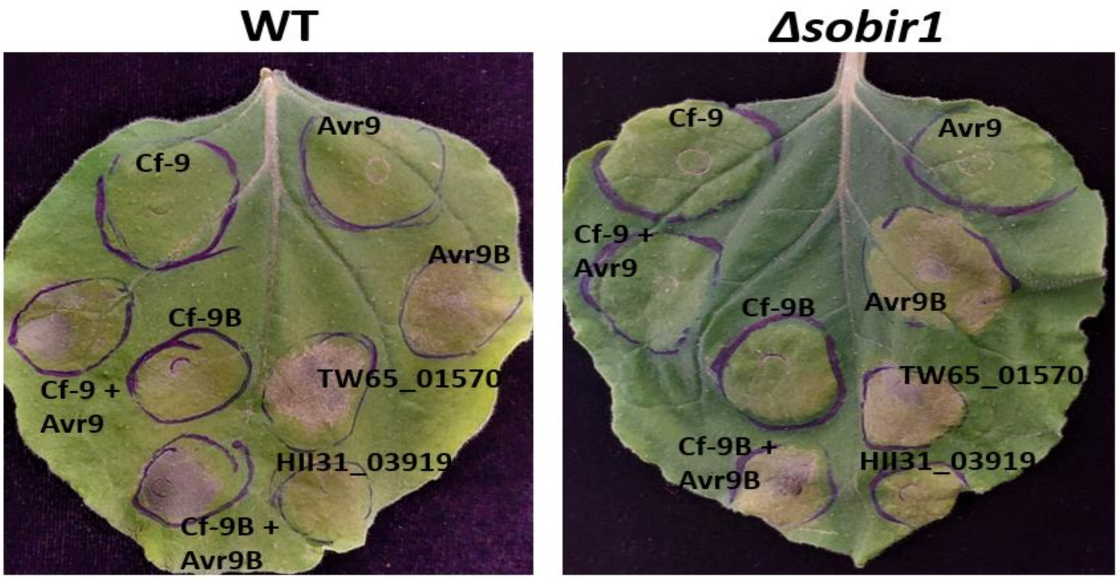


**Figure S5**. Avr9B and Avr9B-like proteins trigger chlorosis and/or cell death in *Nicotiana benthamiana* independent of SOBIR1. Avr9B from *Fulvia fulva*, as well as TW65_01570 (Avr9B-like) from *Stemphylium lycopersici* and HII31_03919 (Avr9B-like) from *Pseudocercospora fuligena* were expressed in leaves of wild-type (WT) and *Δsobir1 N. tabacum* using *Agrobacterium tumefaciens*-mediated transient transformation assays (ATTAs). The Cf-9 + Avr9 pair was used as a positive control in WT plants and a negative control in *Δsobir1* plants. Cf-9 and Avr9 alone were used as negative controls in both WT and *Δsobir1* plants. Leaves were photographed at 7 days post-infiltration and are representative of three independent ATTA experiments.

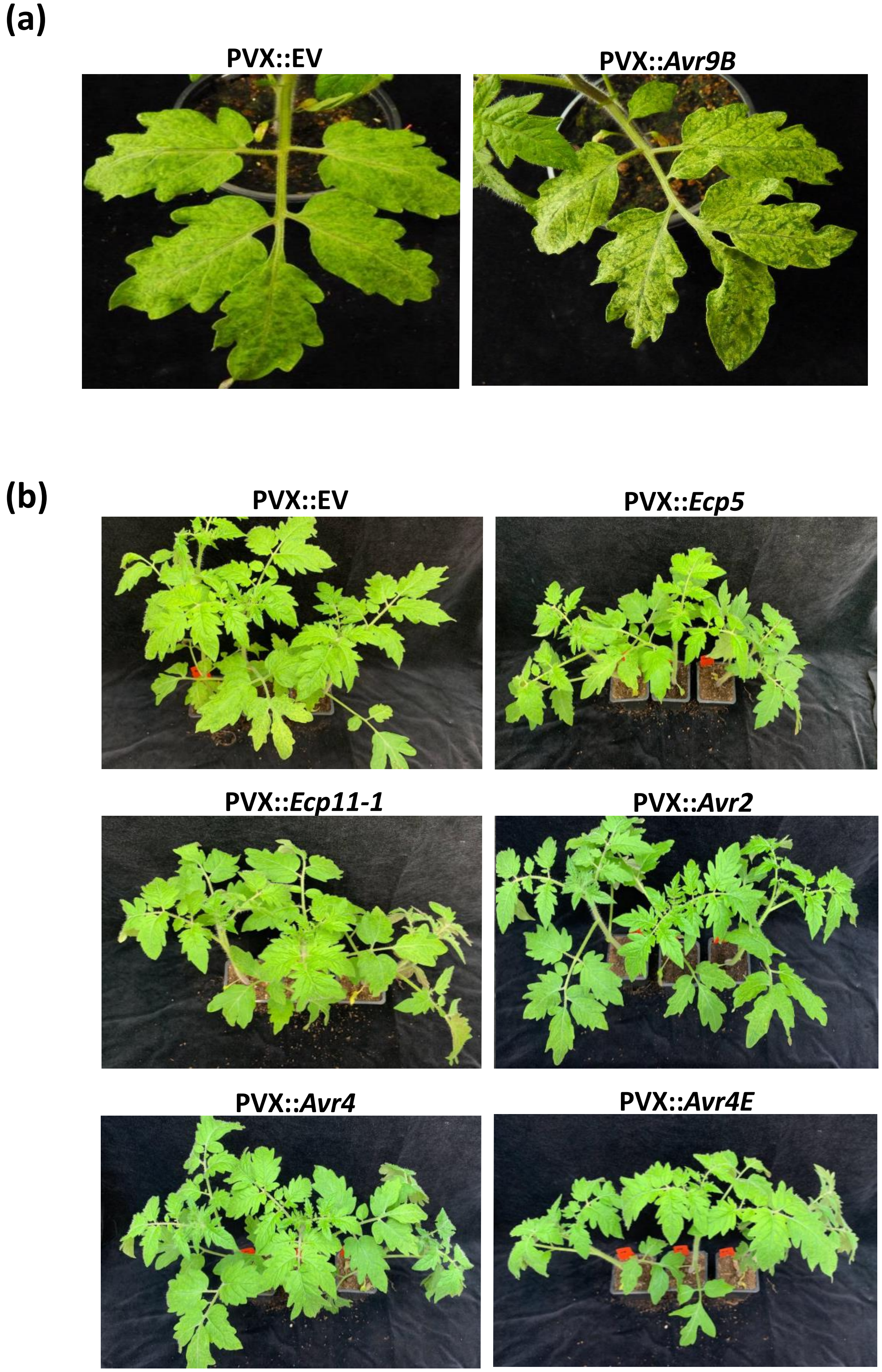
(a)
PVX::EV
PVX::*Avr9B*
(b)
PVX::EV
PVX::*Ecp5*
PVX::*Ecp11-1*
PVX::*Avr2*
PVX::*Avr4*
PVX::*Avr4E*

**PVX::*Avr5*** **PVX::*Avr9***

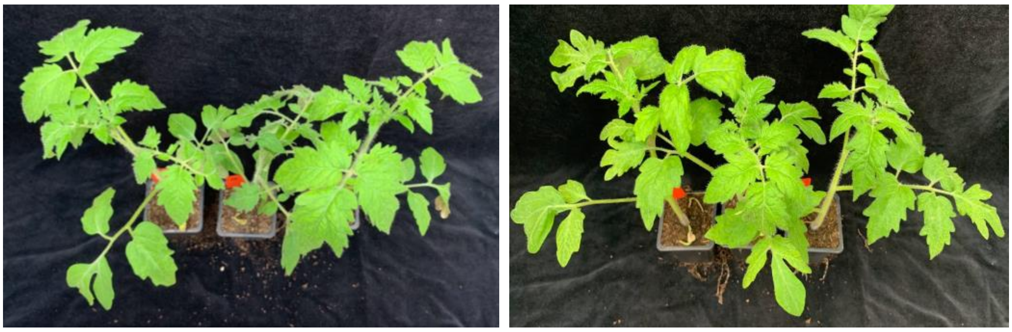

**Figure S6**. Avr9B from *Fulvia fulva* enhances Potato virus X (PVX) symptoms in 'Moneymaker' (MM)-Cf-0 tomato plants. *F. fulva* avirulence effectors Avr2, Avr4, Avr4E, Avr5, Avr9 and Avr9B, as well as candidate avirulence effectors Ecp5 and Ecp11-1, were systemically expressed in MM-Cf-0 (carrying no resistance genes) plants using the PVX-based expression system, with recombinant viruses PVX::*Avr2*, PVX::*Avr4*, PVX::*Avr4E*, PVX::*Avr5*, PVX::*Avr9*, PVX::*Avr9B*, PVX::*Ecp5* and PVX::*Ecp11-1*, as well as PVX::EV (pSfinx empty vector) delivered through cotyledon infiltration of 10-day-old tomato seedlings using *Agrobacterium tumefaciens*-mediated transient transformation. **(a)** Close-up showing that Avr9B enhances PVX symptoms in MM-Cf-0 plants, relative to the pSfinx EV control. **(b)** PVX symptoms in MM-Cf-0 plants associated with Avr2, Avr4, Avr4E, Avr5, Avr9, Ecp5 and Ecp11-1, relative to the Avr9B and the pSfinx EV control. Three plants were tested for each avirulence effector. Photographs were taken at 18 days post-infiltration.

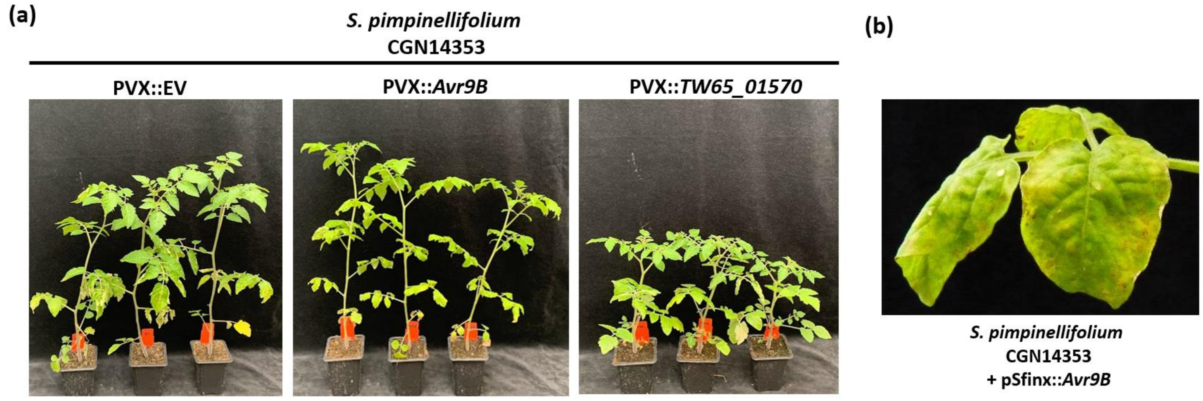


**Figure S7**. Avr9B from *Fulvia fulva* causes leaf curling, chlorosis, and weak cell death, whereas the Avr9B-like protein TW65_01570 from *Stemphylium lycopersici* causes stunting in the wild *Solanum pimpinellifolium* accession CGN14353. **(a)** Avr9B was systemically expressed in CGN14353 using the Potato virus X (PVX)-based expression system. Recombinant viruses PVX::*Avr9B*, PVX::*TW65_01759* and PVX::EV (pSfinx empty vector), were delivered through cotyledon infiltration of 10-day-old tomato seedlings using *Agrobacterium tumefaciens*-mediated transient transformation. For each recombinant virus, three plants were tested. Photographs were taken at 18 days post-infiltration. **(b)** Close-up of leaf curling, chlorosis and weak cell death caused by Avr9B in accession CGN14353.

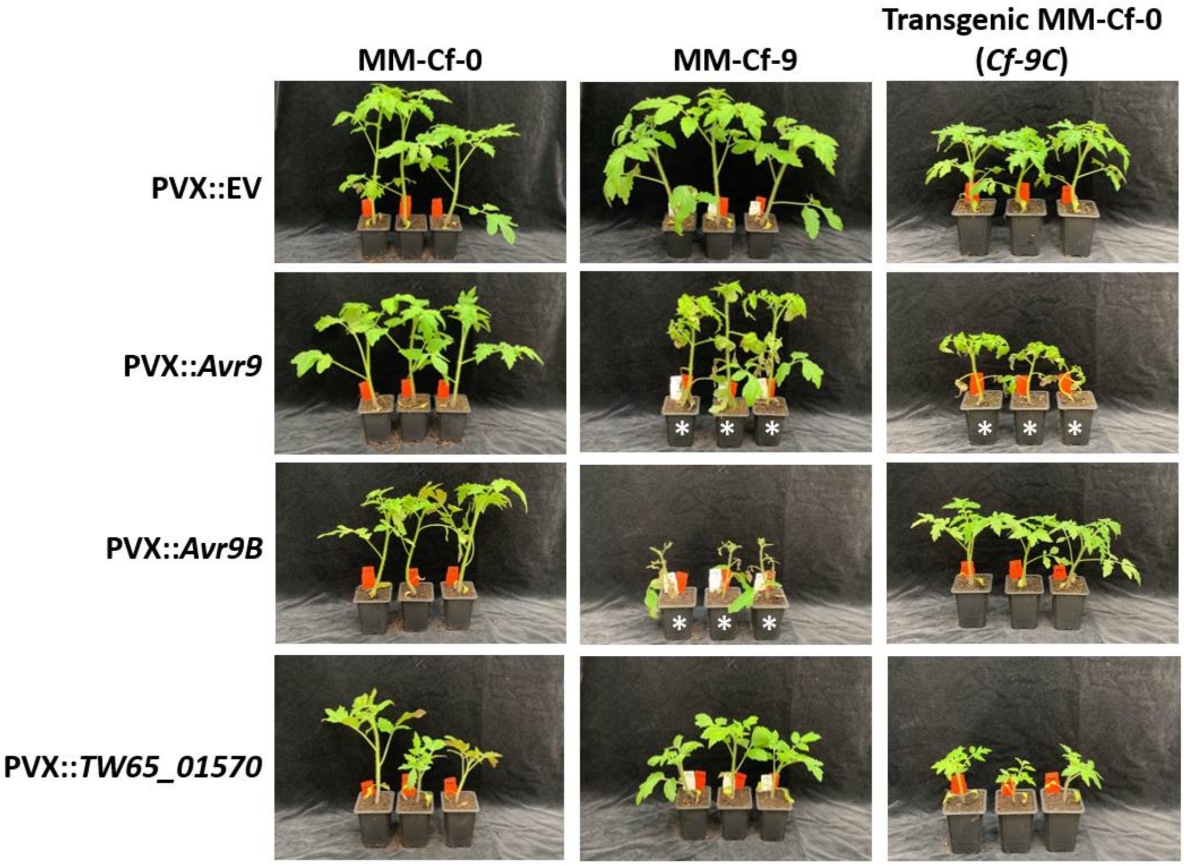


**Figure S8.** The Avr9B-like protein TW65_01570 from *Stemphylium lycopersici* causes stunting in young *Solanum lycopersicum* plants. TW65_01570 was systemically expressed in 'Moneymaker' (MM)-Cf-0 (carrying no resistance genes) and MM-Cf-9 (carrying the *Cf-9* resistance locus) plants, as well as transgenic MM-Cf-0 plants carrying the *Cf-9C* gene using the Potato virus X (PVX)-based expression system. Recombinant viruses PVX::*TW65_01759*, PVX::*Avr9B* and PVX::*Avr9*, encoded by pSfinx::*TW65_01570*, pSfinx::*Avr9* and pSfinx::*Avr9B*, respectively, as well as PVX::EV, encoded by pSfinx::EV (empty vector), were delivered through cotyledon infiltration of 10-day-old tomato seedlings using *Agrobacterium tumefaciens*-mediated transient transformation. Three representatives of each tomato accession were included in the experiment. White asterisks indicate plants undergoing a systemic hypersensitive response (HR). Photographs were taken at 18 days post-agroinfiltration.

**(a)** >*Avr9B*_strain_Tochigi1

ATGCGATCATCTATTGTCTCCCTCCTCGCCATCCAGCTGGCTTTCCTTGCCGTTGGTAACACGATGCCGA
CCGACCCTGCAGGATACACGGATCTATCAGCTCCAGAAATGGAAGGATCCACGGATCTATCAGCTCCAGA
TGTGGAAAAATCCACGGATCTATCAGCTCCAGATGTGGAAAAATCCACGGGTCTATCGGCTCCAGAAATG
AAAGAAGACGTCAACTCCTCTCCACAATCCTCTCTTTGTGAAAACCTTCATATCGAGGTACTGGGCCACC
GGCTCGGTTGGTGTGCGGGTTGTAAACGACTTGAGATTTGCGTCACCGTTCTCGCAGAGACTGGCGCTTC
ATGCGCTGCAGCCTATGTACAGTCCTGGGCATGGATTTTACAGCCCCTGTTCTTCCACCTTTCCTCATTA
GGAAGGATGTATCCACTGCTTGGCGTAGCTCACCACCTGGTGTGTCCTCCATTGTTGGCTATCACTGCCT
CGCAGTGTTCTCGCATACTCTTGCATATACGTTGTATGTCCTCAGGTGTGAGCTTGGCCCATTCTTGATA
GTGAGATTGAAGGTAGCAACACCTTCTGAGATGGACATGGATAGCCGGAAGGGGTTGGCGTAAATACAGG
GGTTGTAAAAACTATGCCCAGGACTGTACCAGGGGCTCAAGGTATGTTTGAATAACGAAGGCCTGGCTTG
GATGTGTATGATAACAGCAATAGCCGCACTCCTTGCCTGTTTGCTCTCAATCGGTGTCGATGAATATGAG
GCGTCAAAGTGTAGGTGTCCTACTCACTGTACATCACATGTCAAGCAAAGGCTGACTGGGATTAGGTACC
GATGGGTTGTGTTAG

**(b)**

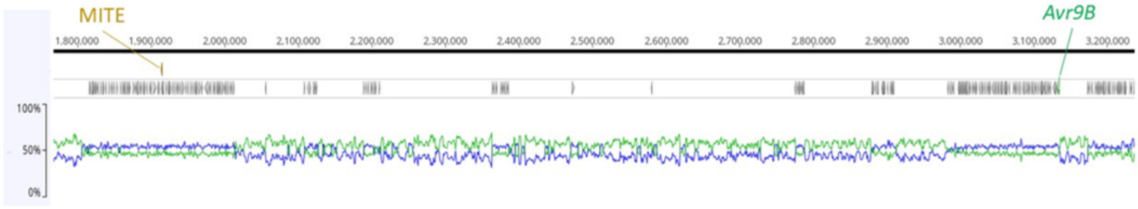


**Figure S9.** The *Avr9B* gene from strain Tochigi1 of *Fulvia fulva* in Japan is disrupted by a miniature inverted-repeat transposable element (MITE). **(a)** The MITE disrupts exon 1 of *Avr9B*. The MITE sequence is in red text. Target site duplication sequences and terminal inverted repeat sequences, both characteristic of MITEs, are highlighted green and cyan, respectively. Intron sequences are highlighted grey. **(b)** Location of the MITE on Chromosome 10 of the *F. fulva* strain Race 5 genome (Zaccaron *et al.*, 2022), relative to *Avr9B*. A zoomed-in region of Chromosome 10 is shown, with the MITE at position 1915484–1915772 coloured tan and the *Avr9B* gene at position 3134996–3134431 coloured green. All other predicted genes are shown in grey. In the genome of strain race 5, the MITE is annotated as "Motif1-Family73A". All other genes are shown in grey. G+C content is shown by a blue line, while A+T content is shown by a green line.

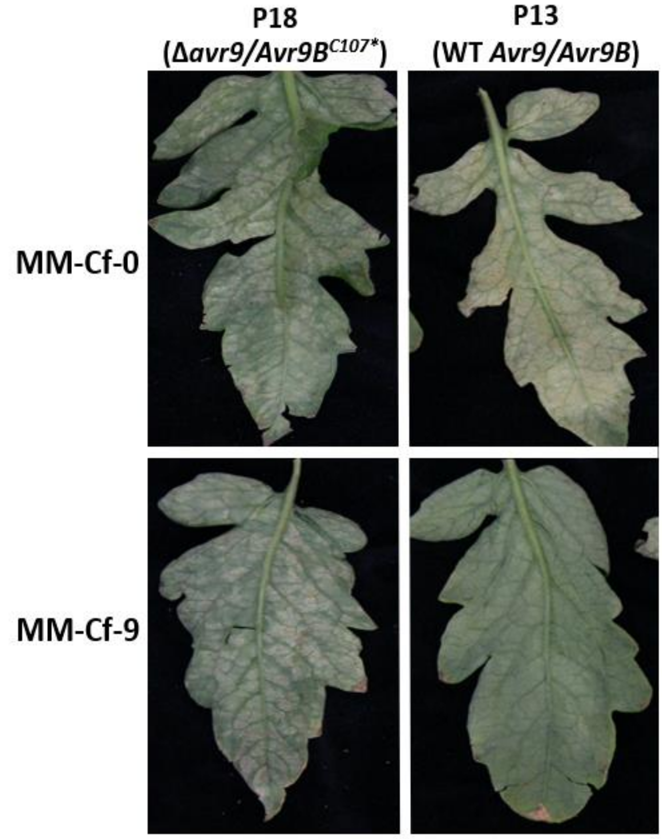


**Figure S10.** The natural *Avr9B*$^{C107*}$ mutant of *Fulvia fulva* is virulent on *Cf-9* plants. *F. fulva* strains P18 (Δ*avr9*/*Avr9B*$^{C107*}$) and P13 (wild-type (WT) *Avr9*/*Avr9B*) were inoculated onto mature (13-week-old) 'Moneymaker' (MM)-Cf-0 (no *Cf* genes) and MM-Cf-9 (carrying the *Cf-9* resistance locus) plants, with photographs taken at 23 days post-inoculation. Results are representative of three biological replicates and were performed at the same time as the pathogenicity assays shown in Fig. 1.
