## Supplementary Information 1 for "Sequential breakdown of the complex *Cf-9* leaf mould resistance locus in tomato by *Fulvia fulva*"

**Supplementary Information.** Predicted Avr9B-like protein and *Avr9B*-like gene sequences in different fungi. Predicted N-terminal signal peptide sequences for extracellular targeting are underlined. Predicted intrinsically disordered regions are highlighted in yellow and predicted transmembrane regions are highlighted in grey. Cysteine residues are depicted in bold font. Predicted intron sequences are in red text. Not all predicted protein and gene sequences are full-length. Some proteins and genes have been reconstructed through removal of a large sequence insertion (sequence insertion points in predicted genes are highlighted in green) or removal of an intron that lacks a canonical GT–AG splice site (variant splice sites are highlighted in cyan).

### Group 1 protein sequences

*Fulvia fulva* 0WU
MRSSIVSLLAIQLAFLAVGNTMPTDPAGYTDLSAPEMEGSTDLSAPDVEKSTDLSAPDVEKSTGLSAPEMKEDVN
SSPQSSLCENLHIEVLGHRLGWCAGCKRLEICVTVLAETGASCAAAYVPGAQAALLACLLSIGVDEYEASKCTDG
LC*
*Alternaria gansuensis* LYZ1412
MKLSSIGLPILQLACLAVALKVPEPASPSSDSSYLSAEFGSLCEHFSHTFNNGVSLGWHAGCKRLEKCLVGFTYE
IFLCVESLKVDPRIGLLTCMTEAIGDTFAKYCLAGLY*
*Alternaria* sp section Undifilum W740-01
MKLSSIGLPILQLACLAVAQKVLEPASPSSNSSHIPTNLGSFCEHWNQTYKNGIHIGWHGGCKRLEECLGVLLFE
AVVCIEVLKSDIQAGMLSCMASSVGDAFAKKCWDGIY*
*Curvularia clavata* yc1106
MKFFQILPVLSLASLAMATPAPATVPQTKNFVPEPANPPKQPANIPLY*SSSLPLMDSTSTCERWHHTFKNGHSL
GWCAGCKRLERCGFLLTAEAAAFLAAITVWGPAAILGCLVSVTATDFANYCFQGVC*
*Curvularia geniculata* P1
MKFFQILPLLQLAGFALATPAAAPAPVPAKKLVKEPANPPKQSTNIPLYLIPEDSGSICENWYHTFKNGNKIGWC
AGCKRLERCGVLLTVEAAACIAGIAVMGPAAILGCLLEVAADDFANYCFEGLC*
*Curvularia geniculata* W3
MKFFQILPLLQLAGFALATPAAAPAPVPAKKLVKEPANPPKQSTNIPLYLIPEDSGSICENWYHTFKNGNKIGWC
AGCKRLERCGVLLTVEAAACIAGIAVMGPAAILGCLLEVAADDFANYCFEGLC*
*Curvularia lunata* CX-3
MKFFQILPLLQLAGFALATPAAAPAPVPAKKLVK*PANPPKQPANIPLYLIPEDSGSICENWHHTFKNGNKIGWC
AGCKRLERCGVLLTVEAAACIAGIAVMGPAAILGCLLEVAADDFANYCFEGLC*
*Curvularia lunata* W3
MKFFQILPLLQLAGFALATPAAAPAPVPAKKLVKEPANPPKQSTNIPLYLIPEDSGSICENWYHTFKNGNKIGWC
AGCKRLERCGVLLTVEAAACIAGIAVMGPAAILGCLLEVAADDFANYCFEGLC*
*Curvularia papendorfii* UM 226
MKFFQILPLLQLAGIALATPAAAPAPVPAKKLVKEPANPPKQPANIPLYLIPEDSGSICENWHHTFKNGNKIGWC
AGCKRLERCGVLLTVEAAACIASIAVMGPAAILGCLLEVAADDFANYCFEGLC*
*Curvularia spicifera* FR9030
MKFFQVLPLLQLAGVALATPAQAPAKPFKTVTEPINPPKQPANIPTYIIPEDSGSICENWHHTFKNGNRIGWCAG
CKRLERCGVLLTVEAAVCLAGLTVFGPAAILGCLLEVAADDFANYCFDGLC*
*Curvularia spicifera* FR9030
MKLSSISLPILHLACLVAGQKVPEPANPPPEPANNKHCVIPANVGSVCENWHHTFNKQITIGWCAGCKRLERCGY
LLTVEAAACIAGLILWETKAVLGCLLEAAADDFATYCLDGVC*
*Curvularia* sp ZM96
MKFFQVLPLLQLAGVALATPAQAPAKPFKTVTEPINPPKQPANIPTYIIPEDSGSICENWHHTFKNGNRIGWCAG
CKRLERCGVLLTVEAAACLAGLEIFGPAAILGCLLEVAADDFANYCFDGLC*
*Curvularia* sp ZM96
MKLSSISLPILHLACLVAGQRVPEPANPPPEPANNKHYVIPANVGSVCENWHHTFNKQITIGWCAGCKRLERCGY
LLTVEAAACIAGLILWETKAVLGCLLEAAADDFATYCLDGVC*
*Stemphylium lycopersici* CIDEFI-212
MKLSTISLSILQLACLAAAAVPAQKSAIAVPGPKAAGPPAQKLSVAPPGEKPAAILHPGRKSVTEPAKPPPMPPN
VQPMFVTDSSFCEGWHHTFKNGNKIGWCAGCKRLERCGLLLTIEAAACIGGLAVFGPAAILGCAAEVVSDDFASY
CVDGIC*
*Stemphylium lycopersici* CIDEFI-213

MKLSTISLSILQLACLAAAAVPAQKSAIAVPGPKAAGPPAQKLSVAPPGEKPAAILHPGRKSVTEPAKPPPMPPN
VQPMFVTDSSFCEGWHHTFKNGNKIGWCAGCKRLERCGLLLTIEAAACIGGLAVFGPAAILGCAAEVVSDDFASY
CVDGIC*
*Stemphylium lycopersici* CIDEFI-216
MKLSTISLSILQLACLAAAAVPAQKSAIAVPGPKAAGPPAQKLSVAPPGEKPAAILHPGRKSVTEPAKPPPMPPN
VQPMFVTDSSFCEGWHHTFKNGNKIGWCAGCKRLERCGLLLTIEAAACIGGLAVFGPAAILGCAAEVVSDDFASY
CVDGIC*
*Stemphylium vesicarium* 173-1a-13FI1M3
MKLSTITLPILSFACLATAAVPAPKPAVAAPGQKAVAPPAQKLAVAANPVHPPAVAAIPGEKPAVVLHPGRKSVI
EPARPPPQPANIQPMYVTDSSFCEGWHHTFKGGNKIGWCAGCKRLERCGILLTVEAAACLAGLTVFGPAAILGCA
AEVAGDDFASYCVDGLC*
*Stemphylium vesicarium* On16-63 18160
MKLSTITLPILSFACLATAAVPAPKPAVAAPGQKAVAPPAQKLAVAANPVHPPAVAAIPGEKPAVVLHPGRKSVI
EPARPPPQPANIQPMYVTDSSFCEGWHHTFKGGNKIGWCAGCKRLERCGILLTVEAAACLAGLTVFGPAAILGCA
AEVAGDDFASYCVDGLC*
*Stemphylium vesicarium* On16-391 18415
MKLSTITLPILSFACLATAAVPAPKPAVAAPGQKAVAPPAQKLAVAANPVHPPAVAAIPGEKPAVVLHPGRKSVI
EPARPPPQPANIQPMYVTDSSFCEGWHHTFKGGNKIGWCAGCKRLERCGILLTVEAAACLAGLTVFGPAAILGCA
AEVAGDDFASYCVDGLC*

### Group 2 protein sequences

*Bipolaris bicolor* ML9021
MLPFITVVLAFLVNSTLAAPSTKFLSYSSSLRSRF*ENQQRRVCTGSHIGKFGWCGGCKRLETCGVYLAFLSGCS
SLSIPGVLGCAGAIGFLGANADYCLEGLCVCQGGHCKT*
*Bipolaris cookei* LSLP18.3 reconstructed
MLPLVTIVLAFLANSTLAAPPTKLLPYDSSLESQFQENQERSICTGFHIGEFGWCGGCKRLETCVANLAFLSACS
SATVVGVLGCAGAVGFNGANADYCLEGLCICQGGHCKT*
*Bipolaris oryzae* ATCC 44560
FQDNQGRSICAGFHFGKFGWCGNYKQPETCKADLAFLSSCSSVIIPGVLSCASAVVFFRANADCGLEGPCVCQGG
HCKTSTI*
*Bipolaris oryzae* Bo-Gvt
FQDNQGRSICAGFHFGKIGWCGNYKQPETCKADLAFLSSCSSVIIPGVLSCASAVVFFRANADCGLEGPCVCQGG
HCKTSTI*
*Bipolaris* sp ADL-507 reconstructed
GICTGLHIGKYGWCGGCKRFETCSAYLAFMSGCTSFTPTGVLACVGASGFLAANADYCLEGLCVCQGGKCHVKIP
PV*
*Bipolaris sorokiniana* ND90Pr
LPGVFSCANAVGTLGYIADYCLEGLCVCQGGHCKTWSITQVD*
*Bipolaris sorokiniana* BRIP10943a
LPGVFSCANAVGTLGYIADYCLEGLCVCQGGHCKTWSITQVD*
*Bipolaris sorokiniana* BRIP27492a
LPGVFSCANAVGTLGYIADYCLEGLCVCQGGHCKTWSITQVD*
*Bipolaris sorokiniana* BS112
LPGVFSCANAVGTLGYIADYCLEGLCVCQGGHCKTWSITQVD*
*Bipolaris sorokiniana* WAI2411
LPGVFSCANAVGTLGYIADYCLEGLCVCQGGHCKTWSITQVD*
*Bipolaris sorokiniana* Pusa2
WCAGCKRLAKCLPGVFSCANAVGTLGYIADYCLEGLCVCQGGHCKTWSITQVD*
*Bipolaris sorokiniana* LK93
LPGVFSCANAVGTLGYIADYCLEGLCVCQGGHCKTWSITQVD*
*Bipolaris victoriae* FI3
MLPLITVVLAFLANSTLAAPPTKPLPYSPSLGSQFQENQERSICTGLHVGKFGWCGGCKRLETCSAYLAFLSGCT
ATELPGLLGCAGSISFVKASADYCLEGLCICQGGRCKT*
*Cercospora canescens* BHU
MRFTTFLLPFLGLSKAAVLPVDEVASPNDNICTKAHVGKYGWCAGCQRLETCTALGLYNVLNCLDVATIGLCIAQ
TAFGVANWNFCVAGLCICKGPNAKPCFIN*
*Cercospora janseana* RL44
MHRSAIYAALLLVASNVVRMWAAPIQEEQLQAVSARSICTHWHFRHIGWCTGCTRLEYRTAFFTLLISECLAAE
*Cercospora sojina* 2.2.3

MRSTTVLLSLLGLSYGAVLDVDYQKAAIELEDSFCGGLHVGKGGWCAGCQRLENCIGFGLFNIINCLAVLDNPVA
IRGCAAQTAFSLGSWNYCAVGFCVCQPYGSQKCTISS*
*Cercospora sojina* CCC
MRSTTVLLSLLGLSYGAVLDVDYQKAAIELEDSFCGGLHVGKGGWCAGCQRLENCIGFGLFNIINCLAVLDNPVA
IRGCAAQTAFSLGSWNYCAVGFCVCQPYGSQKCTISS*
*Cercospora sojina* N1
MRSTTVLLSLLGLSYGAVLDVDYQKAAIELEDSFCGGLHVGKGGWCAGCQRLENCIGFGLFNIINCLAVLDNPVA
IRGCAAQTAFSLGSWNYCAVGFCVCQPYGSQKCTISS*
*Cercospora sojina* RACE15
MRSTTVLLSLLGLSYGAVLDVDYQKAAIELEDSFCGGLHVGKGGWCAGCQRLENCIGFGLFNIINCLAVLDNPVA
IRGCAAQTAFSLGSWNYCAVGFCVCQPYGSQKCTISS*
*Cercospora sojina* S9
MRSTTVLLSLLGLSYGAVLDVDYQKAAIELEDSFCGGLHVGKGGWCAGCQRLENCIGFGLFNIINCLAVLDNPVA
IRGCAAQTAFSLGSWNYCAVGFCVCQPYGSQKCTISS*
*Curvularia geniculata* P1
MLPRMMIIVAFLANRTLAAPSTNMIAHDSLVESPFQQNQERSICTGFHIGKFGWCGGCKRFESCSAFLAGLAACP
SLSVAGVLTCASSAGILGSRADYCLEGLCICQGGHCTT*
*Curvularia geniculata* W3
MLPRMIIIMAFLANRTLAAPSTNMIAHDSLVEPQFQHNQERSICTGFHIGKFGWCGGCKRFESCSAFLAGLAACP
SLSVAGVLTCASSAGILGSRADYCLEGLCICQGGHCKT*
*Curvularia hawaiiensis* PK9021
MQPLITIALALQTTTTLAVPVIDAKLSGTDLAIPLTDIAAQSTDLTDLQARDSSLAPRGICTQFKIGKYGWCGGC
KRFETCSAYLAFMSGCASFTPTGVLACVGASGFLAANADYCLEGLCVCQGGKCHVKIPPA*
*Curvularia lunata* Cl-Gvt
SSVII
*Curvularia lunata* CX-3
MLPRMMIIVAFLANRTLAAPSTNMIAHDSLVESPFQQNQERSICTGFHIGKFGWCGGCKRFESCSAFLAGLAACP
SLSVAGVLTCASSAGILGSRADYCLEGLCICQGGHCTT*
*Curvularia lunata* W3
MLPRMIIIMAFLANRTLAAPSTNMIAHDSLVEPQFQHNQERSICTGFHIGKFGWCGGCKRFESCSAFLAGLAACP
SLSVAGVLTCASSAGILGSRADYCLEGLCICQGGHCKT*
*Curvularia papendorfii* UM 226
MLPRMIIIMAFLANRSLAAPSTNMMAHDSLVEPQFQHNQERSICTGFNIGKFGWCGGCKRFESCSAFLAGLAACP
SLSVAGVLTCASSAGILGSRADYCLEGLCICQGGHCKT*
*Curvularia spicifera* FR9030
MQPLITIALALQVATTIAIPVTDVTVKDIDLTLSGTDLTLAGTDLAIPLADLAAPGTDLTDVQARDSSLESRSIC
TGFHVGRYGWCGGCRRLEACGAYLGFLSGCSSLSVAGVVGCAAASGVLAANANYCLEGLCVCQGGKCHV*
*Curvularia* sp ZM96
MQPLITIALALQVATTIAIPVTDVTVKDIDLTLSGTDLTLAGTDLAIPLADLAAPGTDLTDVQARDSSLESRSIC
TGFHIGRYGWCGGCRRLEACGAYLGFLSGCSSLSVAGVVGCAAASGVLAANANYCLEGLCVCQGGKCHV*
*Exserohilum rostratum* B6094
MVAIHALTFAFLTSSILAAPPTDLKTRDFPVELEVAQNEERSVCTGFNIGKFGWCGGCKRLETCSAYLAFLSGCT
SVSVVGVLGCAGAVGFLGANADYCLEGLCVCQGNHCKT*
*Exserohilum rostratum* B6096
MVAIHTITLAFLASSVLAAPPTDLKTRDFPVELEVAQNEERSVCTGFNIGKFGWCGGCKRLETCSAYLAFLSGCT
SVSVVGVLGCAGAVGFLGANADYCLEGLCVCQGNHCKT*
*Exserohilum rostratum* B6171
MVAIHTITLAFLASSVLAAPPTDLKTRDFPVELEVAQNEERSVCTGFNIGKFGWCGGCKRLETCSAYLAFLSGCT
SVSVVGVLGCAGAVGFLGANADYCLEGLCVCQGNHCKT*
*Exserohilum rostratum* B6177
MVAIHTITLAFLASSVLAAPPTDLKTRDFPVELEVAQNEERSVCTGFNIGKFGWCGGCKRLETCSAYLAFLSGCT
SVSVVGVLGCAGAVGFLGANADYCLEGLCVCQGNHCKT*
*Exserohilum rostratum* B6207
MVAIHTITLAFLASSVLAAPPTDLKTRDFPVELEVAQNEERSVCTGFNIGKFGWCGGCKRLETCSAYLAFLSGCT
SVSVVGVLGCAGAVGFLGANADYCLEGLCVCQGNHCKT*
*Exserohilum rostratum* B6272
MVAIHALTFAFLASSVLAAPPTDLKTRDFPVELEVAQNEERSVCTGFNIGKFGWCGGCKRLETCSAYLAFLSGCT
SVSVVGVLGCAGAVGFLGANADYCLEGLCVCQGNHCKT*
*Exserohilum rostratum* B6284
MVAIQAITFAFLASSVLAAPPTDLKTRDFPVELEVAQNEERSVCTGFNIGKFGWCGGCKRLETCSAYLAFLSGCA
NISVVGVLGCAGAAGFLGANADYCLEGLCVCQGNHCKT*

*Exserohilum rostratum* B9826
MVAIHTITFAFLASSVLAAPPTDLKTRDFPVELEVAQNEERSVCTGFNIGKFGWCGGCKRLETCSAYLAFLSGCT
SVSVVGVLGCAGAVGFLGANADYCLEGLCVCQGNHCKT*
*Exserohilum rostratum* BF9006
MVAIHALTFAFLASSVLAAPPTDLKTRDFPVELEVAQNEERSVCTGFNIGKFGWCGGCKRLETCSAYLAFLSGCT
SVSVVGVLGCAGAVGFLGANADYCLEGLCVCQGNHCKT*
*Exserohilum rostratum* ER1
MVAIHAITFAFLASSVLAAPPTDLKTRDFPVELEVAQNEERSVCTGFNIGKFGWCGGCKRLETCSAYLAFLSGCA
NISVVGVLGCAGAAGFLGANADYCLEGLCVCQGNHCKT*
*Exserohilum rostratum* LWI
MVAIHALTFAFLASSILAAPPTDLKTRDFPVELEVAQNEERSVCTGFNIGKFGWCGGCKRLETCSAYLAFLSGCT
SVSVVGVLGCAGAVGFLGANADYCLEGLCVCQGNHCKT*
*Exserohilum rostratum* M 18 0203
MVAIHALTFAFLASSVLAAPPTDLKTRDFPVELEVAQNEERSVCTGFNIGKFGWCGGCKRLETCSAYLAFLSGCT
SVSVVGVLGCAGAVGFLGANADYCLEGLCVCQGNHCKT*
*Exserohilum rostratum* SR-KPL1
MVAIHTITFAFLASSVLAAPPTDLKTRDFPVELEVAQNEERSVCTGFNIGKFGWCGGCKRLETCSAYLAFLSGCT
SVSVVGVLGCAGAVGFLGANADYCLEGLCVCQGNHCKT*
*Exserohilum rostratum* ZM170581
MVAIHAITFAFLASSVLAAPPTDLKTRDFPVELEVAQNEERSVCTGFNIGKFGWCGGCKRLETCSAYLAFLSGCA
NISVVGVLGCAGAAGFLGANADYCLEGLCVCQGNHCKT*
*Pseudocercospora cruenta* Pscow-1
MHPFIMILSLHAVTTVLAMPNPVAEPNALALPLALPFIENIDGSDKHKKHKHKQNEEDIEDQEEQKSTKQKHKQQ
DDDDDDDSKKDSVSNEDFSLPKCPLSIFKKAGWCGGCQRLETCVGTIGFFAASCLAGSLTVAGALSCFGSAGFAA
GEWNYCVDGVCACQGNTKCHK*
*Pseudocercospora eumusae* CBS 114824
MHSSFLWILSLQSVVTAMPNPTAVAADSICPTALFHKAGWCGGCQRLETCIGSLGFFASSCIAGSLTTSSVLSCI
GSTGFAAGTWNYCVDGVCACQGNTQCHI*
*Pseudocercospora fuligena* PF001
MHPLIIIVSLHAASNVLAMPNPVAEPNALALPHALPMIEEIDGSDKHKKHKQKQNEEDIGDEEEQKSTKHKHKQH
DDHDDDDSKKDSVSDEDFSFHLPKCPTSTLNLAGWCGGCRRLETCLGTLGFLTASCLFGSVTIGGVLSCFGSVGF
ATGEWNYCVDGVCACQGKPECHK*
*Pseudocercospora musae* CBS 116634
MHLFQWILCLHSATAVLAMPNPVAEPNALALPLAHSMVEEKSAKHKDKREDDHDQGAEGDPVSNTDISIKCPLSL
FSKAGWCGGCSRLETCVGTLGFFVSSCLVGSVTIAGVAACAGSTGFAAGSWNNCVDGLCLCQGNTKCHI*
*Pseudocercospora pini-densiflorae* CBS 125139
MHPFIWILSLHSVTTVLAMPNPVAEPNALALPLALPLIEEIDESDKHKKQKHKQDKEDIDAEEGQKSTKHKHKQQ
DDDDDDDSKKDSVSNEDFSIHLPKCPTSILNKAGWCAGCQRLETCLGTLGFFAASCLAGSLTVAGALSCFGSTGF
AAGEWNYCVDGVCACKGNTKCHK*

### Group 3 protein sequences

*Pseudocercospora macadamiae* BRIP 55526
MHFPRLLTIMCYSSITMGHVTWKTCPNPAWYIDEDITPSDSDLNVRGTDKHDPQSVDPPTQHSHSHHLKTQKLQI
ITNGEASSHAKAAIKKAAAAGTVEADLSLCTHLHIGKFGWCAGCRRLEKCIGVGFFVIGGLLAWIASCVLTGGAS
ATTIAAFISFLGSIGFSVSTFNDCIIGLCACQPGRKGSHKCTL*
*Pseudocercospora ulei* ERN8
MRILAILYTISSLLILCYATPATSPNFNWYREKRVTPHFSLNSRDADFYNTEVTFNPQPAHRAIQEAKRVGTIEA
DLSMCRHIVLFGRHIWCAGCKRLAKCIGTGCFGLAGMGALIGLIFMTDGAILPIIVAYVNGIGGLSMGVASFADC
LDGLCPCKHGRCVGDVHGSE*
*Pseudocercospora ulei* GCL012
MRILAILYTISSLLILCYATPATSPNFNWYREKRVTPHFSLNSRDADFYNTEVTFNPQPAHRAIQEAKRVGTIEA
DLSMCRHIVLFGRHIWCAGCKRLAKCIGTGCFGLAGMGALIGLIFMTDGAILPIIVAYVNGIGGLSMGVASFADC
LDGLCPCKHGRCVGDVHGSE*
*Pseudocercospora ulei* ERN8
MRVLAILYTISSLMALCYATPATSPNFDWYRDKQMTPNFRLNSRDVALHQIKLGLDVEQAYNTIHTAYRETYESK
AVAPYEAELTRSMCTHVSIAGVHIWCAGCKRLAKCIAAGCFALAGFGGLLALFFVTGSVILPAILAFFNAVGGFS
ASVAVFNDCMEGLCPCKGGKCKL*
*Pseudocercospora ulei* GCL012

MRVLAILYTISSLMALCYATPATSPNFDWYRDKQMTPNFRLNSRDVALHQIKLGLDVEQAYNTIHTAYRETYESK
AVAPYEAELTRSMCTHVSIAGVHIWCAGCKRLAKCIAAGCFALAGFGGLLALFFVTGSVILPAILAFFNAVGGFS
ASVAVFNDCMEGLCPCKGGKCKL*
*Pseudocercospora ulei* ERN8
MRLLAILYTMSSLLTLCYGTPATSPNFDWYREKRVTPNFGLNSRDADFHDADLTFNPQPAYRAIERYKSIGTDDA
DWNMCKHVSLFGLHVWCAGCKRLEKCLAAGLFAIAGMAALIGLIFFTDGAILPIINAFLNAVGGVAFSIAVFDDC
LEGLCPCQHGECTNGDKRHHH*
*Pseudocercospora ulei* GCL012
MRLLAILYTMSSLLTLCYGTPATSPNFDWYREKRVTPNFGLNSRDADFHDADLTFNPQPAYRAIERYKSIGTDDA
DWNMCKHVSLFGLHVWCAGCKRLEKCLAAGLFAIAGMAALIGLIFFTDGAILPIINAFLNAVGGVAFSIAVFDDC
LEGLCPCQHGECTNGDKRHHH*
*Pseudocercospora ulei* ERN8
MRALAVLYTISSLLAWCYAIPTTSPNFDWYREKRVSPNFSLNSRDADYHDIETTFNPEPARRAIEAAKRVGTFEA
DMSLCSHVTLFHRHIWCAGCKRLAKCIGAGLFQIAGFGALIALIFITDGAILPIIVAWLAAVGSVASGIATIDEC
LDGLCPCKHGQCDL*
*Pseudocercospora ulei* GCL012
MRALAVLYTMSSLLALCYAIPTTSPNFDWYREKRVSPNFSLNSRDADYHDIETTFNPEPARRAIEAAKRVGTFEA
DMSLCSRVTFFHRHIWCAGCKRLAKCIGAGVFEIAGFGALIALIFITDGTILPIIVAWLSAVGTVASGVATLDDC
LDGLCPCKHGNCDV*
*Pseudocercospora ulei* ERN8
Frame 1
MRVLAILYTISSLLTSCLA*L*LVSEKKVDA*FPPK*PRCPTSVRWKRPSIPNRPYRPSKRLKESALSKPT*VCA
ATLPSLAYMX
Frame 2
CASSPSCTLSVAS*PRASPNFNWYRKKRLTPNFRLNSRDVRPP*DGRDLQSPTGPTGHRSG*KSRHFRSRHEYVQ
PHYPLWHTCX
Frame 3
ARPRHPVHYQ*PLNLVPRLTLTGIGKKG*RLISA*IAAMSDLREMEETFNPQPALQAIEAAKRVGTFEADMSMCS
HITLFGIHV
*Pseudocercospora ulei* GCL012
VGTFEADMSMCSHITLFGIHV
*Septoria petroselini* CBS 182.44
MRFLYTFAAICYGYIAIVLASPTRSPNFLWYEAKKMSPNFELLARSISTNTPPPVDGKWYSEILAVIAQAKLDPE
YDFEREDENDSKFKLPKVHCTGLQIQISKTKKFGWCAGCQRLEKCLGVLFFVKGGIAAYIYAFYFSGGLSVPATV
AFLNVVGSIGFAASCFNTCVDGFCATKSGNYHICGD*

### Group 1 gene sequences

*Fulvia fulva* 0WU
ATGCGATCATCTATTGTCTCCCTCCTCGCCATCCAGCTGGCTTTCCTTGCCGTTGGTAACACGATGCCGACCGAC
CCTGCAGGATACACGGATCTATCAGCTCCAGAAATGGAAGGATCCACGGATCTATCAGCTCCAGATGTGGAAAAA
TCCACGGATCTATCAGCTCCAGATGTGGAAAAATCCACGGGTCTATCGGCTCCAGAAATGAAAGAAGACGTCAAC
TCCTCTCCACAATCCTCTCTTTGTGAAAACCTTCATATCGAGGTACTGGGCCACCGGCTCGGTTGGTGTGCGGGT
TGTAAACGACTTGAGATTTGCGTCACCGTTCTCGCAGAGACTGGCGCTTCATGCGCTGCAGCCTATGTACCAGGG
GCTCAAGGTATGTTTGAATAACGAAGGCCTGGCTTGGATGTGTATGATAACAGCAATAGCCGCACTCCTTGCCTG
TTTGCTCTCAATCGGTGTCGATGAATATGAGGCGTCAAAGTGTAGGTGTCCTACTCACTGTACATCACATGTCAA
GCAAAGGCTGACTGGGATTAGGTACCGATGGGTTGTGTTAG
*Alternaria gansuensis* LYZ1412
ATGAAGCTATCCAGCATCGGTCTCCCTATCCTCCAGCTGGCTTGCCTTGCTGTCGCGCTGAAGGTTCCTGAACCA
GCTAGTCCATCGAGCGATTCATCATACCTCTCTGCAGAATTTGGCTCTTTATGTGAGCACTTTAGTCACACATTT
AATAATGGTGTAAGTCTTGGATGGCATGCTGGCTGCAAGCGACTTGAGAAATGCTTGGTGGGTTTTACCTATGAG
ATTTTTCTCTGTGTCGAAAGTTTGAAAGTTGATCCTCGAAGTACGTCTATAATCTCTTCCATATTAACTATAGTC
TCTAATATAATATAGTTGGTTTACTCACGTGCATGACTGAAGCAATTGGTGACACATTTGCAAAATATTGTAAGT
TTTTAAACCTCTAAATACTATAAGCGGGCTTGATATTAACTAGTTTAGGTTTGGCGGGTCTATACTAG
*Alternaria* sp section Undifilum W740-01
ATGAAGCTATCCAGCATCGGTCTTCCTATCCTCCAGCTAGCTTGCCTTGCTGTCGCGCAGAAGGTTCTTGAACCA
GCTAGTCCATCGAGCAATTCATCACACATTCCTACAAATCTTGGCTCTTTTTGTGAGCACTGGAATCAGACATAT
AAGAATGGTATACATATTGGATGGCATGGTGGCTGCAAGCGACTTGAGGAATGCTTGGGGGTTTTACTCTTTGAG
GCTGTTGTCTGTATCGAAGTTTTGAAGTCTGATATTCAAGGTACGTCTATAATCTCTTCTTTATTAACTATAGAC
TCTAATAAAATATAGCTGGTATGCTCTCGTGCATGGCTTCATCAGTTGGTGACGCATTTGCAAAGAAATGTAAGT
TTTTAAACCTCTAAATACTATAAGCGGGCTTGATATAAACTAATTTAGGTTGGGACGGTATATACTAG

*Curvularia clavata* yc1106
ATGAAGTTCTTCCAGATCCTCCCTGTTCTTTCCCTTGCCAGCCTTGCTATGGCTACTCCAGCACCAGCAACAGTA
CCGCAAACGAAAAATTTTGTTCCTGAACCTGCCAACCCACCCAAACAGCCGGCAAACATTCCACTATATTAATCT
TCATCTCTGCCTCTTATGGATTCAACCTCCACGTGTGAGAGATGGCACCACACGTTCAAGAACGGACACAGTCTC
GGATGGTGTGCGGGCTGCAAGCGTCTTGAGCGATGCGGTTTCCTTTTGACGGCTGAGGCTGCTGCCTTTCTTGCT
GCTATCACAGTCTGGGGTCCAGGTGAGTTTAAGAATTGCTGTTCTATTCACTATATGTTCTGATCGATTATAGCT
GCTATCCTTGGTTGCCTTGTTTCAGTCACTGCTACTGACTTTGCCAACTACTGTGAGTCTTCAGCCCTCCCAACC
TCTAGAGCAGGCAACTTTCTAACAAATTTTAGGCTTCCAAGGAGTCTGTTAG
*Curvularia spicifera* FR9030
ATGAAGTTCTTCCAGGTCCTCCCACTCCTTCAACTCGCTGGCGTTGCTCTGGCTACTCCAGCACAAGCACCAGCA
AAGCCGTTCAAGACTGTTACTGAACCGATCAACCCACCTAAACAGCCTGCAAACATCCCAACTTATATAATTCCT
GAGGACTCAGGTTCCATTTGCGAGAACTGGCATCACACGTTCAAAAACGGAAACAGGATTGGATGGTGTGCGGGC
TGCAAGCGACTTGAGCGATGCGGTGTCCTCTTGACGGTTGAGGCTGCTGTTTGTCTCGCGGGTCTCACAGTCTTT
GGTCCAGGTATGTTCAAAATTCGCTATTCTATTCACTATAGGTTCTGATAGATCATAGCTGCTATCCTTGGTTGC
CTTCTTGAAGTCGCTGCTGATGACTTTGCGAACTACTGTAAGTTTTGAGCCCTTTCAGCGTTTTCAGAGCAGGCA
AGGTATTAATAAATATTAGGCTTCGATGGACTCTGCTAG
*Curvularia* sp ZM96
ATGAAGTTCTTCCAGGTCCTCCCACTCCTTCAACTCGCTGGCGTTGCTCTGGCTACTCCAGCACAAGCACCAGCA
AAGCCGTTCAAGACTGTTACTGAACCGATCAACCCACCTAAACAGCCTGCAAACATCCCAACTTATATAATTCCT
GAGGACTCAGGTTCCATTTGCGAGAACTGGCATCACACGTTCAAAAACGGAAACAGGATTGGATGGTGTGCGGGC
TGCAAGCGACTTGAGCGATGCGGTGTCCTCTTGACGGTTGAGGCTGCTGCTTGTCTCGCGGGTCTTGAAATCTTT
GGTCCAGGTATGTTCAAAATTTGCTATTCTATTCACTATAGGTTCTGATATATCATAGCTGCTATCCTTGGTTGC
CTACTTGAAGTCGCTGCTGATGACTTTGCGAACTACTGTAAGTTTTGAGCCCTTTCAGCGTTTTCAGAGCAGGCA
AGGTATTAATAAATATTAGGCTTCGATGGACTCTGCTAG
*Curvularia* sp ZM96
ATGAAGCTATCTAGCATCAGTCTCCCTATCCTTCACCTAGCTTGCCTTGTTGCTGGACAAAGGGTCCCAGAACCA
GCTAATCCACCACCTGAACCGGCAAACAATAAACACTATGTTATTCCTGCGAATGTTGGATCTGTCTGTGAAAAC
TGGCATCACACATTCAACAAACAGATCACAATTGGATGGTGCGCAGGCTGCAAGCGACTTGAAAGATGTGGTTAT
CTTTTAACAGTTGAGGCTGCTGCCTGCATTGCAGGATTGATACTTTGGGAAACAAGTAGGTTTAGAATCCATTTC
TCCATACACAGGCACTAATGAAATTTAGAGGCTGTTCTTGGGTGTCTGCTGGAAGCAGCTGCAGACGATTTCGCG
ACCTACTGTAAGTTTCCAATCTTTAGTCTTTGAGCGCAAACAAGCGGTTAATTGTTTTTAGGTCTTGATGGTGTC
TGCTAG
*Curvularia geniculata* P1
ATGAAATTCTTTCAGATCCTCCCACTTCTTCAACTCGCTGGGTTTGCTCTGGCTACTCCAGCAGCAGCACCAGCG
CCAGTACCAGCGAAAAAGCTGGTTAAGGAACCGGCCAATCCACCAAAACAATCGACAAACATTCCCTTGTACCTC
ATTCCTGAAGACTCAGGCTCCATCTGCGAGAACTGGTACCACACATTCAAAAATGGAAACAAGATTGGATGGTGT
GCGGGCTGCAAGCGCCTTGAGCGATGCGGTGTTCTTTTGACGGTTGAGGCTGCCGCGTGTATTGCAGGTATTGCA
GTCATGGGACCAGGTATATTCAAGGTTCACTGTTCTGTATACTCCAGATACTGATGCATTATAGCTGCTATCCTC
GGTTGCCTTCTCGAAGTCGCTGCTGATGACTTTGCAAACTACTGTAAGTTTTGACCCCTTTCAACAATCTACATT
AGGCAAGGTGCTAATGGGTACTAGGCTTCGAGGGTCTCTGCTAG
*Curvularia geniculata* W3
ATGAAATTCTTTCAGATCCTCCCACTTCTTCAACTCGCTGGGTTTGCTCTGGCTACTCCAGCAGCAGCACCAGCG
CCAGTACCAGCGAAAAAGCTGGTTAAGGAACCGGCCAATCCACCAAAACAATCGACAAACATTCCCTTGTACCTC
ATTCCTGAAGACTCAGGCTCCATCTGCGAGAACTGGTACCACACATTCAAAAATGGAAACAAGATTGGATGGTGT
GCGGGCTGCAAGCGCCTTGAGCGATGCGGTGTTCTTTTGACGGTTGAGGCTGCCGCGTGTATTGCAGGTATTGCA
GTCATGGGACCAGGTATATTCAAGGTTCACTGTTCTGTATACTCCAGATACTGATGCATTATAGCTGCTATCCTC
GGTTGCCTTCTCGAAGTCGCTGCTGATGACTTTGCAAACTACTGTAAGTTTTGACCCCTTTCAACAATCTACATT
AGGCAAGGTGCTAATGGGTACTAGGCTTCGAGGGTCTCTGCTAG
*Curvularia lunata* CX-3
ATGAAATTCTTTCAGATCCTCCCACTTCTTCAACTCGCTGGGTTTGCTCTGGCTACTCCAGCAGCAGCACCAGCG
CCAGTACCAGCGAAAAAGCTGGTTAAGTAACCGGCCAATCCACCAAAACAACCGGCAAACATTCCCTTGTACCTC
ATTCCTGAAGACTCAGGCTCCATCTGCGAGAACTGGCACCACACATTCAAAAATGGAAACAAGATTGGATGGTGT
GCGGGCTGCAAGCGCCTTGAGCGATGCGGTGTTCTTTTGACGGTTGAGGCTGCCGCGTGTATTGCAGGTATTGCA
GTCATGGGACCAGGTATATTCAAGGTTCACTGTTCTGTATACTCCAGATACTGATGCATTATAGCTGCTATCCTC
GGTTGCCTTCTCGAAGTCGCTGCTGATGACTTTGCAAACTACTGTAAGTTTTGACCCCTTTCAACAATCTACATT
AGGCAAGGTGCTAATGGGTACTAGGCTTCGAGGGTCTCTGCTAG
*Curvularia lunata* W3
ATGAAATTCTTTCAGATCCTCCCACTTCTTCAACTCGCTGGGTTTGCTCTGGCTACTCCAGCAGCAGCACCAGCG
CCAGTACCAGCGAAAAAGCTGGTTAAGGAACCGGCCAATCCACCAAAACAATCGACAAACATTCCCTTGTACCTC
ATTCCTGAAGACTCAGGCTCCATCTGCGAGAACTGGTACCACACATTCAAAAATGGAAACAAGATTGGATGGTGT
GCGGGCTGCAAGCGCCTTGAGCGATGCGGTGTTCTTTTGACGGTTGAGGCTGCCGCGTGTATTGCAGGTATTGCA

GTCATGGGACCAGGTATATTCAAGGTTCACTGTTCTGTATACTCCAGATACTGATGCATTATAGCTGCTATCCTC
GGTTGCCTTCTCGAAGTCGCTGCTGATGACTTTGCAAACTACTGTAAGTTTTGACCCCTTTCAACAATCTACATT
AGGCAAGGTGCTAATGGGTACTAGGCTTCGAGGGTCTCTGCTAG
*Curvularia papendorfii* UM 226
ATGAAATTCTTTCAGATCCTCCCACTTCTTCAACTCGCTGGAATTGCTCTGGCTACTCCAGCAGCAGCACCAGCG
CCAGTACCAGCGAAAAAGCTGGTTAAGGAACCGGCCAATCCACCAAAACAACCGGCAAACATTCCCTTGTACCTC
ATTCCTGAAGACTCAGGCTCCATCTGCGAGAACTGGCACCACACATTCAAAAATGGAAACAAGATTGGATGGTGT
GCGGGCTGCAAGCGCCTTGAGCGATGCGGTGTTCTTTTGACGGTTGAGGCTGCCGCGTGTATTGCAAGTATTGCA
GTCATGGGACCAGGTATGTTCAAGGTTCACTGTTCTGTATACTCCAGATACTGATGCATTATAGCTGCTATCCTC
GGTTGCCTTCTCGAAGTCGCTGCTGATGACTTTGCAAACTACTGTAAGTTTTGACCCCTTTCAACAATCTACATT
AGGCAAGGTGCTAATGGGTACTAGGCTTCGAGGGTCTCTGCTAG
*Stemphylium lycopersici* CIDEFI-212
ATGAAGCTCTCCACCATCTCCCTCTCCATCCTCCAGCTGGCTTGCCTTGCCGCCGCTGCCGTCCCAGCGCAAAAG
TCCGCCATCGCTGTTCCAGGGCCAAAAGCCGCGGGCCCTCCAGCCCAAAAACTTTCCGTTGCTCCTCCAGGCGAA
AAGCCCGCCGCCATCCTCCATCCCGGCCGAAAGTCCGTCACCGAACCCGCCAAACCACCCCCAATGCCCCCAAAC
GTCCAGCCCATGTTCGTCACCGACAGCTCCTTCTGCGAGGGCTGGCACCACACCTTCAAAAACGGGAACAAGATT
GGATGGTGCGCAGGTTGCAAGCGCCTTGAACGATGCGGGCTGCTCTTGACCATCGAGGCTGCCGCTTGTATTGGT
GGGTTGGCGGTTTTTGGTCCTGCGGCTATTCTGGGTTGCGCGGCTGAAGTCGTGAGTGATGATTTCGCGAGTTAT
TGTAAGTTTTCTTTTCGAGATGTACTTTTGGATGGGTTCTAACTGGTTTTAGGTGTCGATGGAATTTGCTAG
*Stemphylium lycopersici* CIDEFI-213
ATGAAGCTCTCCACCATCTCCCTCTCCATCCTCCAGCTGGCTTGCCTTGCCGCCGCTGCCGTCCCAGCGCAAAAG
TCCGCCATCGCTGTTCCAGGGCCAAAAGCCGCGGGCCCTCCAGCCCAAAAACTTTCCGTTGCTCCTCCAGGCGAA
AAGCCCGCCGCCATCCTCCATCCCGGCCGAAAGTCCGTCACCGAACCCGCCAAACCACCCCCAATGCCCCCAAAC
GTCCAGCCCATGTTCGTCACCGACAGCTCCTTCTGCGAGGGCTGGCACCACACCTTCAAAAACGGGAACAAGATT
GGATGGTGCGCAGGTTGCAAGCGCCTTGAACGATGCGGGCTGCTCTTGACCATCGAGGCTGCCGCTTGTATTGGT
GGGTTGGCGGTTTTTGGTCCTGCGGCTATTCTGGGTTGCGCGGCTGAAGTCGTGAGTGATGATTTCGCGAGTTAT
TGTAAGTTTTCTTTTCGAGATGTACTTTTGGATGGGTTCTAACTGGTTTTAGGTGTCGATGGAATTTGCTAG
*Stemphylium lycopersici* CIDEFI-216
ATGAAGCTCTCCACCATCTCCCTCTCCATCCTCCAGCTGGCTTGCCTTGCCGCCGCTGCCGTCCCAGCGCAAAAG
TCCGCCATCGCTGTTCCAGGGCCAAAAGCCGCGGGCCCTCCAGCCCAAAAACTTTCCGTTGCTCCTCCAGGCGAA
AAGCCCGCCGCCATCCTCCATCCCGGCCGAAAGTCCGTCACCGAACCCGCCAAACCACCCCCAATGCCCCCAAAC
GTCCAGCCCATGTTCGTCACCGACAGCTCCTTCTGCGAGGGCTGGCACCACACCTTCAAAAACGGGAACAAGATT
GGATGGTGCGCAGGTTGCAAGCGCCTTGAACGATGCGGGCTGCTCTTGACCATCGAGGCTGCCGCTTGTATTGGT
GGGTTGGCGGTTTTTGGTCCTGCGGCTATTCTGGGTTGCGCGGCTGAAGTCGTGAGTGATGATTTCGCGAGTTAT
TGTAAGTTTTCTTTTCGAGATGTACTTTTGGATGGGTTCTAACTGGTTTTAGGTGTCGATGGAATTTGCTAG
*Stemphylium vesicarium* 173-1a-13FI1M3
ATGAAGCTCTCCACCATCACTCTCCCTATCCTCTCGTTTGCTTGCCTTGCCACCGCTGCTGTCCCAGCGCCCAAG
CCCGCGGTCGCAGCTCCAGGCCAAAAGGCCGTCGCACCCCCAGCTCAAAAGCTCGCCGTAGCTGCCAACCCAGTC
CACCCGCCCGCTGTGGCTGCCATCCCAGGCGAAAAGCCCGCCGTCGTCCTCCATCCCGGCCGAAAGTCCGTCATC
GAGCCCGCTCGCCCGCCCCCACAGCCCGCAAACATCCAGCCCATGTATGTCACCGACAGTTCCTTCTGCGAGGGC
TGGCACCACACCTTCAAAGGCGGGAATAAGATCGGATGGTGCGCTGGCTGCAAGCGCCTTGAACGATGCGGTATT
CTCTTGACCGTCGAGGCTGCTGCTTGTCTTGCCGGTTTAACGGTTTTTGGGCCCGCGGCTATCCTGGGTTGCGCG
GCTGAAGTCGCGGGTGATGATTTTGCGAGTTATTGTAAGTTTCTCTCTCTCTCTCTCTCTCTCTCTCTCTCTCTC
AATCCTTTGGATGGTTTGGTGCGTGCATGATGCTAATATTTTGTAGGCGTCGATGGTCTCTGCTAG
*Stemphylium vesicarium* On16-63 18160
ATGAAGCTCTCCACCATCACTCTCCCTATCCTCTCGTTTGCTTGCCTTGCCACCGCTGCTGTCCCAGCGCCCAAG
CCCGCGGTCGCAGCTCCAGGCCAAAAGGCCGTCGCACCCCCAGCTCAAAAGCTCGCCGTAGCTGCCAACCCAGTC
CACCCGCCCGCTGTGGCTGCCATCCCAGGCGAAAAGCCCGCCGTCGTCCTCCATCCCGGCCGAAAGTCCGTCATC
GAGCCCGCTCGCCCGCCCCCACAGCCCGCAAACATCCAGCCCATGTATGTCACCGACAGTTCCTTCTGCGAGGGC
TGGCACCACACCTTCAAAGGCGGGAATAAGATCGGATGGTGCGCTGGCTGCAAGCGCCTTGAACGATGCGGTATT
CTCTTGACCGTCGAGGCTGCTGCTTGTCTTGCCGGTTTAACGGTTTTTGGGCCCGCGGCTATCCTGGGTTGCGCG
GCTGAAGTCGCGGGTGATGATTTTGCGAGTTATTGTAAGTTTCTCTCTCTCTCTCTCTCTCTCTCTCTCTCAATC
CTTTGGATGGTTTGGTGCGTGCATGATGCTAATATTTTGTAGGCGTCGATGGTCTCTGCTAG
*Stemphylium vesicarium* On16-391 18415
ATGAAGCTCTCCACCATCACTCTCCCTATCCTCTCGTTTGCTTGCCTTGCCACCGCTGCTGTCCCAGCGCCCAAG
CCCGCGGTCGCAGCTCCAGGCCAAAAGGCCGTCGCACCCCCAGCTCAAAAGCTCGCCGTAGCTGCCAACCCAGTC
CACCCGCCCGCTGTGGCTGCCATCCCAGGCGAAAAGCCCGCCGTCGTCCTCCATCCCGGCCGAAAGTCCGTCATC
GAGCCCGCTCGCCCGCCCCCACAGCCCGCAAACATCCAGCCCATGTATGTCACCGACAGTTCCTTCTGCGAGGGC
TGGCACCACACCTTCAAAGGCGGGAATAAGATCGGATGGTGCGCTGGCTGCAAGCGCCTTGAACGATGCGGTATT
CTCTTGACCGTCGAGGCTGCTGCTTGTCTTGCCGGTTTAACGGTTTTTGGGCCCGCGGCTATCCTGGGTTGCGCG

GCTGAAGTCGCGGGTGATGATTTTGCGAGTTATTGTAAGTTTCTCTCTCTCTCTCTCTCTCTCTCTCTCAATCCT
TTGGATGGTTTGGTGCGTGCATGATGCTAATATTTTGTAGGCGTCGATGGTCTCTGCTAG

### Group 2 gene sequences

*Bipolaris bicolor* ML9021
ATGCTTCCATTTATCACTGTTGTCTTGGCCTTCCTGGTTAATAGCACCCTCGCTGCTCCATCTACGAAATTCCTT
TCTTACAGTTCGTCACTGAGATCACGATTCTAAGAGAATCAGCAGAGAAGGGTCTGCACTGGCAGTCATATTGGA
AAATTTGGCTGGTGTGGAGGCTGCAAGCGTTTGGAAACTTGCGGCGTGTATCTTGCTTTCCTTTCTGGTTGCAGC
TCTCTTTCGATTCGTACCTTCCAGTACCTTCCCTCCTCCCCAGGAAGTAGGATAAACATTGTGTGGACGTTCTAT
GCAAACCAAATATCTTGTTCGCTGTAGACTAGGTCATTGCTAACTCCATATCGCAGCTGGCGTGCTTGGCTGCGC
AGGTGCTATTGGATTCCTTGGGGCTAATGCAGATTATTGTAAGTTATTACTATGTTGTTTACATATATATATATA
CTTTAACACTTAACTTACTATTGTCAACAGGTCTCGAAGGACTATGCGTCTGCCAGGGAGGCCATTGTAAGACGT
AG
*Bipolaris cookei* LSLP18.3 reconstructed
ATGCTGCCACTTGTTACTATTGTCTTGGCCTTCCTGGCTAATAGCACCCTCGCTGCTCCGCCTACGAAATTACTT
CCTTATGATTCTTCACTAGAATCACAGTTCCAAGAGAATCAGGAGAGAAGTATCTGCACTGGCTTTCATATTGGA
GAATTTGGCTGGTGTGGAGGTTGCAAGCGTCTTGAAACGTGTGTTGCAAATCTTGCTTTCCTATCTGCATGCAGC
TCTGCTACGGTCGGTATGTTCAAGTTCCTCGCCCCTTCCCTCCTAAAGTGGGGCAACCATTGTGTGGAGGTTTCA
TGCAGAGCAAATATCTTGTTCGCTGTAGACTAGATCATTGCTAACTCTGTATCGCAGTTGGTGTGCTTGGCTGCG
CAGGTGCTGTTGGATTCAATGGGGCTAATGCAGATTATTGTAAGTGATTACTACTTCATATATATTGTATGCTCG
AACACCTGACTTATTATTGTCAAAACAGGTCTCGAAGGACTCTGCATCTGCCAGGGAGGCCACTGTAAGACTTAG
*Bipolaris oryzae* ATCC 44560
TTCCAAGACAATCAGGGGAGAAGTATCTGCGCTGGCTTTCATTTTGGAAAATTTGGTTGGTGTGGAAATTACAAG
CAACCGGAAACTTGCAAGGCAGATCTTGCTTTCCTATCCAGTTGCAGTTCTGTTATAATCCATATGTTCTAGTGG
CTTCCCCACTCCCAAGTTAAAGGCGAATATTGTGTATATCTTTCATAGAGACTCAATATCTCGTTCGTTGTAAAC
TAGATAATTGCTAGATCTATATCACAACTGGTGTGCTTAGCTGCGCAAGTGCTGTTGTATTCTTTAGGGCTAATG
CAGATTGTGGTAAGTGGTTATTGTTTCATATATATATGTATGTATGCTCGAACAGTTGACTTATTGTTGCCACCG
CAGGTCTCGAAGGGCCCTGCGTCTGCCAGGGAGGCCACTGCAAGACCTCAACCATATA
*Bipolaris oryzae* Bo-Gvt
TTCCAAGACAATCAGGGGAGAAGTATCTGCGCTGGCTTTCATTTTGGAAAAATTGGTTGGTGTGGAAATTACAAG
CAACCGGAAACTTGCAAGGCAGATCTTGCTTTCCTATCCAGTTGCAGTTCTGTTATAATCCATATGTTTAGTGG
CTTCCCCACTCCCAAGTTAAAGGCAAATATTGTGTATATCTTTCATAGAGACTCAATATCTCGTTCGTTGTAAAC
TAGATAATTGCTAGATCTATATCACAACTGGTGTGCTTAGCTGCGCAAGTGCTGTTGTATTCTTTAGGGCTAATG
CAGATTGTGGTAAGTGGTTATTGTTTCATATATATATGTATGTATGCTCGAACAGTTGACTTTTGTTGCCACCG
CAGGTCTCGAAGGGCCCTGCGTCTGCCAGGGAGGCCACTGCAAGACCTCAACCATATA
*Bipolaris sorokiniana* ND90Pr
CTCCGTGTGTTCTAGTGCCTCCTTCCCCTTCCCATAAAGTGAGGGCAAACATTGCGTACAGGTTTCATGCAGACC
AAATATCTTGTTCGTTGTAGACTGGATTATTGCTAGCTTCATATCGCAGCTGGTGTGTTTAGCTGCGCGAATGCT
GTTGGGACCCTTGGGTATATTGCAGATTATTGTGAGTGACTACCATTTCAAATATATGTGTGGTCGAACAATTGA
CTTATTGTTGCCAGTACGGGTCTCGAAGGACTCTGCGTCTGCCAAGGAGGCCACTGCAAGACTTGGAGCATAACA
CAAGTAGATTGA
*Bipolaris sorokiniana* BRIP10943a
CTCCGTGTGTTCTAGTGCCTCCTTCCCCTTCCCATAAAGTGAGGGCAAACATTGCGTACAGGTTTCATGCAGACC
AAATATCTTGTTCGTTGTAGACTGGATTATTGCTAGCTTCATATCGCAGCTGGTGTGTTTAGCTGCGCGAATGCT
GTTGGGACCCTTGGGTATATTGCAGATTATTGTGAGTGACTACCATTTCAAATATATGTGTGGTCGAACAATTGA
CTTATTGTTGCCAGTACGGGTCTCGAAGGACTCTGCGTCTGCCAAGGAGGCCACTGCAAGACTTGGAGCATAACA
CAAGTAGATTGA
*Bipolaris sorokiniana* BRIP27492a
CTCCGTGTGTTCTAGTGCCTCCTTCCCCTTCCCATAAAGTGAGGGCAAACATTGCGTACAGGTTTCATGCAGACC
AAATATCTTGTTCGTTGTAGACTGGATTATTGCTAGCTTCATATCGCAGCTGGTGTGTTTAGCTGCGCGAATGCT
GTTGGGACCCTTGGGTATATTGCAGATTATTGTGAGTGACTACCATTTCAAATATATGTGTGGTCGAACAATTGA
CTTATTGTTGCCAGTACGGGTCTCGAAGGACTCTGCGTCTGCCAAGGAGGCCACTGCAAGACTTGGAGCATAACA
CAAGTAGATTGA
*Bipolaris sorokiniana* BS112
CTCCGTGTGTTCTAGTGCCTCCTTCCCCTTCCCATAAAGTGAGGGCAAACATTGCGTACAGGTTTCATGCAGACC
AAATATCTTGTTCGTTGTAGACTGGATTATTGCTAGCTTCATATCGCAGCTGGTGTGTTTAGCTGCGCGAATGCT
GTTGGGACCCTTGGGTATATTGCAGATTATTGTGAGTGACTACCATTTCAAATATATGTGTGGTCGAACAATTGA
CTTATTGTTGCCAGTACGGGTCTCGAAGGACTCTGCGTCTGCCAAGGAGGCCACTGCAAGACTTGGAGCATAACA
CAAGTAGATTGA

*Bipolaris sorokiniana* WAI2411
CTCCGTGTGTTCTAGTGCCTCCTTCCCTTCCCATAAAGTGAGGGCAAACATTGCGTACAGGTTTCATGCAGACCA
AATATCTTGTTCGTTGTAGACTGGATTATTGCTAGCTTCATATCGCAGCTGGTGTGTTTAGCTGCGCGAATGCTG
TTGGGACCCTTGGGTATATTGCAGATTATTGTGAGTGACTACCATTTCAAATATATGTGTGGTCGAACAATTGAC
TTATTGTTGCCAGTACGGGTCTCGAAGGACTCTGCGTCTGCCAAGGAGGCCACTGCAAGACTTGGAGCATAACAC
AAGTAGATTGA
*Bipolaris sorokiniana* Pusa2
CTCCGTGTGTTCTAGTGCCTCCTTCCCCTTCCCATAAAGTGAGGGCAAACATTGCGTACAGGTTTCATGCAGACC
AAATATCTTGTTCGTTGTAGACTGGATTATTGCTAGCTTCATATCGCAGCTGGTGTGTTTAGCTGCGCGAATGCT
GTTGGGACCCTTGGGTATATTGCAGATTATTGTGAGTGACTACCATTTCAAATATATGTGTGGTCGAACAATTGA
CTTATTGTTGCCAGTACGGGTCTCGAAGGACTCTGCGTCTGCCAAGGAGGCCACTGCAAGACTTGGAGCATAACA
CAAGTAGATTGA
*Bipolaris sorokiniana* LK93
CTCCGTGTGTTCTAGTGCCTCCTTCCCCTTCCCATAAAGTGAGGGCAAACATTGCGTACAGGTTTCATGCAGACC
AAATATCTTGTTCGTTGTAGACTGGATTATTGCTAGCTTCATATCGCAGCTGGTGTGTTTAGCTGCGCGAATGCT
GTTGGGACCCTTGGGTATATTGCAGATTATTGTGAGTGACTACCATTTCAAATATATGTGTGGTCGAACAATTGA
CTTATTGTTGCCAGTACGGGTCTCGAAGGACTCTGCGTCTGCCAAGGAGGCCACTGCAAGACTTGGAGCATAACA
CAAGTAGATTGA
*Bipolaris* sp ADL-507 reconstructed
GCGCGTATCTTGCCTTCATGTCTGGATGCACCTCTTTTACCCCTAGTATGTTACGTCCTATGCAACTGTCTCTTA
TACACATCTGACGCTGCCGACGAAGAGGATAGTGTAGATCTCGCTATGCCGCTCACTGATATCGCCGCCCGGGGC
ACTGATCTCACTGATCTGCAAGCTCATGGTCCTTCACTAGAATCACGAGGTATCTGCACGGGATTACATATTGGC
AAATATGGCTGGTGTGGAGGTTGCAAGCGCTTTGAAACTTGCAGCGCGTATCTTGCCTTCATGTCTGGATGCACC
TCTTTTACCCCTAGTATGTTACGTCCTATGCAAAGAAAGCAGCGCCATGATTGAGATCGAGAGCCAGGCAGAATA
ATTGCTGACCTTCTTACACAGCTGGTGTGCTTGCTTGCGTAGGGGCTTCTGGATTTCTTGCAGCTAATGCAGATT
ACTGTAAGTGATTAATATCTTATGGATATATAATTACATGGTTGACTAATTGCTTGTAATAAATATAGGTCTGGA
AGGACTCTGCGTCTGCCAGGGAGGCAAATGTCATGTTAAAATACCACCAGTTTAG
*Bipolaris victoriae* FI3
ATGCTTCCACTTATCACTGTTGTCTTGGCCTTCCTAGCTAATAGCACCCTCGCTGCTCCGCCTACGAAACCCCTT
CCTTATAGTCCGTCACTGGGATCACAATTCCAAGAGAATCAGGAGAGAAGTATCTGCACTGGCTTACATGTTGGA
AAATTTGGCTGGTGTGGAGGCTGCAAGCGTTTGGAAACTTGCAGCGCGTATCTTGCTTTCCTTTCTGGTTGCACC
GCTACTGAGCTCCATACGTTCCAGTGCCTTGCCCCCCCTCCATGAAGTGGGGTAAACATTGTGTGGACGTTCCAT
GCAGACCAAATTTCTTGTTCGCTGTAGACTAGATCATTGCTAACTCCATATCACAGCTGGCTTGCTTGGCTGCGC
AGGTTCTATCTCATTCGTTAAGGCCAGTGCAGATTATTGTAAGTTATTGCTATGTTGTATATATATATGCTTCAG
CATTTGACTTATTGTTGTCAACACAAGTCTCGAAGGACTCTGCATCTGCCAGGGAGGCCGCTGTAAGACGTAG
*Cercospora canescens* BHU
ATGCGCTTCACGACCTTCCTCTTGCCATTTCTCGGCCTCTCCAAGGCAGCAGTCCTGCCTGTAGATGAAGTGGCA
AGCCCCAACGACAATATATGTACTAAAGCGCACGTTGGAAAGGTGACCACTACGCCACGATGTGATAGTTTTGAT
GAGTTGTTAACGGTTCCTAGTACGGCTGGTGTGCTGGCTGTCGTATGCCTCCCATTGCGATTACAAGTCTCTGGA
ACGGAGTCGACACTAACACCGAGAGCAGAGCGTCTGGAAACATGTACCGCTCTTGGTCTGTACAATGTCCTTAAC
TGCCTCGACGTTGCGACTATTGGGCTTTGCATTGCGCAGACAGCGTTCGGCGTGGCGAATTGGAATTTCTGTAGG
AAACTCATAGCACGCATCGAAATTGTTCTGTGCAGCTAACACGGTCAAGGTGTTGCGGGACTTTGCATCTGTAAG
GGGCCGAATGCGAAGCCCTGCTTTATAAATTGA
*Cercospora janseana* RL44
ATGCATCGCAGCGCAATTTACGCAGCGCTCCTCCTCGTAGCGTCCAATGTCGTGCGCATGTGGGCTGCTCCGATC
CAAGAAGAACAGCTCCAGGCCGTTTCTGCACGAAGCATCTGCACACACTGGCACTTCAGGCATATTGGATGGTGC
ACAGGCTGCACAAGACTCGAGTACCGCACCGCGTTTTTACCCTTCTCATAAGTGAATGCCTAGCG
*Cercospora sojina* 2.2.3
ATGCGCTCCACGACCGTCCTTCTCTCGCTTCTTGGCCTTTCATACGGAGCAGTCTTGGACGTCGATTATCAGAAG
GCGGCCATTGAGCTTGAGGATAGTTTCTGCGGAGGTCTGCACGTCGGAAAGGTAAGAGCTACCAAATACGTTTAT
GGGCGCTGTGGACAACTGATAATGCCTAGGGCGGCTGGTGTGCCGGTTGTCGTAGGTTCTAAACAGCTTCCCAAA
TCTCAGAATTCGAAACATACTTAAAGTCCTGGACAGAGAGACTAGAAAACTGTATCGGGTTTGGTCTATTCAACA
TTATTAATTGCCTGGCTGTCCTAGACAATCCTGTCGCAATTAGAGGTTGTGCTGCGCAGACCGCTTTCTCTCTAG
GCAGTTGGAATTACTGTAAGCATACAAGCGTGTAACTTCGAGGTGTTCCATGCGGCTAATTTAGTACAGGTGCTG
TTGGATTTTGCGTTTGTCAACCTTATGGATCCCAGAAATGCACTATTTCATCATAA
*Cercospora sojina* CCC
ATGCGCTCCACGACCGTCCTTCTCTCGCTTCTTGGCCTTTCATACGGAGCAGTCTTGGACGTCGATTATCAGAAG
GCGGCCATTGAGCTTGAGGATAGTTTCTGCGGAGGTCTGCACGTCGGAAAGGTAAGAGCTACCAAATACGTTTAT
GGGCGCTGTGGACAACTGATAATGCCTAGGGCGGCTGGTGTGCCGGTTGTCGTAGGTTCTAAACAGCTTCCCAAA
TCTCAGAATTCGAAACATACTTAAAGTCCTGGACAGAGAGACTAGAAAACTGTATCGGGTTTGGTCTATTCAACA
TTATTAATTGCCTGGCTGTCCTAGACAATCCTGTCGCAATTAGAGGTTGTGCTGCGCAGACCGCTTTCTCTCTAG

GCAGTTGGAATTACTGTAAGCATACAAGCGTGTAACTTCGAGGTGTTCCATGCGGCTAATTTAGTACAGGTGCTG
TTGGATTTTGCGTTTGTCAACCTTATGGATCCCAGAAATGCACTATTTCATCATAA
*Cercospora sojina* N1
ATGCGCTCCACGACCGTCCTTCTCTCGCTTCTTGGCCTTTCATACGGAGCAGTCTTGGACGTCGATTATCAGAAG
GCGGCCATTGAGCTTGAGGATAGTTTCTGCGGAGGTCTGCACGTCGGAAAGGTAAGAGCTACCAAATACGTTTAT
GGGCGCTGTGGACAACTGATAATGCCTAGGGCGGCTGGTGTGCCGGTTGTCGTAGGTTCTAAACAGCTTCCCAAA
TCTCAGAATTCGAAACATACTTAAAGTCCTGGACAGAGAGACTAGAAAACTGTATCGGGTTTGGTCTATTCAACA
TTATTAATTGCCTGGCTGTCCTAGACAATCCTGTCGCAATTAGAGGTTGTGCTGCGCAGACCGCTTTCTCTCTAG
GCAGTTGGAATTACTGTAAGCATACAAGCGTGTAACTTCGAGGTGTTCCATGCGGCTAATTTAGTACAGGTGCTG
TTGGATTTTGCGTTTGTCAACCTTATGGATCCCAGAAATGCACTATTTCATCATAA
*Cercospora sojina* RACE15
ATGCGCTCCACGACCGTCCTTCTCTCGCTTCTTGGCCTTTCATACGGAGCAGTCTTGGACGTCGATTATCAGAAG
GCGGCCATTGAGCTTGAGGATAGTTTCTGCGGAGGTCTGCACGTCGGAAAGGTAAGAGCTACCAAATACGTTTAT
GGGCGCTGTGGACAACTGATAATGCCTAGGGCGGCTGGTGTGCCGGTTGTCGTAGGTTCTAAACAGCTTCCCAAA
TCTCAGAATTCGAAACATACTTAAAGTCCTGGACAGAGAGACTAGAAAACTGTATCGGGTTTGGTCTATTCAACA
TTATTAATTGCCTGGCTGTCCTAGACAATCCTGTCGCAATTAGAGGTTGTGCTGCGCAGACCGCTTTCTCTCTAG
GCAGTTGGAATTACTGTAAGCATACAAGCGTGTAACTTCGAGGTGTTCCATGCGGCTAATTTAGTACAGGTGCTG
TTGGATTTTGCGTTTGTCAACCTTATGGATCCCAGAAATGCACTATTTCATCATAA
*Cercospora sojina* S9
ATGCGCTCCACGACCGTCCTTCTCTCGCTTCTTGGCCTTTCATACGGAGCAGTCTTGGACGTCGATTATCAGAAG
GCGGCCATTGAGCTTGAGGATAGTTTCTGCGGAGGTCTGCACGTCGGAAAGGTAAGAGCTACCAAATACGTTTAT
GGGCGCTGTGGACAACTGATAATGCCTAGGGCGGCTGGTGTGCCGGTTGTCGTAGGTTCTAAACAGCTTCCCAAA
TCTCAGAATTCGAAACATACTTAAAGTCCTGGACAGAGAGACTAGAAAACTGTATCGGGTTTGGTCTATTCAACA
TTATTAATTGCCTGGCTGTCCTAGACAATCCTGTCGCAATTAGAGGTTGTGCTGCGCAGACCGCTTTCTCTCTAG
GCAGTTGGAATTACTGTAAGCATACAAGCGTGTAACTTCGAGGTGTTCCATGCGGCTAATTTAGTACAGGTGCTG
TTGGATTTTGCGTTTGTCAACCTTATGGATCCCAGAAATGCACTATTTCATCATAA
*Curvularia geniculata* P1
ATGCTGCCACGAATGATAATTATCATGGCCTTCCTGGCTAACAGGACTCTCGCTGCTCCGTCTACGAATATGATA
GCTCATGATTCTCTAGTGGAACCTCAATTTCAACATAATCAGGAGAGAAGCATCTGCACTGGCTTTCATATCGGC
AAATTTGGCTGGTGTGGAGGTTGTAAGCGTTTTGAAAGTTGCAGCGCATTCCTTGCTGGCCTAGCTGCCTGCCCC
TCTCTTTCCGTCGGTATGTTTTAGTGCCCCAACCCTACCTTAATTGCTAACTCTGTTTCGTAGCTGGCGTACTTA
CCTGCGCATCGTCTGCAGGAATCCTTGGGTCTCGCGCAGATTATTGTAAGTGGTTATTATTTGATGTATATATAC
ATGCTTCAGCACTTGATTAATGATTGCAAATCCAGGTCTAGAAGGACTCTGCATCTGCCAGGGAGGCCACTGCAA
GACTTAG
*Curvularia geniculata* W3
ATGCTGCCACGAATGATAATTATCATGGCCTTCCTGGCTAACAGGACTCTCGCTGCTCCGTCTACGAATATGATA
GCTCATGATTCTCTAGTGGAACCTCAATTTCAACATAATCAGGAGAGAAGCATCTGCACTGGCTTTCATATCGGC
AAATTTGGCTGGTGTGGAGGTTGTAAGCGTTTTGAAAGTTGCAGCGCATTCCTTGCTGGCCTAGCTGCCTGCCCC
TCTCTTTCCGTCGGTATGTTTTAGTGCCCCAACCCTACCTTAATTGCTAACTCTGTTTCGTAGCTGGCGTACTTA
CCTGCGCATCGTCTGCAGGAATCCTTGGGTCTCGCGCAGATTATTGTAAGTGGTTATTATTTGATGTATATATAC
ATGCTTCAGCACTTGATTAATGATTGCAAATCCAGGTCTAGAAGGACTCTGCATCTGCCAGGGAGGCCACTGCAA
GACTTAG
*Curvularia hawaiiensis* PK9021
ATGCAGCCACTTATCACTATCGCCTTGGCCCTCCAGACCACTACTACTCTCGCTGTTCCAGTCATAGATGCCAAG
CTTTCAGGCACTGATCTCGCTATCCCGCTCACTGATATCGCCGCCCAGAGCACTGATCTCACTGATCTGCAAGCT
CGCGATTCTTCACTAGCACCACGAGGTATCTGCACGCAATTTAAAATTGGCAAATATGGCTGGTGTGGAGGTTGC
AAGCGCTTTGAAACTTGCAGCGCATATCTTGCCTTCATGTCTGGATGCGCCTCTTTTACCCCTAGTATGTTATCT
CCTATGAGAAGAAAGCAGCGCCATGATTGAGATCGAGAGCTAGGCAGAATAATTGCTAACTTTGTTACACAGCTG
GTGTGCTTGCTTGCGTAGGGGCTTCTGGATTTCTTGCAGCTAATGCAGATTACTGTAAGTGATCAATATCTTATG
GATATATGTAGTTAAATGGTTGACTGATTGCTTGTAACAAATATAGGTCTGGAAGGACTCTGCGTCTGCCAGGGA
GGCAAATGCCATGTTAAAATACCACCAGCTTA
*Curvularia lunata* Cl-Gvt
CAGTTCTGTTATAATCCATATGTTTTAGTGGCTTCCCCACTCCCAAGTTAAAGGCAAATATTGTGTATATCTTTC
ATAGAGACTCAATATCTCGTTCGTT
*Curvularia lunata* CX-3
ATGCTACCACGAATGATGATTATCGTGGCCTTCCTGGCTAACAGGACTCTCGCTGCTCCGTCTACGAATATGATA
GCTCATGATTCTCTAGTGGAATCTCCATTCCAACAGAATCAAGAGAGAAGTATCTGCACTGGCTTTCATATCGGC
AAATTTGGCTGGTGTGGAGGTTGTAAGCGTTTTGAAAGTTGCAGCGCATTCCTTGCTGGCCTAGCTGCCTGCCCC
TCTCTTTCCGTCGGTATGTTTTAGTGCCCCAACCCTACCTTAATTGCTGACTCTGTTGCGTAGCTGGCGTACTTA
CCTGCGCATCTTCTGCAGGAATCCTTGGGTCTCGCGCAGATTATTGTAAGTGGTTACTATTTGATGTATATATGT

ATGCTTCAACACTTGATTAATTATTGCAAATACAGGTCTAGAAGGACTCTGCATCTGCCAGGGAGGCCACTGCAC
GACTTAG

*Curvularia lunata* W3

ATGCTGCCACGAATGATAATTATCATGGCCTTCCTGGCTAACAGGACTCTCGCTGCTCCGTCTACGAATATGATA
GCTCATGATTCTCTAGTGGAACCTCAATTTCAACATAATCAGGAGAGAAGCATCTGCACTGGCTTTCATATCGGC
AAATTTGGCTGGTGTGGAGGTTGTAAGCGTTTTGAAAGTTGCAGCGCATTCCTTGCTGGCCTAGCTGCCTGCCCC
TCTCTTTCCGTCGGTATGTTTTAGTGCCCCAACCCTACCTTAATTGCTAACTCTGTTTCGTAGCTGGCGTACTTA
CCTGCGCATCGTCTGCAGGAATCCTTGGGTCTCGCGCAGATTATTGTAAGTGGTTATTATTTGATGTATATATAC
ATGCTTCAGCACTTGATTAATGATTGCAAATCCAGGTCTAGAAGGACTCTGCATCTGCCAGGGAGGCCACTGCAA
GACTTAG

*Curvularia papendorfii* UM 226

ATGCTGCCACGAATGATAATTATCATGGCCTTCCTGGCTAACAGGAGTCTCGCTGCTCCGTCTACGAATATGATG
GCTCATGATTCTCTAGTGGAACCTCAATTTCAACATAATCAGGAGAGAAGCATCTGCACTGGGTTTAATATCGGC
AAATTTGGCTGGTGTGGAGGTTGTAAGCGTTTTGAAAGTTGCAGCGCATTCCTTGCTGGCCTAGCTGCCTGCCCT
TCTCTTTCCGTCGGTGTGTTTTAGTGCCCCAACCCTACCTTAATTGCTAACTCTGTTTCGTAGCTGGCGTACTTA
CCTGCGCATCTTCTGCAGGAATCCTTGGGTCTCGCGCAGATTATTGTAAGTGGTTATTATTTGATGTATATATAC
ATGCTTCAGCACTTGATTAATGATTGCAAATACAGGTCTAGAAGGACTCTGCATCTGCCAGGGAGGCCACTGCAA
GACTTAG

*Curvularia spicifera* FR9030

ATGAAGCTATCTAGCATCAGTCTCCCTATCCTTCACCTAGCTTGCCTTGTTGCTGGACAAAAGGTCCCAGAACCA
GCTAATCCACCACCTGAACCGGCAAACAATAAACACTGTGTTATTCCTGCGAATGTTGGATCTGTCTGTGAAAAC
TGGCATCACACGTTCAACAAACAGATCACAATTGGATGGTGCGCAGGCTGCAAGCGACTTGAAAGATGTGGTTAT
CTTTTAACAGTTGAGGCTGCTGCCTGTATTGCAGGATTGATACTTTGGGAAACAAGTCGGTTTAGAATCCATTTC
TCCATACACAGGCACTAATGAAATTTAGAGGCTGTTCTTGGGTGTCTGCTGGAAGCAGCTGCAGACGATTTCGCG
ACATACTGTAAGTTTCCAATCTTTAGTCTTTGAGCGCAAACAAGAGGTTAATTGTTTTTAGGTCTTGATGGTGTC
TGCTAG

*Curvularia spicifera* FR9030

ATGCAGCCACTTATTACTATCGCTTTGGCCCTCCAGGTCGCTACTACTATCGCTATCCCAGTCACTGATGTCACG
GTTAAGGACATTGATCTCACGCTTTCAGGCACTGATCTCACGCTTGCGGGCACTGATCTCGCTATCCCGCTCGCC
GATCTCGCTGCCCCGGGCACTGATCTCACTGACGTGCAGGCTCGTGATTCCTCACTAGAATCACGAAGTATTTGC
ACCGGATTTCATGTTGGCCGATATGGCTGGTGTGGAGGTTGCAGGCGCTTGGAAGCTTGCGGCGCGTATCTTGGT
TTCTTGTCTGGATGCAGCTCTCTTAGCGTTGGTATGTTTTCCCCCATAAAAAAGCAGCACCATAATTGCGATAGA
CAGACCAGGCAGAAAAAACATGTTATTCTTGGTAGACTGGATAATTGCTGACTTTGTTGCACAGCTGGTGTGGTT
GGTTGCGCAGCTGCGTCTGGAGTCCTTGCAGCTAATGCAAATTATTGTAAGTAATAAACATCTTATGTATATATA
TGTGCTAGGATACTTGATTAATTATTGCCCACAAATATAGGCCTGGAAGGACTTTGCGTTTGCCAGGGAGGCAAA
TGCCATGTTTGA

*Curvularia* sp ZM96

ATGCAGCCACTTATTACTATCGCTTTGGCCCTCCAGGTCGCTACTACTATCGCTATCCCGGTCACTGATGTCACG
GTTAAGGACATTGATCTCACGCTTTCAGGCACTGATCTCACGCTTGCGGGCACTGATCTCGCTATCCCGCTCGCC
GATCTCGCTGCCCCGGGCACTGATCTCACTGACGTGCAGGCTCGTGATTCCTCACTAGAATCACGAAGTATCTGC
ACCGGATTTCATATTGGCCGATATGGCTGGTGTGGAGGTTGCAGGCGCTTGGAAGCTTGCGGCGCGTATCTTGGT
TTCTTGTCTGGATGCAGCTCTCTTAGCGTTGGTATGTTTTCCCCCATAAAAGCAGCACCATAATTGCGATAGACA
GACCAGGCAGAAAAAACATGTTATTCCTGGTAGACTGGATAATTGCTGACTTTGTTGCACAGCTGGTGTGGTTGG
TTGCGCAGCTGCGTCTGGAGTCCTTGCAGCTAATGCAAATTATTGTAAGTAATAAACATCTTATGTATATATATG
TGCTAGGATACTTGATTAATTATTGCCCACAAATATAGGCCTGGAAGGACTTTGCGTCTGCCAGGGAGGCAAATG
CCATGTTTGA

*Exserohilum rostratum* B6094

ATGGTCGCCATTCACGCCCTCACCTTTGCCTTCCTGACCAGCAGTATTCTTGCTGCTCCGCCTACCGATCTCAAA
ACCCGTGATTTCCCAGTGGAATTGGAAGTCGCACAGAACGAAGAAAGAAGCGTCTGTACCGGCTTCAACATTGGA
AAGTTTGGATGGTGCGGAGGTTGCAAGCGTCTTGAGACTTGCTCCGCTTATCTTGCTTTCCTCTCTGGATGCACC
AGTGTTTCTGTCGGTATGTTTTAGTGCTCTTCACTCCTCCCTCTCCTCTTCCAAAGCTGGAAGATTGGATGTCTT
GTTGGTGGTGTTGAGAAACGTCATTGCTAACTTTATCGTGTAGTTGGTGTTCTTGGCTGCGCAGGTGCGGTCGGA
TTCCTCGGTGCAAACGCAGACTACTGTAAGTGATTATCATTGTATGCTCGAACACTTTGTTGAGTATCGCTAACA
AAAACAGGTCTGGAAGGACTTTGCGTCTGCCAGGGAAACCACTGCAAGACTTAG

*Exserohilum rostratum* B6096

ATGGTCGCCATTCACACCATCACCCTTGCCTTCCTGGCCAGCAGCGTTCTCGCTGCTCCCCCCACTGATCTCAAA
ACCCGTGATTTCCCAGTCGAATTGGAAGTCGCACAGAACGAAGAAAGAAGCGTCTGTACCGGCTTCAACATTGGA
AAGTTTGGATGGTGCGGAGGTTGCAAGCGTCTTGAGACTTGCTCCGCTTATCTTGCTTTCCTGTCTGGATGCACC
AGTGTTTCTGTCGGTATGTTTTAGTGCCCTTCACTTCTCCCTCTCCTCTTCCAAAGCTGGAAGGCTGGATGTCTT
GTTGGTGGTGTTGAGAAACATCATTGCTAACTTTATCGTGTAGTTGGTGTTCTTGGCTGCGCAGGTGCGGTCGGA

TTCCTCGGTGCAAACGCAGACTACTGTAAGTGATTGTCATTGTATGCTCGAACACTTTATTGAGTATCGCTAACA
AAAACAGGTCTGGAAGGACTTTGCGTCTGCCAGGGAAACCACTGTAAGACTTAG

*Exserohilum rostratum* B6171

ATGGTCGCCATTCACACCATCACCCTTGCCTTCCTGGCCAGCAGCGTTCTCGCTGCTCCCCCCACTGATCTCAAA
ACCCGTGATTTCCCAGTCGAATTGGAAGTCGCACAGAACGAAGAAAGAAGCGTCTGTACCGGCTTCAACATTGGA
AAGTTTGGATGGTGCGGAGGTTGCAAGCGTCTTGAGACTTGCTCCGCTTATCTTGCTTTCCTGTCTGGATGCACC
AGTGTTTCTGTCGGTATGTTTAGTGCCCTTCACTTCTCCCTCTCCTCTTCCAAAGCTGGAAGGCTGGATGTCTT
GTTGGTGGTGTTGAGAAACATCATTGCTAACTTTATCGTGTAGTTGGTGTTCTTGGCTGCGCAGGTGCGGTCGGA
TTCCTCGGTGCAAACGCAGACTACTGTAAGTGATTGTCATTGTATGCTCGAACACTTTATTGAGTATCGCTAACA
AAAACAGGTCTGGAAGGACTTTGCGTCTGCCAGGGAAACCACTGTAAGACTTAG

*Exserohilum rostratum* B6177

ATGGTCGCCATTCACACCATCACCCTTGCCTTCCTGGCCAGCAGCGTTCTCGCTGCTCCCCCCACTGATCTCAAA
ACCCGTGATTTCCCAGTCGAATTGGAAGTCGCACAGAACGAAGAAAGAAGCGTCTGTACCGGCTTCAACATTGGA
AAGTTTGGATGGTGCGGAGGTTGCAAGCGTCTTGAGACTTGCTCCGCTTATCTTGCTTTCCTGTCTGGATGCACC
AGTGTTTCTGTCGGTATGTTTAGTGCCCTTCACTTCTCCCTCTCCTCTTCCAAAGCTGGAAGGCTGGATGTCTT
GTTGGTGGTGTTGAGAAACATCATTGCTAACTTTATCGTGTAGTTGGTGTTCTTGGCTGCGCAGGTGCGGTCGGA
TTCCTCGGTGCAAACGCAGACTACTGTAAGTGATTGTCATTGTATGCTCGAACACTTTATTGAGTATCGCTAACA
AAAACAGGTCTGGAAGGACTTTGCGTCTGCCAGGGAAACCACTGTAAGACTTAG

*Exserohilum rostratum* B6207

ATGGTCGCCATTCACACCATCACCCTTGCCTTCCTGGCCAGCAGCGTTCTCGCTGCTCCCCCCACTGATCTCAAA
ACCCGTGATTTCCCAGTCGAATTGGAAGTCGCACAGAACGAAGAAAGAAGCGTCTGTACCGGCTTCAACATTGGA
AAGTTTGGATGGTGCGGAGGTTGCAAGCGTCTTGAGACTTGCTCCGCTTATCTTGCTTTCCTGTCTGGATGCACC
AGTGTTTCTGTCGGTATGTTTAGTGCCCTTCACTTCTCCCTCTCCTCTTCCAAAGCTGGAAGGCTGGATGTCTT
GTTGGTGGTGTTGAGAAACATCATTGCTAACTTTATCGTGTAGTTGGTGTTCTTGGCTGCGCAGGTGCGGTCGGA
TTCCTCGGTGCAAACGCAGACTACTGTAAGTGATTGTCATTGTATGCTCGAACACTTTATTGAGTATCGCTAACA
AAAACAGGTCTGGAAGGACTTTGCGTCTGCCAGGGAAACCACTGTAAGACTTAG

*Exserohilum rostratum* B6272

ATGGTCGCCATTCACGCCCTCACCTTTGCCTTCCTGGCCAGCAGTGTTCTTGCTGCTCCCCCTACCGATCTCAAA
ACCCGTGATTTCCCAGTGGAATTGGAAGTCGCACAGAACGAAGAAAGAAGCGTCTGTACCGGCTTCAACATTGGA
AAGTTTGGATGGTGCGGAGGTTGCAAGCGTCTTGAGACTTGCTCCGCTTATCTTGCTTTCCTCTCTGGATGCACC
AGTGTTTCTGTCGGTATGTTTAGTGCTCTTCACTCCTCCCTCTCCTCTTCCAAAGCTGGAAGACTGGATGTCTT
GTTGGTGGTGTTGAGAAACATCATTGCTAACTTTATCGTGTAGTTGGTGTTCTTGGCTGCGCAGGTGCGGTTGGA
TTCCTTGGTGCAAACGCAGACTACTGTAAGTGATTATCATTGTATGCTCGAACACTTTGTTGAGTATCGCTAACA
AAAATAGGTCTGGAAGGACTTTGCGTCTGCCAGGGAAACCACTGCAAGACTTAG

*Exserohilum rostratum* B6284

ATGGTCGCCATTCAAGCCATCACCTTTGCCTTCCTGGCCAGCAGCGTTCTCGCTGCTCCCCCTACGGATCTCAAA
ACCCGTGATTTCCCAGTAGAATTGGAAGTCGCACAGAACGAAGAAAGAAGCGTCTGTACCGGCTTCAACATTGGA
AAGTTTGGATGGTGCGGAGGTTGCAAGCGTCTTGAGACTTGCTCCGCTTATCTTGCTTTCCTGTCTGGATGCGCC
AATATTTCTGTCGGTATGTTTAGTGACCTTACCTCCTCTTCCAAAGCTGGCAGACCGGATGTCTTGTTGGTGGT
GGTAAAAAAACATCATTGCTAACTTTATCGTGTAGTTGGTGTTCTTGGCTGCGCAGGTGCGGCCGGATTCCTTGG
TGCAAATGCAGACTACTGTAAGTGATTATCATTGTATGCTCGAACACTTTGTTGAGTATCGCTAACAAAAACAGG
TCTGGAAGGACTTTGCGTTTGCCAGGGAAACCACTGCAAGACTTAG

*Exserohilum rostratum* B9826

ATGGTCGCCATTCACACCATCACCTTTGCCTTCCTGGCCAGCAGCGTTCTCGCTGCCCCCCCTACGGATCTCAAA
ACCCGTGATTTCCCAGTCGAATTGGAAGTCGCACAGAATGAAGAAAGAAGCGTCTGTACCGGCTTCAACATTGGA
AAGTTTGGATGGTGCGGAGGTTGCAAGCGTCTTGAGACTTGCTCCGCTTATCTTGCTTTCCTCTCTGGATGCACC
AGTGTTTCTGTCGGTATGTTTAGTGCTCTTCACTCCTCCCTCTCCTCTTCCAAAGCTGGAAGACTGGATGTCTT
GTTGGTGGTGTTAAGAAACATCATTGCTAACTTTATCGTGTAGTTGGTGTTCTTGGCTGCGCAGGTGCGGTTGGA
TTCCTTGGTGCAAACGCAGACTACTGTAAGTGATTATCATTGTATGCTCGAACACTTTGTTGAGTATCGCTTAAC
AAAAATAGGTCTGGAAGGACTTTGCGTCTGCCAGGGAAACCACTGCAAGACTTAG

*Exserohilum rostratum* BF9006

ATGGTCGCCATTCACGCCCTCACCTTTGCCTTCCTGGCCAGCAGTGTTCTTGCTGCTCCCCCTACCGATCTCAAA
ACCCGTGATTTCCCAGTGGAATTGGAAGTCGCACAGAACGAAGAAAGAAGCGTCTGTACCGGCTTCAACATTGGA
AAGTTTGGATGGTGCGGAGGTTGCAAGCGTCTTGAGACTTGCTCCGCTTATCTTGCTTTCCTCTCTGGATGCACC
AGTGTTTCTGTCGGTATGTTTAGTGCTCTTCACTCCTCCCTCTCCTCTTCCAAAGCTGGAAGACTGGATGTCTT
GTTGGTGGTGTTAAGAAACATCATTGCTAACTTTATCGTGTAGTTGGTGTTCTTGGCTGCGCAGGTGCGGTTGGA
TTCCTTGGTGCAAACGCAGACTACTGTAAGTGATTATCATTGTATGCTCGAACACTTTGTTGAGTATCGCTAACA
AAAATAGGTCTGGAAGGACTTTGCGTCTGCCAGGGAAACCACTGCAAGACTTAG

*Exserohilum rostratum* ER1

ATGGTCGCCATTCACGCCATCACCTTTGCCTTCCTGGCCAGCAGCGTTCTCGCTGCTCCCCCTACGGATCTCAAA
ACCCGTGATTTCCCAGTAGAATTGGAAGTCGCACAGAACGAAGAAAGAAGCGTCTGTACCGGCTTCAACATTGGA

AAGTTTGGATGGTGCGGAGGTTGCAAGCGTCTTGAGACTTGCTCCGCTTATCTTGCTTTCCTGTCTGGATGCGCC
AATATTTCTGTCGGTATGTTTTAGTGACCTTACCTCCTCTTCCAAAGCTGGCAGACCGGATGTCTTGTTGGTGGT
GGTAAAAAACATCATTGCTAACTTTATCGTGTAGTTGGTGTTCTTGGCTGCGCAGGTGCGGCCGGATTCCTTGGT
GCAAATGCAGACTACTGTAAGTGATTATCATTGTATGCTCGAACACTTTGTTGAGTATCGCTAACAAAAACAGGT
CTGGAAGGACTTTGCGTTTGCCAGGGAAACCACTGCAAGACTTAG

*Exserohilum rostratum* LWI
ATGGTCGCCATTCACGCCCTCACCTTTGCCTTCCTGGCCAGCAGTATTCTTGCTGCTCCGCCTACCGATCTCAAA
ACCCGTGATTTCCCAGTGGAATTGGAAGTCGCACAGAACGAAGAAAGAAGCGTCTGTACCGGCTTCAACATTGGA
AAGTTTGGATGGTGCGGAGGTTGCAAGCGTCTTGAGACTTGCTCCGCTTATCTTGCTTTCCTCTCTGGATGCACC
AGTGTTTCTGTCGGTATGTTTTAGTGCTCTTCACTCCTCCCTCTCCTCTTCCAAAGCTGGAAGATTGGATGTCTT
GTTGGTGGTGTTGAGAAACGTCATTGCTAACTTTATCGTGTAGTTGGTGTTCTTGGCTGCGCAGGTGCGGTCGGA
TTCCTCGGTGCAAACGCAGACTACTGTAAGTGATTATCATTGTATGCTCGAACACTTTGTTGAGTATCGCTAACA
AAAACAGGTCTGGAAGGACTTTGCGTCTGCCAGGGAAACCACTGTAAGACTTAG

*Exserohilum rostratum* M 18 0203
ATGGTCGCCATTCACGCCCTCACCTTTGCCTTCCTGGCCAGCAGTGTTCTTGCTGCTCCCCCTACGGATCTCAAA
ACCCGTGATTTCCCAGTCGAATTGGAAGTCGCACAGAATGAAGAAAGAAGCGTCTGTACCGGCTTCAACATTGGA
AAGTTTGGATGGTGCGGAGGTTGCAAGCGTCTTGAGACTTGCTCCGCTTATCTTGCTTTCCTCTCTGGATGCACC
AGTGTTTCTGTCGGTATGTTTTAGTGCTCTTCACTCCTCCCTCTCCTCTTCCAAAGCTGAAAGACTGGATGTCTT
GTTGGTGGTGTTGAGAAACATCATTGCTAACTTTATCGTGTAGTTGGTGTTCTTGGCTGCGCAGGTGCGGTCGGA
TTCCTCGGTGCAAACGCAGACTACTGTAAGTGATTATCATTGTATGCTCGAACACTTTGTTGAGTATCGCTAACA
AAAACAGGTCTGGAAGGACTTTGCGTCTGCCAGGGAAACCACTGTAAGACTTAG

*Exserohilum rostratum* SR-KPL1
ATGGTCGCCATTCACACCATCACCTTTGCCTTCCTGGCCAGCAGCGTTCTCGCTGCCCCCCCTACGGATCTCAAA
ACCCGTGATTTCCCAGTCGAATTGGAAGTCGCACAGAATGAAGAAAGAAGCGTCTGTACCGGCTTCAACATTGGA
AAGTTTGGATGGTGCGGAGGTTGCAAGCGTCTTGAGACTTGCTCCGCTTATCTTGCTTTCCTCTCTGGATGCACC
AGTGTTTCTGTCGGTATGTTTTAGTGCTCTTCACTCCTCCCTCTCCTCTTCCAAAGCTGGAAGACTGGATGTCTT
GTTGGTGGTGTTAAGAAACATCATTGCTAACTTTATCGTGTAGTTGGTGTTCTTGGCTGCGCAGGTGCGGTCGGA
TTCCTTGGTGCAAACGCAGACTACTGTAAGTGGTCATCATTGTATGCTCGAACTCTTCGTTGAGTATCGCTAACA
ACAACAGGTTTGGAAGGACTTTGCGTCTGCCAGGGAAACCACTGCAAGACTTAG

*Exserohilum rostratum* ZM170581
ATGGTCGCCATTCACGCCATCACCTTTGCCTTCCTGGCCAGCAGCGTTCTCGCTGCTCCCCCTACGGATCTCAAA
ACCCGTGATTTCCCAGTAGAATTGGAAGTCGCACAGAACGAAGAAAGAAGCGTCTGTACCGGCTTCAACATTGGA
AAGTTTGGATGGTGCGGAGGTTGCAAGCGTCTTGAGACTTGCTCCGCTTATCTTGCTTTCCTGTCTGGATGCGCC
AATATTTCTGTCGGTATGTTTTAGTGACCTTACCTCCTCTTCCAAAGCTGGCAGACCGGATGTCTTGTTGGTGGT
GGTAAAAAACATCATTGCTAACTTTATCGTGTAGTTGGTGTTCTTGGCTGCGCAGGTGCGGCCGGATTCCTTGGT
GCAAATGCAGACTACTGTAAGTGATTATCATTGTATGCTCGAACACTTTGTTGAGTATCGCTAACAAAAACAGGT
CTGGAAGGACTTTGCGTTTGCCAGGGAAACCACTGCAAGACTTAG

*Pseudocercospora cruenta* Pscow-1
ATGCATCCATTCATCATGATCCTGTCTCTTCACGCAGTGACCACTGTGCTCGCCATGCCCAACCCTGTTGCTGAG
CCAAATGCTCTTGCTTTGCCTCTTGCCCTCCCGTTCATCGAAAACATTGACGGTTCAGACAAGCACAAGAAGCAC
AAGCATAAGCAGAACGAAGAAGACATCGAGGATCAGGAAGAGCAGAAGTCGACAAAGCAGAAGCATAAGCAGCAA
GATGATGATGATGACGATGACTCCAAGAAGGACTCAGTCAGCAACGAAGACTTCAGCCTACCAAAGTGCCCACTA
AGCATTTTTAAAAAGGCTGTGCGCGGCTCACGACATGATACACAGCGACCGACTGCTAACCGTCTCTTCGCAGGG
CTGGTGCGGCGGATGTCAAAGACTTGAAACTTGTGTTGGTACAATCGGATTCTTTGCGGCTTCGTGCCTTGCTGG
GTCTCTTACTGTTGGTAAGCATGTCATCTATCGCGCGCTTATGACGAGAAATTGACAGGATTTTGTAGCCGGGGC
ACTGTCCTGCTTTGGCTCGGCTGGCTTTGCGGCTGGTGAATGGAATTACTGTGAGTCAAGCGCATCTATTGATGG
AAGCTATCTCACTGACACTTTCCAAATAGGCGTCGATGGAGTTTGTGCGTGCCAGGGCAATACCAAATGCCATAA
GTGA

*Pseudocercospora eumusae* CBS 114824
ATGCATTCATCATTCCTCTGGATCCTATCTCTTCAATCCGTGGTAACTGCCATGCCCAATCCAACGGCCGTTGCC
GCAGACAGCATATGTCCAACAGCACTTTTCCACAAAGCTGTAAGTTGCACGTTACGTAGCACGGAAGAATAGAAC
TCGCTGATTCTTCGTTAGGGCTGGTGCGGCGGATGTCAAAGACTCGAGACTTGCATTGGATCACTGGGATTCTTT
GCGTCTTCGTGCATTGCTGGGTCTCTGACGACTTGTGAGACATGGCGTGCGCTCGTGTCGAGAGGATTTTTTTTT
TGACTGACGGGATTGCTTTGTATAGCTAGTGTGCTGTCTTGCATTGGTTCGACTGGCTTTGCGGCTGGGACTTGG
AATTATTGTAAGTGGAATCCCCCATCAGAGAGAGAGAGAGAGACAGAGAGACAGCGGTGATTAATTGGTCGACTT
TGAATAGGTGTCGATGGAGTTTGTGCATGCCAGGGAAATACCCAATGCCATATCTAG

*Pseudocercospora fuligena* PF001
ATGCATCCATTGATTATCATCGTGTCTCTTCACGCAGCGAGTAATGTGCTCGCCATGCCCAACCCCGTTGCCGAG
CCAAATGCTCTTGCTTTACCTCATGCTCTGCCAATGATCGAGGAGATCGATGGATCCGATAAGCACAAGAAGCAC
AAACAGAAGCAAAACGAAGAAGACATTGGTGACGAGGAAGAGCAGAAGTCGACAAAGCATAAGCATAAGCAACAC
GACGATCACGATGACGATGATTCCAAGAAGGATTCAGTCAGCGACGAAGACTTTAGCTTTCATCTTCCAAAATGT

CCGACATCTACCCTGAACCTGGCTGTAAGTTGCACGCAGCATGAACCAGAGTCACTGCTTGCTGAGTCTCTCTTC
ATAGGGCTGGTGCGGCGGATGCCGAAGACTTGAGACCTGTCTTGGCACGCTAGGATTCTTGACGGCTTCGTGCCT
TTTCGGATCTGTTACTATCGGTATGCATATCGCAGATTGTGTGCTTCAATTCAGTAATCGACTAATTCTTATAGG
CGGTGTACTTAGTTGTTTCGGCTCGGTGGGCTTTGCGACTGGAGAATGGAACTACTGTAAATCAAGCGCGTTCAT
TGATTGAAGCTGTCCCACTGACTCTCTCGAACAGGCGTCGATGGTGTTTGTGCGTGCCAAGGCAAGCCAGAATGC
CATAAGTGA

*Pseudocercospora musae* CBS 116634
ATGCATCTATTCCAATGGATCTTGTGTCTTCACTCGGCGACTGCCGTCCTCGCCATGCCCAATCCTGTTGCCGAG
CCAAATGCTCTTGCTTTGCCTCTTGCCCACTCAATGGTTGAAGAAAAGTCCGCAAAGCATAAGGACAAGCGAGAA
GATGATCATGATCAAGGCGCGGAAGGGGATCCAGTCAGCAACACAGACATCAGCATAAAGTGTCCGCTTTCACTT
TTTTCCAAGGCTGTAAGTTGCATGCAAAATGAAGCCAAGGGACGTATGACTGCTGACGGTCAGTCTTATAGGGCT
GGTGCGGTGGATGCTCAAGACTTGAGACTTGTGTTGGAACTCTAGGGTTCTTCGTGTCTTCGTGCCTTGTTGGGT
CAGTTACTATCGGTATGGTTATCACATGTCGTGTGCTTGGAATGAGAAATTGACTCGCTACCATAGCCGGTGTAG
CCGCCTGTGCAGGATCGACTGGCTTTGCAGCTGGGAGTTGGAATAATTGTATGTCAAGCGTGCTCATCGACAGAA
GCTATCCCACTGACTCTCTCGAATAGGCGTTGATGGTCTTTGTTTGTGCCAGGGCAACACCAAGTGCCACATATG
A

*Pseudocercospora pini-densiflorae* CBS 125139
ATGCATCCATTCATCTGGATCCTGTCTCTTCACTCGGTTACCACCGTGCTCGCCATGCCCAACCCTGTTGCAGAA
CCAAATGCTCTTGCTTTGCCTCTTGCTCTACCACTGATTGAGGAGATTGATGAATCCGACAAGCACAAGAAGCAA
AAGCATAAGCAAGACAAAGAAGACATTGACGCAGAGGAGGGGCAGAAGTCGACAAAGCATAAGCACAAGCAGCAA
GATGATGATGATGACGATGACTCCAAGAAGGACTCAGTCAGCAACGAAGACTTCAGCATTCATCTTCCAAAATGT
CCGACATCTATTCTGAACAAGGCTGTAAGTTGCCCGCAGCATGAACTAGAGCCACTGCTTGCTGACTCTCTCTTC
ACAGGGCTGGTGCGCCGGATGCCAAAGACTTGAGACCTGTCTTGGCACGTTGGGATTCTTTGCAGCTTCGTGCCT
TGCTGGGTCTCTTACTGTTGGTGAGCATGTGACGTTTCGTGCGCTCTTGACGAGAAGTTGACCAGATTTTACAGC
TGGCGCACTGTCTTGCTTCGGCTCGACGGGCTTCGCAGCTGGGGAATGGAATTATTGTAAGTCAAGCGCATCTAT
TGATGGAAGCTGTCCCACTGACAATCCCCAACAGGCGTCGATGGTGTTTGTGCATGCAAGGGCAATACCAAATGC
CATAAGTAA

### Group 3 gene sequences

*Pseudocercospora macadamiae* BRIP 55526
ATGCACTTCCCTCGCCTCCTCACCATCATGTGCTACAGCTCAATCACAATGGGCCATGTCACATGGAAAACCTGC
CCCAATCCTGCCTGGTACATAGACGAAGACATCACGCCTTCTGACTCCGACCTGAACGTCCGTGGCACAGACAAG
CACGACCCACAAAGCGTCGATCCACCAACTCAGCATTCTCACAGCCATCATCTAAAGACGCAGAAACTCCAGATT
ATCACCAATGGTGAAGCTTCATCCCACGCCAAAGCAGCAATCAAAAAAGCCGCAGCGGCCGGTACAGTCGAAGCA
GATTTATCATTATGCACACATCTCCATATCGGGAAATTCGTAAGTCCCTAATACCTCCTCCTAAAATTCCTCATC
ACCCCAAACTGAATTCTTCTTCTCTTTTCTAGGGCTGGTGCGCAGGCTGTCGCCGCCTAGAAAAATGTATCGGAG
TAGGATTCTTCGTGATCGGAGGCTTACTCGCATGGATAGCAAGTTGTGTTCTTACAGGCGGAGCCAGTGCGACTA
CGATCGCAGCATTTATAAGTTTTCTGGGATCGATTGGGTTCTCGGTTAGTACGTTTAATGATTGTATTATTGGGC
TTTGTGCTTGTCAACCTGGGAGGAAGGGGAGTCATAAATGTACTTTGTAG

*Pseudocercospora ulei* ERN8
ATGCGCATTCTCGCTATCCTGTACACCATTAGTAGCCTCTTGATCTTGTGCTACGCAACACCAGCGACCTCGCCT
AACTTTAACTGGTATCGGGAAAAGAGAGTAACGCCCCACTTCAGCCTAAACAGCCGCGATGCCGACTTTTATAAT
ACGGAAGTCACCTTCAATCCCCAGCCAGCTCACCGCGCTATTCAAGAGGCCAAAAGAGTCGGCACTATCGAAGCC
GACTTGAGTATGTGCAGACATATTGTCCTCTTTGGCAGGCATGTAAGTCCCTCTCCTACATCGAGCACCGGCCCC
TTGCTACACTTCTAACTATCTTCCTCTAGATATGGTGTGCAGGCTGCAAGAGACTCGCGAAGTGTATTGGCACGG
GATGTTTCGGACTAGCGGGCATGGGCGCTCTAATAGGTCTGATTTTTATGACAGACGGAGCTATATTGCCGATCA
TTGTAGCTTATGTTAACGGTATTGGGGGTCTCTCTATGGGCGTTGCTTCGTTTGCTGATTGCCTGGATGGCCTCT
GCCCTTGCAAACACGGCAGATGTGTGGGTGATGTACATGGTAGTGAATAA

*Pseudocercospora ulei* GCL012
ATGCGCATTCTCGCTATCCTGTACACCATTAGTAGCCTCTTGATCTTGTGCTACGCAACACCAGCGACCTCGCCT
AACTTTAACTGGTATCGGGAAAAGAGAGTAACGCCCCACTTCAGCCTAAACAGCCGCGATGCCGACTTTTATAAT
ACGGAAGTCACCTTCAATCCCCAGCCAGCTCACCGCGCTATTCAAGAGGCCAAAAGAGTCGGCACTATCGAAGCC
GACTTGAGTATGTGCAGACATATTGTCCTCTTTGGCAGGCATGTAAGTCCCTCTCCTACATCGAGCACCGGCCCC
TTGCTACACTTCTAACTATCTTCCTCTAGATATGGTGTGCAGGCTGCAAGAGACTCGCGAAGTGTATTGGCACGG
GATGTTTCGGACTAGCGGGCATGGGCGCTCTAATAGGTCTGATTTTTATGACAGACGGAGCTATATTGCCGATCA
TTGTAGCTTATGTTAACGGTATTGGGGGTCTCTCTATGGGCGTTGCTTCGTTTGCTGATTGCCTGGATGGCCTCT
GCCCTTGCAAACACGGCAGATGTGTGGGTGATGTACATGGTAGTGAATAA

*Pseudocercospora ulei* ERN8

ATGCGTGTCCTCGCTATCCTGTACACTATCAGTAGCCTTATGGCCTTGTGCTACGCAACACCAGCGACCTCGCCG
AACTTCGACTGGTATCGGGACAAGCAGATGACGCCAAACTTCAGACTAAACAGCCGCGATGTCGCCTTGCATCAG
ATTAAATTGGGCTTAGATGTCGAGCAGGCCTACAACACCATCCACACCGCCTACCGGGAAACCTACGAGAGCAAA
GCGGTCGCCCCATATGAAGCAGAGTTGACGCGGAGTATGTGCACCCATGTCAGTATAGCTGGCGTACATGTAAGT
CCCTCTCCTACTTCGAGCATCGCCACTTGCTGCATTCTGACTATATTCCTCCCCTAGATATGGTGTGCAGGCTGC
AAGAGACTCGCGAAGTGTATTGCCGCAGGTTGTTTCGCCCTAGCTGGCTTTGGCGGTCTTTTGGCTCTGTTTTTC
GTTACAGGATCAGTTATATTGCCTGCTATTTTGGCTTTCTTCAACGCTGTTGGGGGTTTTAGTGCAAGCGTTGCA
GTATTCAATGACTGCATGGAAGGCCTTTGTCCTTGCAAAGGCGGCAAGTGTAAGCTATAA
*Pseudocercospora ulei* GCL012
ATGCGTGTCCTCGCTATCCTGTACACTATCAGTAGCCTTATGGCCTTGTGCTACGCAACACCAGCGACCTCGCCG
AACTTCGACTGGTATCGGGACAAGCAGATGACGCCAAACTTCAGACTAAACAGCCGCGATGTCGCCTTGCATCAG
ATTAAATTGGGCTTAGATGTCGAGCAGGCCTACAACACCATCCACACCGCCTACCGGGAAACCTACGAGAGCAAA
GCGGTCGCCCCATATGAAGCAGAGTTGACGCGGAGTATGTGCACCCATGTCAGTATAGCTGGCGTACATGTAAGT
CCCTCTCCTACTTCGAGCATCGCCACTTGCTGCATTCTGACTATATTCCTCCCCTAGATATGGTGTGCAGGCTGC
AAGAGACTCGCGAAGTGTATTGCCGCAGGTTGTTTCGCCCTAGCTGGCTTTGGCGGTCTTTTGGCTCTGTTTTTC
GTTACAGGATCAGTTATATTGCCTGCTATTTTGGCTTTCTTCAACGCTGTTGGGGGTTTTAGTGCAAGCGTTGCA
GTATTCAATGACTGCATGGAAGGCCTTTGTCCTTGCAAAGGCGGCAAGTGTAAGCTATAA
*Pseudocercospora ulei* ERN8
ATGCGCCTCCTCGCCATCCTGTACACCATGAGTAGCCTCTTGACCTTGTGCTACGGAACACCAGCGACCTCGCCG
AACTTCGACTGGTATCGAGAAAAGAGAGTGACGCCCAACTTCGGCCTCAACAGCCGCGATGCCGACTTTCATGAT
GCGGATTTGACCTTCAATCCCCAGCCGGCTTACCGCGCTATTGAACGGTACAAAAGCATCGGCACTGACGACGCC
GACTGGAATATGTGCAAACATGTTTCCCTCTTTGGCTTGCATGTAAGTCCCTCTCCTACATCGAGCACCGCCCGC
GTGCTACATTTCTAATCATCTTCCTCTAGGTATGGTGTGCAGGCTGCAAGAGACTCGAGAAGTGTCTCGCCGCAG
GATTATTCGCAATAGCCGGCATGGCCGCTCTGATAGGTCTGATTTTCTTTACAGACGGAGCTATACTACCTATTA
TTAACGCTTTCCTGAACGCTGTTGGGGGTGTCGCTTTCAGCATTGCTGTGTTTGATGATTGCCTGGAAGGCCTTT
GCCCTTGCCAACACGGCGAATGTACGAATGGTGATAAACGTCATCATCATTAA
*Pseudocercospora ulei* GCL012
ATGCGCCTCCTCGCCATCCTGTACACCATGAGTAGCCTCTTGACCTTGTGCTACGGAACACCAGCGACCTCGCCG
AACTTCGACTGGTATCGAGAAAAGAGAGTGACGCCCAACTTCGGCCTCAACAGCCGCGATGCCGACTTTCATGAT
GCGGATTTGACCTTCAATCCCCAGCCGGCTTACCGCGCTATTGAACGGTACAAAAGCATCGGCACTGACGACGCC
GACTGGAATATGTGCAAACATGTTTCCCTCTTTGGCTTGCATGTAAGTCCCTCTCCTACATCGAGCACCGCCCGC
GTGCTACATTTCTAATCATCTTCCTCTAGGTATGGTGTGCAGGCTGCAAGAGACTCGAGAAGTGTCTCGCCGCAG
GATTATTCGCAATAGCCGGCATGGCCGCTCTGATAGGTCTGATTTTCTTTACAGACGGAGCTATACTACCTATTA
TTAACGCTTTCCTGAACGCTGTTGGGGGTGTCGCTTTCAGCATTGCTGTGTTTGATGATTGCCTGGAAGGCCTTT
GCCCTTGCCAACACGGCGAATGTACGAATGGTGATAAACGTCATCATCATTAA
*Pseudocercospora ulei* ERN8
ATGCGCGCCCTCGCCGTCCTGTACACCATCAGTAGCCTCTTAGCCTGGTGCTACGCAATACCAACGACCTCGCCT
AACTTTGACTGGTATCGAGAAAAGAGGGTGTCGCCTAATTTCAGCCTGAACAGCCGCGATGCCGACTACCATGAC
ATAGAAACGACCTTTAATCCCGAGCCTGCCCGTCGCGCTATTGAAGCGGCCAAAAGAGTCGGCACCTTCGAGGCC
GACATGAGTCTGTGCAGCCACGTCACCTTATTTCACAGACATGTAAGTCCCTCTCCTGCACCGAGTACCGGCCCC
TTGCTGCATTCTGACCACCTGGCTATAGATATGGTGTGCAGGCTGCAAGAGACTAGCGAAGTGTATTGGCGCAGG
ACTTTTCCAAATAGCTGGCTTTGGCGCCCTGATAGCTCTGATTTTTATTACAGATGGAGCCATCCTGCCGATTAT
TGTGGCTTGGCTCGCCGCTGTTGGGAGCGTGGCTTCAGGCATAGCCACAATCGATGAGTGCCTGGATGGCCTTTG
CCCTTGTAAACACGGGCAATGTGATTTATAA
*Pseudocercospora ulei* GCL012
ATGCGCGCCCTCGCCGTCCTGTACACCATGAGTAGTCTCTTAGCCTTGTGCTACGCAATACCAACGACCTCGCCT
AACTTTGACTGGTATCGAGAAAAGAGGGTGTCGCCTAATTTCAGCCTGAACAGCCGCGATGCCGACTACCATGAC
ATAGAAACGACCTTTAATCCCGAGCCTGCCCGTCGCGCTATTGAAGCGGCCAAAAGAGTCGGCACCTTCGAGGCC
GACATGAGTCTGTGCAGCCGCGTCACCTTCTTTCACAGACATGTAAGTCCCTCTCCTGCACCGAGTACCGGCCCC
TTGCTGCATTCTGACCACCTGGCTATAGATATGGTGTGCAGGCTGCAAGAGACTCGCGAAGTGTATTGGCGCAGG
AGTTTTCGAAATAGCTGGCTTTGGCGCCCTGATAGCTCTGATTTTCATCACAGATGGAACCATCCTGCCGATTAT
TGTGGCTTGGCTCAGCGCTGTTGGGACCGTGGCTTCAGGCGTAGCCACACTCGATGACTGCCTGGATGGCCTTTG
CCCTTGTAAACACGGGAATTGTGATGTATAA
*Pseudocercospora ulei* ERN8
ATGCGCGTCCTCGCCATCCTGTACACTATCAGTAGCCTCTTAACCTCGTGCCTCGCCTAACTTTAACTGGTATCG
GAAAAAAAGGTTGACGCCTAATTTCCGCCTAAATAGCCGCGATGTCCGACCTCCGTGAGATGGAAGAGACCTTCA
ATCCCCAACCGGCCCTACAGGCCATCGAAGCGGCTAAAAGAGTCGGCACTTTCGAAGCCGACATGAGTATGTGCA
GCCACATTACCCTCTTTGGCATACATGTA
*Pseudocercospora ulei* GCL012
AGTCGGTACTTTCGAAGCCGACATGAGTATGTGCAGCCACATTACCCTCTTTGGCATACATGTA
*Septoria petroselini* CBS 182.44

```
ATGCGTTTCTTATACACTTTTGCCGCAATCTGTTATGGATATATCGCAATTGTGCTCGCAAGCCCTACGCGCTCA
CCTAACTTCCTCTGGTACGAGGCCAAGAAGATGTCTCCCAACTTCGAGCTGCTCGCTCGGAGCATAAGCACAAAT
ACGCCGCCTCCAGTAGACGGAAAATGGTACTCGGAAATACTAGCCGTGATTGCACAAGCTAAACTCGACCCGGAG
TACGACTTCGAAAGGGAGGACGAAAACGACTCGAAATTTAAGTTGCCGAAGGTCCACTGCACTGGATTACAGATT
CAAATCTCGAAAACCAAAAAATTCGTAGGGCATCAACATCACAGCTCGATAGTAGCATGGACTGACCTCCCCATT
CTATTAGGGCTGGTGCGCAGGATGTCAGAGATTAGAAAAATGCCTCGGAGTACTTTTCTTTGTGAAAGGAGGGAT
TGCTGCTTACATCTATGCTTTCTACTTTAGCGGAGGACTCTCCGTGCCAGCAACTGTTGCCTTTCTTAACGTTGT
TGGAAGTATCGGATTTGCAGCCTCATGCTTCAACACTTGTGTAGATGGCTTCTGCGCAACCAAAAGCGGAAACTA
TCATATTTGCGGTGATTAA
```

References for proteomes and genomes from which Avr9B-like protein and *Avr9B*-like gene sequences, respectively, were retrieved:


**Aggarwal R, Sharma S, Singh K, Gurjar MS, Saharan MS, Gupta S, Bashyal BM, Gaikwad K. 2019.** First draft genome sequence of wheat spot blotch pathogen *Bipolaris sorokiniana* BS_112 from India, obtained using hybrid assembly. *Microbiology Resource Announcements* **8**: e00308-19.

**Akinsanmi OA, Carvalhais LC. 2020.** Draft genome of the Macadamia husk spot pathogen, *Pseudocercospora macadamiae*. *Phytopathology* **110**: 1503–1506.

**Cao Y, Cai Q, Li C, Song G, Lu N, Yang Z. 2023.** Genome sequence resource of *Curvularia clavata* causing leaf spot disease on tobacco by Oxford Nanopore PromethION. *Plant Disease* PDIS-09.

**Chand R, Pa C, Singh V, Kumar M, Singh VK, Chowdappa P. 2015.** Draft genome sequence of *Cercospora canescens*: a leaf spot causing pathogen. *Current Science* **109**: 2103–2110.

**Chang TC, Salvucci A, Crous PW, Stergiopoulos I. 2016.** Comparative genomics of the Sigatoka disease complex on banana suggests a link between parallel evolutionary changes in *Pseudocercospora fijiensis* and *Pseudocercospora eumusae* and increased virulence on the banana host. *PLoS Genetics* **12**: e1005904.

**Condon BJ, Leng Y, Wu D, Bushley KE, Ohm RA, Otillar R, Martin J, Schackwitz W, Grimwood J, MohdZainudin N et al. 2013.** Comparative genome structure, secondary metabolite, and effector coding capacity across *Cochliobolus* pathogens. *PLoS Genetics* **9**: e1003233.

**de Siqueira KA, Senabio JA, Pietro-Souza W, de Oliveira Mendes TA, Soares MA. 2021.** *Aspergillus* sp. A31 and *Curvularia geniculata* P1 mitigate mercury toxicity to *Oryza sativa* L. *Archives of Microbiology* **203**: 5345–5361.

**de Wit PJGM, van der Burgt A, Ökmen B, Stergiopoulos I, Abd-Elsalam KA, Aerts AL, Bahkali AH, Beenen HG, Chettri P, Cox MP et al. 2012.** The genomes of the fungal plant pathogens *Cladosporium fulvum* and *Dothistroma septosporum* reveal adaptation to different hosts and lifestyles but also signatures of common ancestry. *PLoS Genetics* **8**: e1003088.

**Franco MEE, López S, Medina R, Saparrat MCN, Balatti P. 2015.** Draft genome sequence and gene annotation of *Stemphylium lycopersici* strain CIDEFI-216. *Genome Announcements* **3**: e01069-15.

**Gao S, Li Y, Gao J, Suo Y, Fu K, Li Y, Chen J. 2014.** Genome sequence and virulence variation-related transcriptome profiles of *Curvularia lunata*, an important maize pathogenic fungus. *BMC Genomics* **15**: 1–18.

**Gazzetti K, Diaconu EL, Nanni IM, Ciriani A, Collina M. 2019.** Genome sequence resource for *Stemphylium vesicarium*, causing brown spot disease of pear. *Molecular Plant–Microbe Interactions* **32**: 935–938.

**González-Sayer S, Oggenfuss U, García I, Aristizabal F, Croll D, Riaño-Pachon DM. 2022.** High-quality genome assembly of *Pseudocercospora ulei* the main threat to natural rubber trees. *Genetics and Molecular Biology* **45**: e50510051.

**He K, Zhao C, Zhang M, Li J, Zhang Q, Wu X, Wei S, Wang Y, Chen X, Li C. 2023.** The chromosome-scale genomes of *Exserohilum rostratum* and *Bipolaris zeicola* pathogenic fungi causing rice spikelet rot disease. *Journal of Fungi* **9**: 177.
**Kuan CS, Yew SM, Toh YF, Chan CL, Ngeow YF, Lee KW, Na SL, Yee WY, Hoh C-C, Ng KP. 2015.** Dissecting the fungal biology of *Bipolaris papendorfii*: from phylogenetic to comparative genomic analysis. *DNA Research* **22**: 219–232.
**Litvintseva AP, Hurst S, Gade L, Frace MA, Hilsabeck R, Schupp JM, Gillece JD, Roe C, Smith D, Keim P, Lockhart SR. 2014.** Whole-genome analysis of *Exserohilum rostratum* from an outbreak of fungal meningitis and other infections. *Journal of Clinical Microbiology* **52**: 3216–3222.
**Luo X, Cao J, Huang J, Wang Z, Guo Z, Chen Y, Ma S, Liu J. 2018.** Genome sequencing and comparative genomics reveal the potential pathogenic mechanism of *Cercospora sojina* Hara on soybean. *DNA Research* **25**: 25–37.
**Ma Q, Cheng C, Geng Y, Zang R, Guo Y, Yan L, Xu C, Zhang M, Wu H. 2023.** The draft genome sequence and characterization of *Exserohilum rostratum*, a new causal agent of maize leaf spot disease in Chinese Mainland. *European Journal of Plant Pathology* **165**: 57–71.
**McDonald MC, Taranto AP, Hill E, Schwessinger B, Liu Z, Simpfendorfer S, Milgate A, Solomon PS. 2019.** Transposon-mediated horizontal transfer of the host-specific virulence protein ToxA between three fungal wheat pathogens. *MBio* **10**: e01515-19.
**Medina R, Franco MEE, Lucentini G, Saparrat MCN, Balatti PA. 2018.** Draft genome sequences of sporulating (CIDEFI-213) and nonsporulating (CIDEFI-212) strains of *Stemphylium lycopersici*. *Microbiology Resource Announcements* **7**: e00960-18.
**Searight J, Famoso AN, Zhou XG, Doyle VP, Richards JK. 2023.** A high-quality genome assembly for *Cercospora janseana*, causal agent of narrow brown leaf spot of rice. *Molecular Plant-Microbe Interactions* https://doi.org/10.1094/MPMI-10-22-0222-A.
**Sharma S, Hay FS, Pethybridge SJ. 2020.** Genome resource for two *Stemphylium vesicarium* isolates causing Stemphylium leaf blight of onion in New York. *Molecular Plant–Microbe Interactions* **33**: 562–564.
**Wingfield BD, de Vos L, Wilson AM, Duong TA, Vaghefi N, Botes A, Kharwar RN, Chand R, Poudel B, Aliyu H et al. 2022**. IMA Genome-F16: Draft genome assemblies of *Fusarium marasasianum*, *Huntiella abstrusa*, two *Immersiporthe knoxdaviesiana* isolates, *Macrophomina pseudophaseolina*, *Macrophomina phaseolina*, *Naganishia randhawae*, and *Pseudocercospora cruenta*. *IMA Fungus* **13**: 3.
**Zaccaron AZ, Bluhm BH. 2017.** The genome sequence of *Bipolaris cookei* reveals mechanisms of pathogenesis underlying target leaf spot of sorghum. *Scientific Reports* **7**: 1–15.
**Zaccaron AZ, Stergiopoulos I. 2020.** First draft genome resource for the tomato black leaf mold pathogen *Pseudocercospora fuligena*. *Molecular Plant–Microbe Interactions* **33**: 1441–1445.
**Zeng F, Wang C, Zhang G, Wei J, Bradley CA, Ming R. 2017.** Draft genome sequence of *Cercospora sojina* isolate S9, a fungus causing frogeye leaf spot (FLS) disease of soybean. *Genomics Data* **12**: 79–80.
**Zhang W, Yang Q, Yang L, Li H, Zhou W, Meng J, Hu Y, Wang L, Kang R, Li H, Ding S, Li G. 2023.** High-quality nuclear genome and mitogenome of *Bipolaris sorokiniana* strain LK93, a devastating pathogen causing wheat root rot. *Molecular Plant-Microbe Interactions* https://doi.org/10.1094/MPMI-09-22-0196-A.
**Zhao P, Crous PW, Hou LW, Duan WJ, Cai L, Ma ZY, Liu F. 2021.** Fungi of quarantine concern for China I: Dothideomycetes. *Persoonia-Molecular Phylogeny and Evolution of Fungi* **47**: 45–105.
*A. gansuensis* LYZ1412: https://www.ncbi.nlm.nih.gov/nuccore/WBOK00000000.1

*Alternaria* sp. section Undifilum W740-01: https://www.ncbi.nlm.nih.gov/nuccore/2274091898
*B. bicolor* ML9021: https://www.ncbi.nlm.nih.gov/nuccore/JAODYD000000000.1
*B. oryzae* Bo-Gvt: https://www.ncbi.nlm.nih.gov/sra/SRX8380151
*B. sorokiniana* ND90Pr: https://www.ncbi.nlm.nih.gov/nuccore/AEIN00000000.1
*B. sorokiniana* Pusa2: https://www.ncbi.nlm.nih.gov/sra/SRX7473635
*Bipolaris* sp. ADL-507: https://www.ncbi.nlm.nih.gov/sra/SRX244593
*C. sojina* 2.2.3: https://www.ncbi.nlm.nih.gov/nuccore/1466071976
*C. sojina* CCC: https://www.ncbi.nlm.nih.gov/biosample/SAMN19354503
*C. sojina* RACE15: https://www.ncbi.nlm.nih.gov/biosample/SAMN10531911
*C. geniculata* W3: https://www.ncbi.nlm.nih.gov/nuccore/1356663259
*C. hawaiiensis* PK9021: https://www.ncbi.nlm.nih.gov/nuccore/JAODYC000000000.1
*C. lunata* Cl-Gvt: https://www.ncbi.nlm.nih.gov/sra/SRX8380154
*C. lunata* W3: https://www.ncbi.nlm.nih.gov/nuccore/1633470750
*Curvularia* sp. ZM96: https://www.ncbi.nlm.nih.gov/nuccore/2202949488
*C. spicifera* FR9030: https://www.ncbi.nlm.nih.gov/nuccore/JAODYB000000000.1
*P. ulei* ERN8: https://mycocosm.jgi.doe.gov/Pseule1/Pseule1.home.html
*P. pini-densiflorae* CBS 125139: https://www.ncbi.nlm.nih.gov/nuccore/AWYD00000000.2
*E. rostratum* BF9006: https://www.ncbi.nlm.nih.gov/nuccore/JANXEY000000000.1
*E. rostratum* ER1: https://www.ncbi.nlm.nih.gov/nuccore/2076348907
*E. rostratum* M 18 0203: https://www.ncbi.nlm.nih.gov/sra/ERX3211936
*E. rostratum* SR-KPL1: https://www.ncbi.nlm.nih.gov/nuccore/JABMLL000000000.1
