## Supplementary material for "Sequential breakdown of the complex *Cf-9* leaf mould resistance locus in tomato by *Fulvia fulva*": Table S1 and Table S4

**Table S1.** Statistics for the *Fulvia fulva* strain IPO 2679 Illumina genome assembly.

| Statistic | Value |
|---|---|
| Number of assembled scaffolds | 13,307 |
| N50 value[1] | 36 kb |
| BUSCO score[2] | ~97% |
| Coverage | 106x |

[1]N50 value: where 50% of the assembly is contained in contigs equal to or larger than the value.
[2]Benchmarking Universal Single-Copy Orthologs (BUSCO) score: percentage of highly conserved genes in the assembly.

**Table S4.** Primers used in this study.

| Primer name | Sequence (5′–3′) | Reference |
|---|---|---|
| **Gene presence/absence screen (polymerase chain reaction, PCR) and allelic variation analysis (PCR amplicon sequencing)** | | |
| Avr9-F | AGTAGATCCGGCCGAGAGAG | Iida *et al.* (2015) |
| Avr9-R | AAAGCCTTCAATATGAACGAAT | Iida *et al.* (2015) |
| Avr9B-F | CATATATAAACTCCTCGCTCGCCCTC | This study |
| Avr9B-R | CACCGAGCACTATCATTTCACATGC | This study |
| **Gene expression analysis (real-time quantitative PCR, RT-qPCR) and copy number assessment (quantitative PCR, qPCR)** | | |
| qCfActin-F | GGCACCAATCAACCCAAAG | Mesarich *et al.* (2014) |
| qCfActin-R | TACGACCAGAAGCGTACAG | Mesarich *et al.* (2014) |
| qAvr9B-F | GATTTGCGTCACCGTTCTCG | This study |
| qAvr9B-R | CCGATTGAGAGCAAACAGGC | This study |
| qECP5-F | TACGACACGACTGGAGAAC | This study |
| qECP5-R | CGAACATCAAACGTCAAATGC | This study |
| ***Agrobacterium tumefaciens*-mediated transient transformation assays (ATTAs)** | | |
| pICH86988-F | AGGACACGCTCGAGTATAAG | This study |
| pICH86988-R | CTCTTCGGATACTAGCGTAC | This study |
| CfCE54_BsaI-F | GGTCTCGAATGATGGGATTTG | This study |
| CfCE54_BsaI-R | GGTCTCAAAGCCTAACACAAC | This study |
| NSPCfCE54_BsaI-F | GGTCTCGAATGGACTACAAGG | This study |
| W97C-F | CGGTTGCTGTGCGGGTT | This study |
| W97C-R | TACAACCCGCACAGCAACCG | This study |
| **Gene complementation experiment** | | |
| pFBTS1-F | GCACGTGAGAACGCTAATAGCCCTTTC AGATCAACAGCTT | This study |

| | | |
|---|---|---|
| pFBTS1-R | CTATTAGCGTTCTCACGTGC | This study |
| pFBTS1BB-F | TTGATATCGAATTCCTGCAGC | This study |
| pFBTS1BB-R | AGCTTGATATCTGTTAGTAATC | This study |
| GibEcp5-F | GATTACTAACAGATATCAAGCTGATTC AGCTTCTCGCTATAG | This study |
| GibEcp5-R | GCTGCAGGAATTCGATATCAACTCTAC ACGGGCGCGACTGC | This study |
| GibCfCE54-F | GATTACTAACAGATATCAAGCTCCTAT CGTAAAGCGCCTGAG | This study |
| GibCfCE54-R | GCTGCAGGAATTCGATATCAAGATGGA TCCCTACTTCGCCTTC | This study |
| Comp-F | TGTTATTGCCAGTGCCGCTTCAATTCAT GATG | This study |
| CompEcp5-R | CGCCACTAGTCTCGAGTTAATTAACCG | This study |
| CompCfCE54-R | GCTGCAGGAATTCGATATCAAGATGGA TCCCTACTTCGCCTTC | This study |
| **PCR analysis of deleted genome region containing *Avr9B*** | | |
| Avr9B_deletion-F_1 | GGAGCGTGCTTAAATGAGCG | This study |
| Avr9B_deletion-R_1 | GTCCCGAGATCTAGGACGAGT | This study |
| Avr9B_deletion-F_2 | ACACAACACGCGAAACACAG | This study |
| Avr9B_deletion-R_2 | GTCCCGAGATCTAGGACGAGTA | This study |
